# The structure of a thermostable phage’s portal vertex and neck complex illuminates the headful maturation mechanism

**DOI:** 10.1101/2025.04.14.648768

**Authors:** Emma L. Sedivy, Emily Agnello, Julia E. Hobaugh, Rakeyah Ahsan, Kangkang Song, Chen Xu, Brian A. Kelch

**Affiliations:** Department of Biochemistry and Molecular Biotechnology, University of Massachusetts Chan Medical School, Worcester MA

## Abstract

Viruses assemble from component parts inside their host cells, but the mechanisms coordinating this complex process are not completely understood. In tailed bacteriophages, the genome is packaged into its capsid shell through the portal complex. The portal complex then closes to retain DNA and connects to the tail, which is required for host recognition and infection. The trigger to stop pumping DNA and assemble the mature virus has been a longstanding conundrum in the field. We determined the structure of the portal, the proteins that connect it to the tail, and portal vertex in the hyperthermophilic phage Oshimavirus using cryo-Electron Microscopy (cryo-EM). We find highly intertwined loop structures, like in a wicker basket, stabilizing the portal vertex against high temperatures. Moreover, we observe that the portal protrudes from the capsid in mature virions. We propose that portal is repositioned by packaged DNA, forming a pressure-sensitive switch that terminates genome packaging and triggers tail attachment in headful phages.

## INTRODUCTION

Bacteriophages (phages), viruses that infect bacteria, are the most abundant biological entities on Earth (1). Phages abound in every environment inhabited by bacteria, from human microbiomes to scalding geothermal vents to open oceans (2). They have vital impacts on our lives because they shape bacterial populations as vectors of horizontal gene transfer, drivers of microbial evolution and population dynamics, and bactericidal agents (3). Since their discovery in 1915, phages have been used as a model to understand the fundamental rules of molecular biology and as a source of therapeutic agents and molecular tools (4, 5). Nevertheless, the mechanisms powering many of their intricate components are still poorly understood. Closing these gaps in our understanding will enable biotechnology to take greater advantage of the opportunities that phages offer.

Members of the *Caudovirales* order, the majority of isolated phages, encapsulate their double-stranded DNA genome inside an icosahedral or prolate capsid and use a tail for host recognition and as a conduit for DNA passage into the cell (2). The neck complex attaches the filled capsid to the tail. The neck joins the capsid at a 5-fold symmetric vertex, where a single pentamer of capsid proteins is replaced with the homo-dodecameric portal complex, which provides access to the interior of the capsid and nucleates assembly of the initial, smaller procapsid form of the capsid (2, 6, 7). The DNA packaging motor, terminase, binds to the portal vertex and portal provides a conduit for DNA to be packaged by terminase into the capsid (8–11). During packaging, the procapsid also expands to form a mature capsid in the shape of an icosahedron with the portal at one vertex (12, 13). Double-stranded DNA phages generally replicate their genomes as concatemers that are subsequently cleaved by terminase after a full genome is packaged into the capsid. There are several strategies the packaging motor can use to recognize the appropriate point at which to cleave the phage DNA. Some terminases recognize a fixed sequence or position in the genome at which to cleave, cutting at the same position in each phage. In contrast, in headful phages, terminase cuts when the capsid is full, cutting at a slightly different position each time and generating phages with circularly permuted genomes (12). It is not known how headful phages signal to terminase that the capsid is full and DNA should be cleaved, although it is clear that portal plays a major role in the headful signal (14). After packaging is complete, terminase dissociates and portal changes conformation to retain packaged DNA (8, 15). It then binds to the neck complex and joins the icosahedral capsid head to the tubular tail. Because the capsid and tail have different symmetries, the neck must accommodate the mismatch between these components (8). The genome must be retained inside the capsid until the host is found, as premature genome release renders the virion inert. However, the packaged genome generates extremely high pressure inside the capsid, which must be contained by both the capsid and the neck (16, 17). Attachment to the host triggers irreversible rearrangements in the virion and drives genome ejection through the neck and the tail into the host, where the phage can replicate (18, 19).

Phages following this basic architecture and life cycle make up most known viruses and are found in every environment, including the human microbiome (2). Furthermore, capsid and portal components are conserved among human pathogens in the Herpesvirus order, including those that cause chickenpox, shingles, infectious mononucleosis, and Kaposi’s sarcoma (20, 21). In contrast to the prodigious diversity of phages, phage biology has historically focused on a handful of model phages that infect lab strains of bacteria (2). Examining a greater diversity of phages, especially those that inhabit extreme niches, may help to uncover general principles of phage biology. The *Oshimavirus* genus, comprising nearly identical isolates P74-26, P23-45, and G20C, infect the bacterium Thermus thermophilus HB8 in geothermal hot springs at 65-75 °C and have the longest tails of all known phages (22, 23). *Oshimaviruses* have recently been developed as a model system for revealing basic phage assembly mechanisms because of the high thermostability and novel mechanisms these phages utilize to overcome the challenges of their extreme environment (15, 24–31). We and others described features of the P74-26 capsid and tail tube proteins that enhance their stability (28, 29, 31), and others described the procapsid portal vertex of P23-45 (15).

Their extreme habitat intensifies a problem faced by all viruses: protecting the genome until the appropriate time to release it. The challenge of retaining the genome is even greater at elevated temperatures, and the portal vertex creates a unique vulnerability in the capsid shell. We wondered how a long-tailed thermophilic phage makes a durable portal vertex and neck. Furthermore, it remains unknown how the portal vertex rearranges during the viral maturation process. We can now reveal these mechanisms by determining the structure of the neck in the mature virion and comparing it with the structures of the portal vertex in the procapsid and empty mature capsid states (15, 28).

We report the structure of the Oshimavirus P74-26 neck at high resolution using cryo-EM and identify its constituent proteins. Analysis of this structure provides insight into mechanisms of thermal stability of large complexes and how pressure regulates the maturation of headful bacteriophages.

## RESULTS

### The Oshimavirus neck includes regions of C12, C6, C5, and C3 symmetry

We determined the structures of the Oshimavirus neck by cryo-EM. The organization of the neck was not known, but the Oshimavirus tail tube has 3-fold symmetry (31) and the portal complex is dodecameric (15, 28). The capsid vertex in which the neck is located has 5-fold symmetry (8) (Fig. 1A). Known components of the neck and portal vertex include the portal, Decoration Protein (Dec), Major Capsid Protein (MCP), and Tail Tube Protein (TTP), which are located near each other in Oshimavirus genomes along with many unannotated open reading frames that may include neck proteins (Fig. 1B) (23, 29).

**Figure 1:**
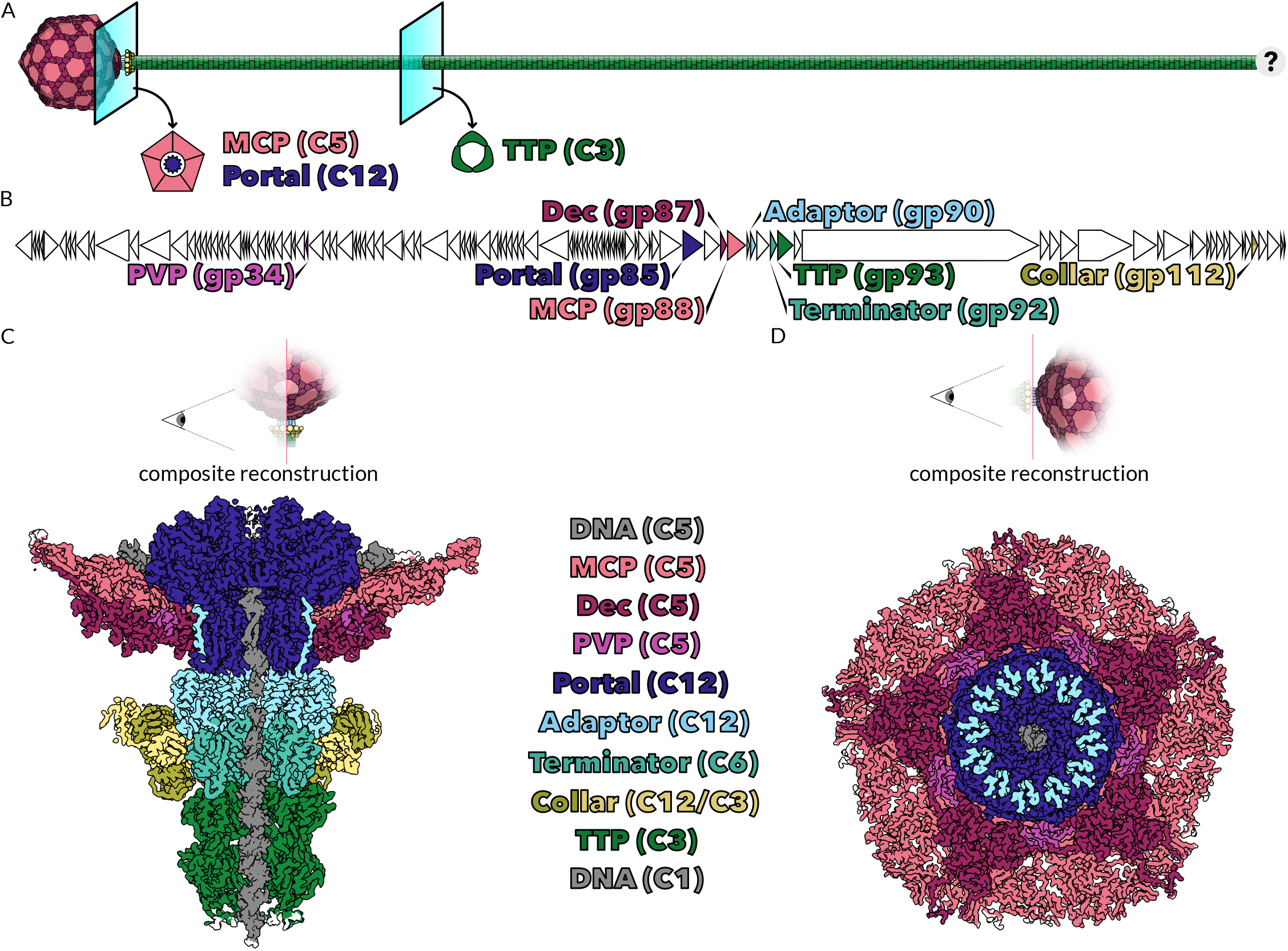
Reconstruction of Oshimavirus P74-26 Neck components revealed proteins not previously annotated. **A. Diagram of the Oshimavirus virion**. Cross-sections show the symmetry of previously studied phage components. **B. Linear map of the Oshimavirus P74-26 genome, with the location of structural genes highlighted**. Each ORF in the 82kb genome is displayed as a triangle proportional to its length. Genes are colored according to the key in C. **C. Cross-section of a composite reconstruction of the Oshimavirus P74-26 Neck complex**. The C5, C3, and central DNA maps of the Neck complex were aligned by fitting into the asymmetric map and are colored according to protein identity. The symmetry of each component is noted in parentheses. C5 DNA is found inside the capsid, wrapping around Portal, while C1 DNA is found in the central lumen of the neck. The Portal Vertex Protein (PVP), Adaptor, Terminator, and Collar were not previously annotated. **D. View of the portal vertex from outside the capsid**. The composite reconstruction in C was rotated 90° and sliced to show the portal vertex.

We first reconstructed the Oshimavirus neck with 12-fold symmetry imposed (Fig. S1-3). There was significant variation in local resolution, and regions that exhibit other symmetries (such as capsid, packaged genome, and tail tube) were blurry, as expected (Fig. S5A). This reconstruction reached 2.3-Å Gold Standard Fourier Shell Correlation (GSFSC) resolution with a mask excluding off-symmetry regions of the map (Fig. S5A). The neck forms a hollow tube with a central channel ∼45 Å in diameter, similar to the lumen of the tail tube (31) and connects to density consistent with the mushroom-shaped portal that sits in the aperture of the capsid opening. In Oshimavirus, the portal is ∼120 Å diameter at its widest point (15, 28). We observed that the portal is connected to the tail tube by a neck that is ∼80 Å in length and surrounded by a ring with 12 lobes protruding to 200 Å wide. We termed this ring around the neck the collar region. The lobes of the collar did not achieve high resolution with 12-fold symmetry imposed. To separate virions with conformational or compositional heterogeneity in this region, we 3D classified the particle set with a mask applied to the region (Fig. S3). We reconstructed two classes, one lacking the lobes entirely and another with a complete set of lobes in the collar region. The resolution of the lobes was still weak relative to the rest of the neck (Fig. 5A). Because there were no other structural changes between necks with and without the collar lobes, we concluded that they are not required for structural integrity of the virion but may be required or beneficial for the phage during infection (Fig. S5B).

The lumen of phage necks often contains DNA and/or the N-terminal region of TMP (32, 33). The C12 reconstruction of the neck showed amorphous density running through the central channel. To further examine this density, we performed symmetry expansion around the C12 space group upon the particle set of complete necks. We then performed focused 3D classification with a cylindrical mask covering the neck lumen below the portal pore loops (Fig. S4). Many of the resulting classes showed clear density for B-form double stranded DNA running through the entire lumen, while the other classes remained amorphous. We selected the clearest class for further refinement and the resulting map had strong (∼3-Å resolution) density for the neck itself and 4.5-Å resolution for the central channel (Fig. S5C). dsDNA travels through at least 2 rings of the tail tube, ∼66 base pairs beyond the portal pore loops, without any density for TMP.

To visualize the region of the neck that contacts the C3 tail tube, we performed symmetry expansion of the particle set around the C12 symmetry group followed by 3D classification with a mask applied to the collar region (Fig. S6). Duplicate particles were removed from the largest class, which was then refined with 3-fold symmetry to 2.6-Å GSFSC resolution (Fig. S8A). With 3-fold symmetry imposed, it became clear that while the core of the collar has 12-fold symmetry, the region that interacts with the first ring of the tail tube has 3-fold symmetry (Fig. S6, S19). In addition, some of the lobes of the collar region may be flexible and/or unstructured, explaining their poor resolution (Fig. S19). We also applied this workflow to a stack of particles that were selected because they resembled the C12-refined neck without density for the capsid (Fig. S7). This particle set yielded a 2.7-Å resolution reconstruction that was identical to the complete neck map generated in Fig. S6, except for the absence of the capsid itself (Fig. S8B). We hypothesize that these particles were formed by removal or fragmentation of the capsid and conclude that the survival of the neck highlights its durability.

To visualize how the capsid accommodates the aperture in which the portal sits, we refined the same particle set with 5-fold symmetry imposed to 2.9-Å GSFSC resolution (Fig. 1D, S9-10). In addition to the capsid vertex, we also resolved a pseudo-circle of DNA that adopts apparent 5-fold symmetry around the portal. To visualize how the C5 capsid interacts with the C12 portal, we performed symmetry expansion of the particle set around the C5 symmetry group followed by 3D classification with a mask applied to the neck (Fig. S11-12). Duplicate particles were removed from the largest resulting class, and it was refined asymmetrically. This reconstruction reached 3.5-Å GSFSC resolution (Fig. S12).

### Oshimavirus neck proteins have structural similarity to neck proteins of other phages

Other than portal, Dec, MCP, and TTP, we did not know the identities of the proteins that compose the neck region. Oshimavirus has low sequence similarity to other phages, and the genome does not contain any hypothetical proteins with sequence similarity to known neck proteins. To identify the proteins in our reconstructions, we leveraged the automated cryo-EM atomic model-building software ModelAngelo (34). After performing unbiased model-building (without sequence input), we used the NCBI-BLAST server to search for sequence similarity between built peptides and all known proteins in the non-redundant RefSeq database (Table S1). Using this approach, we were able to identify all neck proteins in our reconstructions. In the C3 map, ModelAngelo built peptides corresponding to the portal Protein and Tail Tube Protein (TTP), as expected, as well as gp90, gp92, and gp112 of Oshimavirus P74-26, previously annotated as hypothetical proteins (Fig. 1B, C). Into the C5 map, ModelAngelo built peptides corresponding to the Major Capsid Protein (MCP) and Decoration Protein (Dec), as expected, as well as hypothetical protein gp34 (Fig. 1B,D). To confirm our assignments, we used mass spectrometry to identify the proteins that are present in the mature virions (Table S2). All putative neck proteins are present in the mature virion samples as measured by mass spectrometry.

We then performed ModelAngelo model-building in the C3 and C5 maps, this time including the consensus sequences of identified Oshimavirus P74-26 proteins as input. We also generated a segment of ideal B-form DNA and fit it into the central channel DNA density in the C1 map. These models were manually adjusted and refined (Table S4). We used the DALI structural comparison server to search for proteins with similarity to these models. We found that gp34, which we named Portal Vertex Protein (PVP), is similar only to the gp2 RNA Polymerase inhibitor of phage T7 (35) and an auxiliary capsid protein of phage ΦcrAss001 (Fig. S15A, S16) (36). We found that gp90 is structurally similar to adaptor proteins present in the neck of a wide variety of tailed phages, and therefore named it adaptor protein (Fig. S15B, S17) (37). We found that gp92 closely resembles the phage Lambda tail terminator protein gpU and therefore named it terminator protein (Fig. S15C) (38, 39). gp112, which we named collar protein, consists of two Immunoglobulin (Ig) domains, common in phage necks and Tails (40), connected by a flexible linker (Fig. S15D, E, S16). The ModelAngelo results, mass spectrometry data, and structural similarity to known neck components give us high confidence in our protein assignments.

The portal vertex of the capsid is bordered by 5 hexameric capsomers of MCP, with trimers of Dec at the junctions between capsomers. 5 subunits of PVP also border the aperture, bridging Dec trimers (Fig. 1D). The portal complex sits inside this aperture. A dodecamer of adaptor forms a ring that joins the portal dodecamer to the hexameric ring of terminator, which in turn connects to the first trimeric ring of TTP in the tail tube. The 12 lobes of the collar region encircle the terminator ring, each formed by a dimer of collar proteins. Within the collar dimer, the two N-terminal domains interact with each other, leaving the two C-terminal domains free (Fig. S19). One of these C-terminal domains forms the distal part of each lobe in the upper collar, extending perpendicularly from the main tube of the neck. The other C-terminal domain of each dimer extends towards the first ring of TTP, although only 6 of them can be resolved. The resolved 6 C-terminal domains form the lower collar and interact with TTP in 3 pairs arranged with 3-fold symmetry around the ring of TTP, which also has 3-fold symmetry. Although the remaining 6 C-terminal domains are not clearly resolved, when the contour threshold is relaxed, 3 areas of density appear in the gaps where C-terminal domains are missing. This density likely represents the missing domains of the lower collar, which have been displaced by steric clashes with TTP and are either flexibly tethered or unstructured (Fig. S19).

### Portal is in the closed conformation

The portal complex plays vital roles in the necks of tailed phages and herpesviruses, providing a conduit for DNA to be translocated into the capsid. Oshimavirus portal protein, like other portals, is dodecameric and forms a mushroom-shaped complex with a central pore. The crown domain forms the top of the mushroom, with the Wing domain forming the outer rim of the mushroom cap and the clip domain forming the stem (Fig. 2A). The portal has been proposed to act as a gatekeeper during DNA packaging and genome ejection, with its central pore changing conformation to either allow passage of DNA or to retain it in the capsid (15, 41, 42). Previous studies of closely related isolates P23-45 and G20c have reported structures of the portal complex in procapsids and in crystals (PDB 6QJT and 6IBG, respectively). The authors hypothesized that the procapsid portal is in an open conformation, allowing passage of DNA through the pore, while the crystallized form of portal represents the conformation in mature virions that retains DNA inside the capsid (15, 28). On the other hand, studies of the T7 podophage suggest that the portal is in the open conformation and that the nozzle protein at the tip of the tail is the gate that retains DNA (43). Thus, it is unclear whether portal is the gatekeeper that retains DNA in long-tailed thermophilic phage.

**Figure 2:**
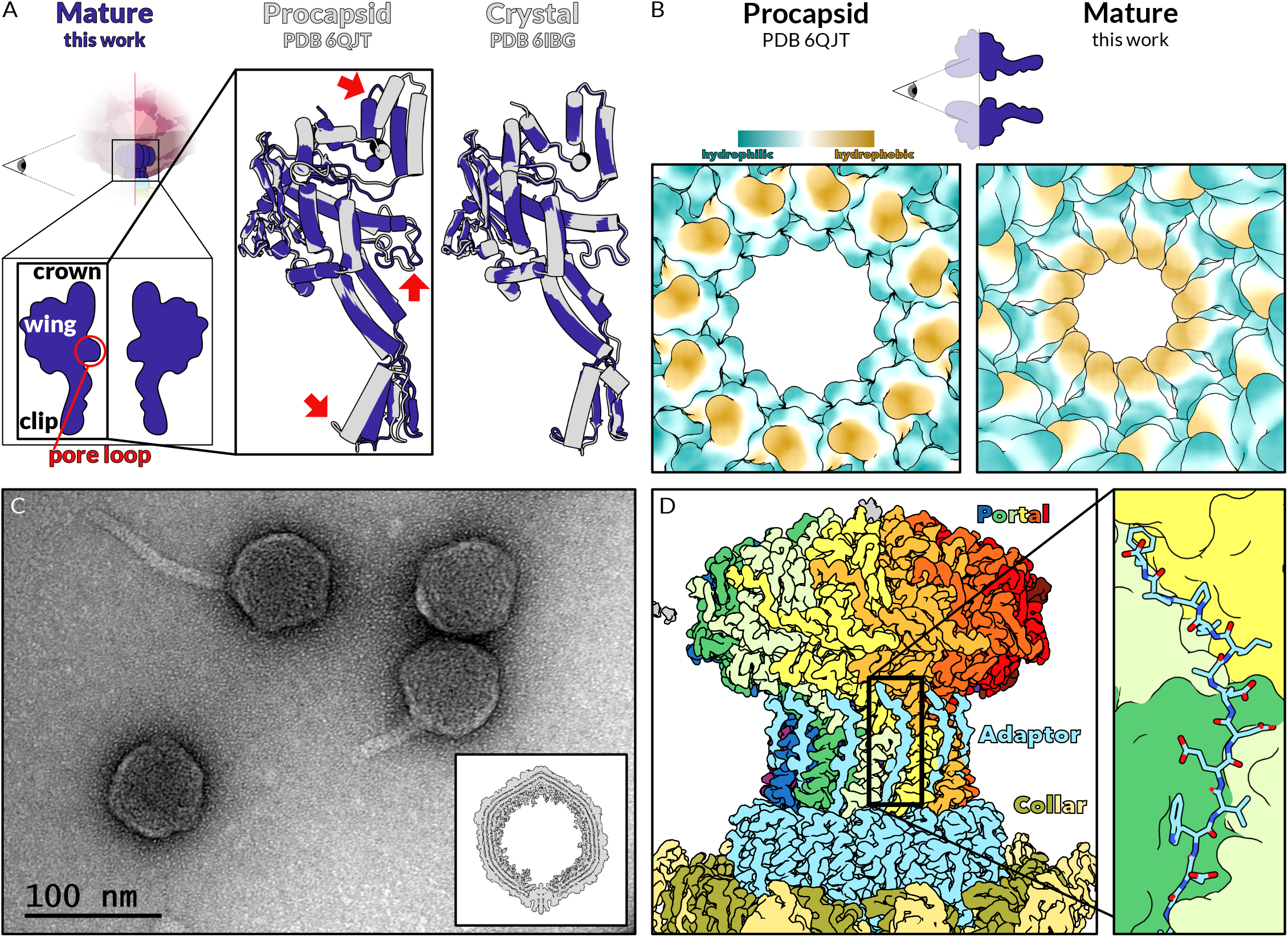
Portal protein is in the closed conformation and forms strong interactions with the C-terminal tail of adaptor. **A. Comparison of mature Portal protein (dark blue) with subunits of the portal protein crystal and procapsid portal protein (grey)**. Significant domains of the portal protein are noted in the cartoon diagrams, left. Regions with significant differences (clip, crown, and pore loop) are noted with red arrows. **B. Comparison of the portal pore loop constriction in procapsids and mature virions**. Surface view of the models of the procapsid (open) and mature (closed) Portals colored according to hydrophobicity. **C. Virions with broken tails imaged by negative stain EM**. A band migrating below mature virions in a Cesium Chloride gradient was purified and stained with Uranyl Acetate. *Inset:* We visualized the same sample by cryoEM and reconstructed filled capsids. 3D classicfication revealed a subset of particles consist of filled capsids with only a portal complex and no other components of neck or tail. **D. EM density of the neck complex with portal in rainbow**. Adaptor and collar proteins are colored as in Figure 1. Inset shows the atomic model of the adaptor tail (sticks) interacting with the clip region of three different portal subunits (surfaces).

Supporting portal’s role as a gatekeeper, the conformation that we observe in mature Oshimavirus virions matches the closed form of the crystal structure (Ca RMSD = 1.028 Å; Fig. 2A). In contrast, the portal conformation in mature virions is inconsistent with that found in procapsids (PDB 6QJT), with substantial differences in the crown domain and the pore loops and smaller alterations in the clip domain (Fig. 2A). As reported by Bayfield et al., the procapsid form of portal has a wider and more hydrophilic pore, while the crystal and mature portal’s pore is narrower and more hydrophobic (Fig. 2B). We were surprised to find that none of the other neck proteins formed constrictions in the neck pore that could contribute to genome retention, as have been found in other similar phages (Fig. S23) (32, 44). Furthermore, we fortuitously isolated a form of Oshimavirus from a band that migrates lower than intact virions in a Cesium Chloride gradient in which the tail is broken along the length of the tail (Fig. S13). Negative stain EM and Cryo-EM imaging of these particles clearly shows DNA retained in the head, even though the tail tip complex is missing (Fig. 2D, S13). We performed asymmetric reconstructions of these phage capsids, which showed clear density for packaged DNA (Fig. S14). 3D classification with a mask focused around the unique vertex revealed that a subset of these capsids contained an intact portal complex, but lacked any neck or tail components (Fig. 2C, S14). These particles indicate that the portal complex is the main gate for retaining DNA inside the capsid, and support the hypothesis that hydrophobic residues V325 and I330 are responsible for gating DNA (15). It remains unknown how the portal transitions from an open conformation to a closed conformation, and what provides the signal to trigger this change.

### Consecutive rings transition from 12-fold symmetry of portal to the 3-fold symmetry of the tail tube

The adaptor protein forms a dodecameric ring and uses its C-terminal tail to attach to the portal clip region (Fig. 2D). Adaptor tails bind in the clefts formed in the clip domain where adjacent portal subunits meet. Each adaptor tail contacts 3 subunits of portal (Fig. 2D). The interface between the adaptor tail and the portal clip exhibits complementary hydrophobic (Fig. S20B) and electrostatic interactions (Fig. S20C). For example, the C-terminal residue (Phe130) inserts its sidechain into a hydrophobic pocket between the clip and wing domains, while the C-terminal carboxylate moiety forms electrostatic interactions with basic residues of the clip domain. Thus, the adaptor interaction appears strong, which makes sense given its vital role in anchoring the 0.9 µm tail onto the capsid.

The hexameric terminator bridges the 3-fold symmetry of the tail tube to the 12-fold symmetry of adaptor and portal. Two adaptor subunits interact with the “top” loops of one terminator subunit through electrostatic and hydrophobic interactions (Fig S21). Meanwhile, a long loop at the “bottom” of each terminator subunit binds to the sockets of the 3-fold symmetric TTP ring. This arrangement mimics the loop-and-socket geometry that TTP rings use to stack into the 0.9 µm long tail tube (Fig. S22) (31). Each TTP subunit contains two sockets that both need to be occupied, for a total of 6 sockets per TTP ring. Each of the 6 terminator subunits contains a single loop, which can insert into either of the two TTP sockets. In contrast, in a typical stack of rings of the tail tube, the two loops from a TTP in the ring above insert into the sockets of a TTP subunit of the ring below (31). The loops of TTP are each specific to one socket, with Loop1 binding to Socket1 and Loop2 binding to Socket2. In contrast, the loop of terminator is bivalent and occupies either socket of TTP. Likewise, the sockets of TTP can be occupied by loops either from terminator or from another TTP subunit. To accomplish this bivalency, all loop-and-socket interactions use similar anchor points that consist of complementary hydrophobic and electrostatic sites (Fig. S22B,C). The overall loop configuration varies widely between terminator and TTP (Fig S22D), but these anchor points are found in all interaction pairs and likely account for a large proportion of the binding energy between these components.

#### Alterations to a capsid vertex allow it to accommodate the neck

Naively, we expected that in Oshimavirus, interaction between portal and MCP would be strong to prevent the portal from popping out of the capsid at high temperatures. We measured the buried surface area, solvation free energy, and numbers of hydrogen bonds or salt bridges in the interface between the portal and capsid in Oshimavirus and mesophilic phages P22 and T4. Surprisingly, the interaction between portal and the portal vertex is substantially weaker in Oshimavirus than in its mesophilic counterparts, with a smaller buried surface area (∼9,000 Å2 vs ∼28,000 Å2 and ∼16,000 Å2 respectively) and lower estimated hydrophobic interaction energy (−0.7 kcal/mol vs −121 kcal/mol and −67 kcal/mol respectively) (Table S3). Because the interaction area cannot explain the remarkable thermostability of Oshimavirus, we then examined the arrangement of subunits at the portal vertex.

Phage capsids must accommodate the neck at one vertex without sacrificing integrity. The capsid is composed of sub-assemblies of MCP called capsomers, which can be either pentameric (pentons) or hexameric (hexons), and which are held together through specific interactions of MCP (Fig. S24, 25A). The capsid contains eleven pentons, each at one of the 12 vertices of the icosahedron. Each penton is secured to 5 hexons using helical elements in the N-arms and S-loops of MCP (Fig. S25A,B). Dec protein strengthens the inter-capsomer interface, with the N-arms of two Dec subunits interacting with the E-loops of one penton MCP and one hexon MCP on the exterior surface of the capsid. The twelfth vertex, the portal vertex, replaces the penton with the aperture in which the neck sits (Fig. S25C), leaving the hexons to compensate for these important interactions that normally occur with a penton.

Key structural motifs of MCP and Dec are rearranged in the portal vertex to maintain stability without a penton (Fig. 3, S25). Instead of folding back to interact with its own P-domain (Fig. 3A), the MCP N-arm stretches into the neighboring hexon to interact with an S-loop there (Fig. 3B).Where the MCP N-arm and S-loop both form helices in typical hexons, in the portal vertex they interact through two β-strands, illustrating the conformational flexibility of these regions. This stands in stark contrast to the MCP N-arms of mesophilic phages, which either move very little at the portal vertex, or contract to consolidate interactions within the same capsomer (Fig. S26). Oshimavirus MCP is unique in creating new interactions with neighboring capsomer, creating an interwoven portal aperture. In addition, the Dec N-arm undergoes a large motion to thread through the aperture to the interior of the capsid, where it interacts with a hexon’s E-loop (Fig. 3C, S25D). In contrast, it interacts with an E-loop of the penton on the exterior of the capsid in a typical vertex.

**Figure 3:**
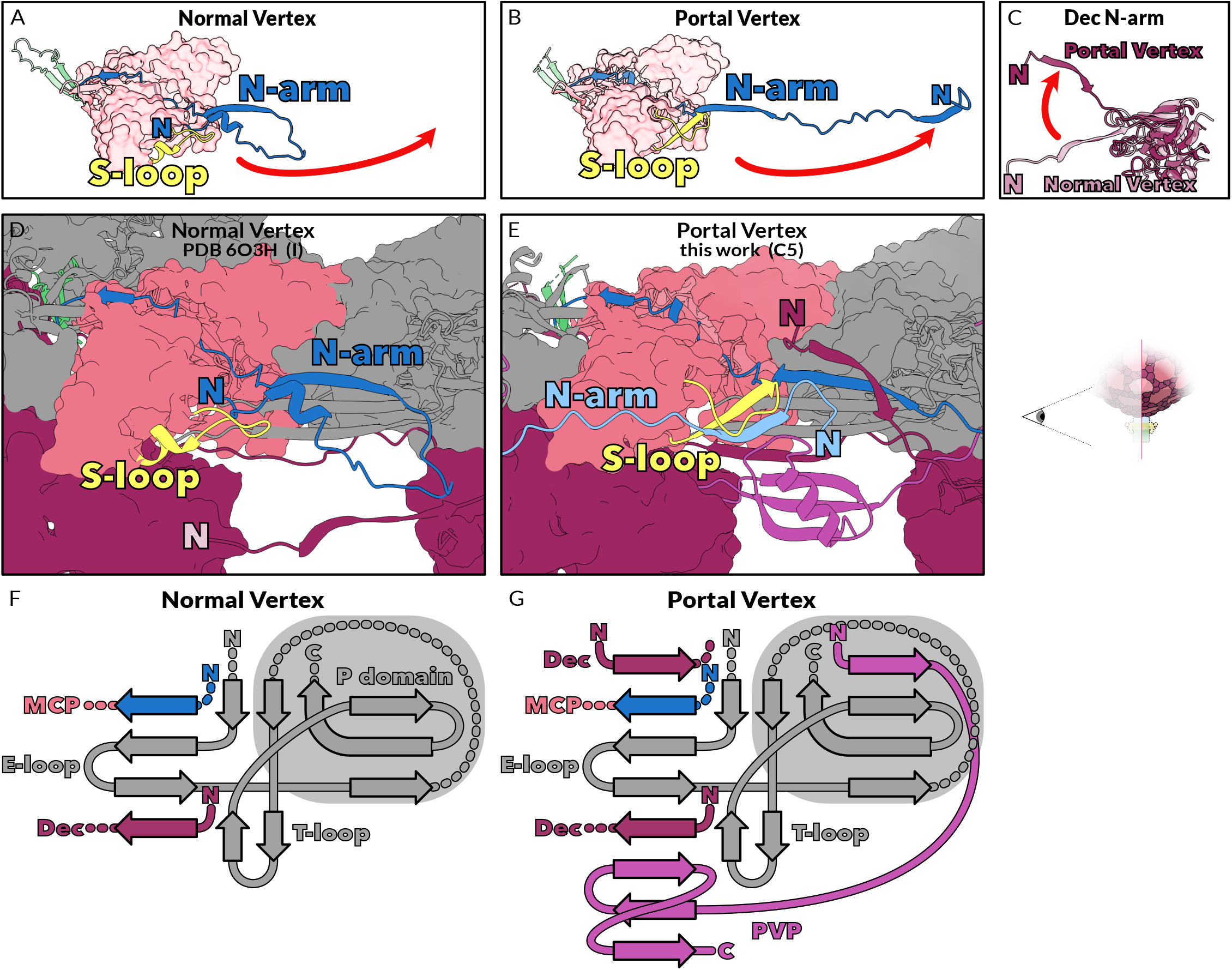
The MCP N-arm extends substantially in the portal vertex, which is strengthened by topological linkages between MCP, Dec, and PVP. **A. A single MCP subunit from a normal vertex of the capsid**. Loops and domains of MCP are colored as in figure S24, and the red arrow shows the direction and magnitude of the movement between the normal vertex and the portal vertex. **B. A single MCP subunit from the portal vertex of the capsid**. Loops and domains of MCP are colored as in figure S24, and the red arrow shows the direction and magnitude of the movement between the normal vertex and the portal vertex. **C. A single Dec subunit from the portal vertex (maroon) superimposed over a single Dec subunit from the normal vertex (faded maroon)**. The red arrow shows the direction and magnitude of the movement between the normal vertex and the portal vertex. **D. View of a normal capsid vertex, with penton subunits hidden, looking from the center of the penton towards a neighboring hexon**. One subunit of MCP (salmon), has its S-loop (yellow) and N-arm (blue) highlighted. The neighboring MCP subunit within the same capsomer is shown in grey, including its E-loop. **E. View of the portal vertex Pore from the center of the pore, with portal hidden, looking at the edge of the pore**. One subunit of MCP (salmon), has its S-loop (yellow) and N-arm (blue) highlighted. The neighboring MCP subunit within the same capsomer is shown in grey, including its E-loop. The N-arm of the counterclockwise neighboring capsomer (light blue) and the S-loop (yellow) interact with each other. The N-arm of PVP (purple) interacts with strands of the grey MCP P-domain. **F. Topology of β-strand interactions in icosahedral MCP**. The N-arm of MCP (blue), the E-loop of the adjacent MCP monomer (grey), and the N-arm of Dec (maroon, on the outer surface of the capsid) form a 4-stranded β-sheet. Dotted lines indicate that additional secondary structure elements were omitted for clarity. **G. Topology of β-strand interactions in the pore of the unique vertex**. The 4-stranded β-sheet becomes 8-stranded with the addition of the N-arm of another Dec on the inner surface of the capsid and 3 strands from PVP (magenta). The N-arm of PVP interacts with β-strands in the P domain of MCP.

The portal vertex is further stabilized by the addition of PVP at the edge of the aperture, which intertwines with MCP and Dec. Though the globular domain of PVP sits on the exterior of the pore, it also has an N-arm that folds through the aperture to join a β-sheet with the P-domain of MCP on the interior of the capsid (Fig. 3E, S25C,D). While in a normal vertex the MCP E-loop forms a four-stranded β-sheet with strands from the neighboring MCP N-arm and Dec N-arm (Fig. 3F), in the portal vertex this sheet becomes eight-stranded with the addition of 3 strands of PVP on the exterior of capsid and one strand of the Dec N-arm folded into the interior (Fig. 3G). Thus, all 3 components of the portal vertex are intricately interwoven, forming multivalent interactions that prevent the aperture from widening and maintaining structural integrity of the capsid. The intertwined topology of proteins stabilizing the portal vertex is reminiscent of the topological links that link the rest of the capsomers (29).

### Portal is repositioned in response to DNA packaging

Our C5 symmetric reconstruction of the neck reveals a loop of DNA around the portal wing region, with clearly visible major and minor grooves (Fig. S27A). All 3 capsid proteins (MCP, Dec, and PVP) directly interact with the DNA, thus imparting pseudo-5-fold symmetry to this segment of DNA. Lys2 of the N-arm of PVP inserts into the minor groove (Fig. S27C), while the T-loop of MCP inserts into the major groove (Fig. S27D), and the Dec N-arm contacts the DNA backbone (Fig. S27E). Because the DNA interacts so closely with the capsid and is positioned directly around the portal wing, it could be influencing the conformation of the portal vertex.

Our C1 reconstruction of the Oshimavirus neck reveals that the portal drastically alters how it contacts MCP during viral maturation. In the procapsid, the portal wing domain interacts with a T-loop (T-loop1) of MCP, while the portal clip domain interacts with the MCP S-loop and P-domain (Fig. 4A,C) (15). In contrast, in the mature virion the portal wing still contacts a T-loop of MCP, but it contacts the T-loop of a different MCP subunit within the same hexon (T-loop2). This movement allows T-loop1 to bind the pseudo-5-fold symmetric ring of packaged DNA wrapped around the portal (Fig. 4B,C). Because T-loop2 is lower in the capsid than T-loop1, portal must move toward the outside of capsid pore, pushing the clip domain outside of the capsid so that it does not bind MCP at all but is instead exposed to bind the adaptor C-terminal tail. When aligned on T-loop1, the geometric center of portal moves downward by over 22 Å (Fig. 5). This movement is not due to the internal motions of portal, because the geometric centers of the procapsid portal and mature virion portal change minimally when the two are aligned against each other (∼1 Å) (Fig. S28). We hypothesize that the repositioning of portal is driven by the internal pressure of packaged DNA.

**Figure 4:**
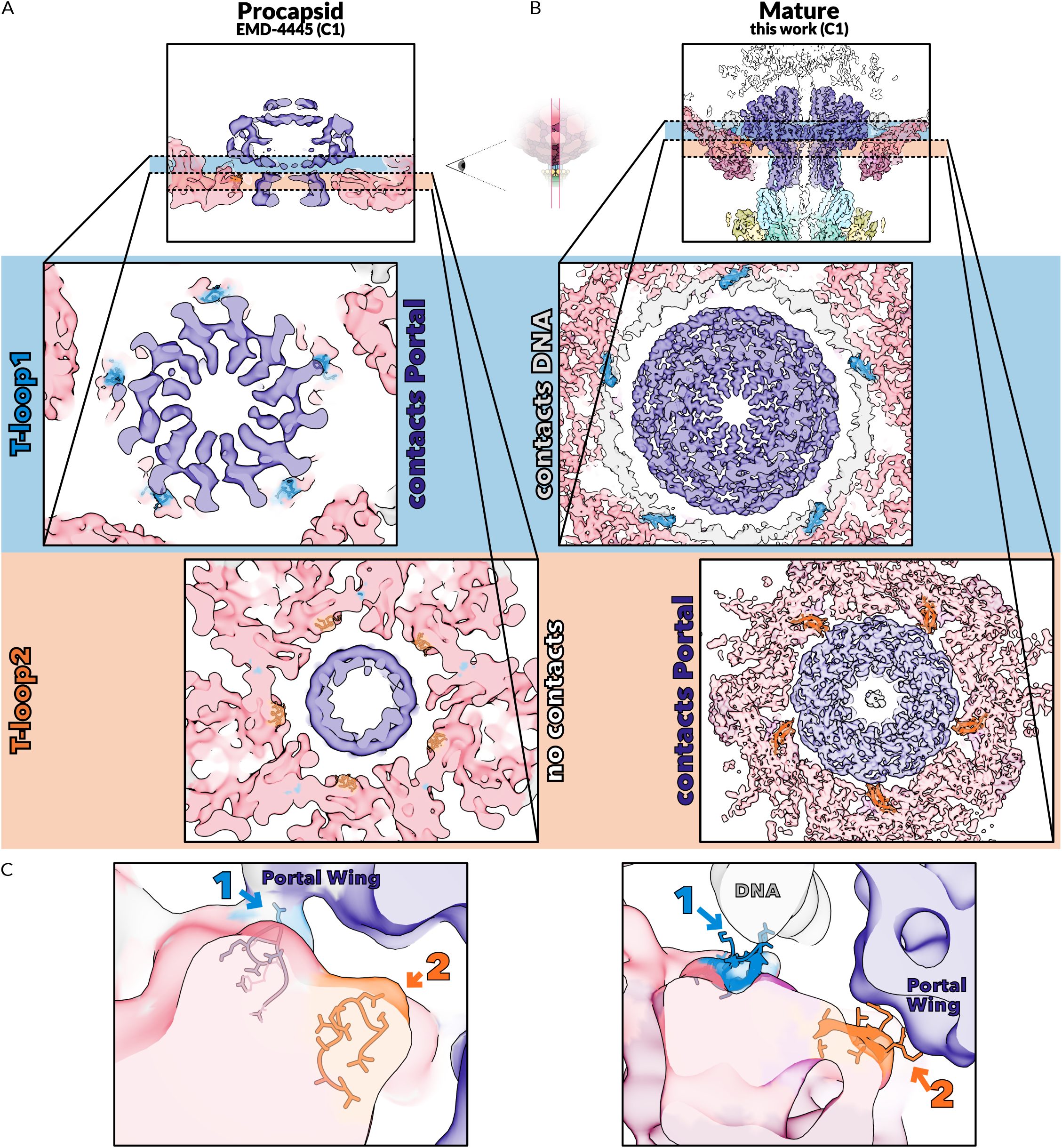
Different T-loops of MCP contact the portal wing in procapsids and mature virions. **A. Frontal and transverse slices of EM density for the procapsid and its portal (EMD-4445)**. EM density is shown as a transparent surface with MCP in salmon and portal in indigo. T-loops 1 and 2 are shown as cartoons with stick sidechains and colored as in blue and orange, respectively. The location of the T-loop within MCP and within the portal vertex are diagrammed in figures S22 and S24. Blue and orange shading highlight the areas shown in transverse sections. **B. Frontal and transverse slices of EM density for the mature virion (C1 reconstruction)**. T- loops 1 and 2 are depicted as in A and the EM density is shown as a transparent surface and colored by protein identity as in Figure 1. **C. Frontal slices of the regions of procapsids (left) and mature virions (right) that contact the portal**. The EM density for the procapsid and its portal (EMD-4445, left) and of the C1 mature portal and capsid (Gaussian filtered, right) are shown as transparent surfaces colored by protein identity. T-loops 1 and 2 are depicted as in A and B.

To determine whether this repositioning is due to the changes in the capsid conformation or to the pressure of packaged DNA, we compared the mature portal and portal vertex to those of expanded empty capsids, which have a capsid conformation similar to the mature virion but lack tails and packaged DNA (28). Empty expanded capsids can be isolated from phage preparations and most likely represent an assembly or disassembly byproduct. To compare the expanded empty capsid structure to the mature virion, we fit our model of the mature portal and capsid shell into the C5-symmetric map of expanded empty capsids (EMD4446) (28) (Fig. S29A).The mature capsid hexons fit the corresponding region in the map of expanded empty capsids, but the portal does not fit into the map, indicating that portal changes position between mature virions and expanded empty capsids. Compared to mature virions, the density for portal moves into the capsid lumen and less of the clip protrudes on the exterior of the capsid in capsids lacking packaged DNA.

Because the expanded empty capsid was reconstructed with C5 symmetry, it was not possible to determine the precise interactions between portal and MCP. However, portal is not close enough to T-loop1 to interact with it as it does in procapsids. To measure the position of portal in expanded empty capsids, we fit both the mature and procapsid portals (without MCP) into the expanded empty map (Fig. S29B). The geometric center of portal is at an intermediate position in expanded empty capsids, ∼9 Å more inward than in mature virions, and ∼14 Å more outward than in procapsids. The geometric centers of the mature and procapsid portals in the expanded empty map are only minimally different from each other (∼1 Å). Thus, in empty expanded capsids, we find that the portal takes an intermediate position, with more of the clip protruding from the capsid than in procapsids, but less than in mature capsids (Fig. S25).

To test whether portal movement is driven by tail attachment, we examined the structure of the filled, tail-less capsids (Fig. 2D, S30). This reconstruction has clear density for both packaged DNA and DNA passing through the portal pore loops. The atomic model of the mature portal and portal vertex fits this density quite well, indicating that the tail-less portal is in the same position as in mature virions. Thus, the repositioning of portal is not due to tail attachment. These results suggest that pressure from packaged DNA is a major driver of the portal repositioning during phage maturation.

We hypothesized that this motion provides a mechanism for Oshimavirus to terminate headful packaging. To test this hypothesis, we searched for pairs of procapsid and mature portal vertex structures in EMDB. We compared the repositioning of portal in Oshimavirus to P22, a headful phage, and Lambda and phi29, non-headful phages. We observe a downward motion of portal in P22 but not in Lambda and phi29 (Fig. 5). Thus, we identified a dramatic repositioning of the portal relative to the capsid shell in headful phages, which is coupled to the packaging of DNA. We propose that this forms a pressure-sensitive switch that triggers termination of headful packaging in a broad class of phages.

## DISCUSSION

We have visualized the neck of the thermophile Oshimavirus at high resolution using cryoEM. Though many of the components of this phage were previously unknown, we were able to identify them from our EM maps using the computational model building tool ModelAngelo. This achievement was made possible by the enormous strides in cryoEM methodology and demonstrates the utility of ModelAngelo for future studies of complexes with unknown components. Analysis of these proteins revealed homology to previously studied phage proteins with some surprising connections, such as the similarity between the portal vertex protein found in the capsid to an RNA Polymerase inhibitor from phage T7. These relationships underscore the modular construction of phages and the high degree of horizontal gene transfer, as well as the extent to which the diversity of phage proteins is still under sampled. Our analysis provides significant insights into how the DNA genome and the portal are retained within the capsid at high temperatures and how bacteriophage maturation is coordinated.

### Stabilization of the portal vertex

The portal vertex, found in the vast majority of phages as well as in pathogens of the Herpesvirus order, presents a conundrum for the viruses. The portal and the aperture in the capsid in which it sits are required to translocate the viral genome across the capsid shell, which occurs during both genome packaging and infection of the host cell. However, creating an opening in the capsid introduces a weak point that could compromise capsid stability, endangering the virus particle’s ability to persist long enough to infect a new host. The viral genome is packed tightly into the capsid, creating enormous pressure that the capsid must contain in order for the virus to survive. This pressure is exacerbated at high temperatures, which can also destabilize or even denature viral proteins. We previously described how the capsid proteins of a thermophilic phage have evolved to withstand high temperature by increasing topological connections between subunits (29). We therefore wondered how this phage would strengthen its portal vertex to withstand both high temperature and the pressure of its own packaged genome.

We observed that the interactions between portal and MCP in Oshimavirus are less numerous than in mesophilic phages. Instead, we observed that portal is held in the capsid by topological constraints between hexons that prevent dilation of the portal vertex aperture. If the aperture were to widen even slightly, it would allow the portal wing to pass through and thereby release the phage genome. To prevent this, P74-26 uses a highly intertwined network of interactions that lock the aperture width. The N-arm of MCP transitions from an intra-subunit interaction in a normal vertex to an inter-subunit interaction with a neighboring capsomer, moving the Ca of the N-terminal residue over 100 Å. Such conformational changes of MCP are not present in mesophilic phages. The N-arm of one subunit of Dec wraps through the aperture to interact with the interior of MCP. PVP is added to the exterior of portal vertex, and its N-arm also wraps under the aperture to interact with the interior of MCP. The formation of an 8-stranded sheet including strands from two MCP subunits, two Dec N-arms, and PVP highlights the interconnectedness of these interactions. Thus, these subunits are woven together as in a wicker basket, strengthening the aperture through the number and topology of interactions rather than strengthening individual interactions. This is similar to the topological linkages between MCP subunits described in Stone et al. (29).

We hypothesize that this mechanism of topological intertwinement could play a significant role in generating thermostability for other large assemblies that must withstand high temperatures. We further note that this strategy of interweaving structural elements is distinct from the mechanisms for generating thermostability in individual protein domains, which tends toward increased hydrophobic and electrostatic interactions or decreased conformational entropy (45–49). We propose that incorporating a topological component into future protein design efforts might allow for engineering of large protein assemblies with enhanced thermostability.

### Gating of DNA by the pore of portal

The portal is the gatekeeper of DNA retention in Oshimavirus. We observe that viral particles completely lacking necks and tails still retain DNA in the capsid despite the loss of the tail tip complex, indicating that the portal is the primary determinant of genome retention. Moreover, our structure shows that the portal adopts a closed pore conformation that would restrict the movement of DNA through the pore. This observation confirms a previous hypothesis that the closed pore conformation would be found in mature, DNA-filled virions (15). No other constrictions are observed along the length of the neck region. Thus, we demonstrate that Oshimavirus retains its DNA primarily using the portal and without the aid of other components that some phages use to retain their genome, such as the stopper protein (50) or the tail tip complex (43). Our work indicates that the signal for genome release must somehow transmit across the nearly micron-long tail of Oshimavirus to the portal, a remarkable feat of long-range communication. How this signal is transmitted will be the subject of future studies.

### Packaged DNA is coupled with portal repositioning

We observe pseudo-5-fold symmetric DNA wrapped around portal. The interactions in Oshimavirus do not appear to be sequence-specific, with primarily minor groove and backbone interactions. The sole major groove interaction occurs with insertion of an Asparagine and Lysine, which have limited potential for sequence-specific interactions. Oshimavirus uses a headful packaging mechanism, so individual virions have circularly permuted genome termini such that the sequence of this stretch of DNA is different in each virion (28). By binding to T-loop1, the DNA displaces the portal wing, which interacts with T-loop1 in the procapsid. In the mature capsid, portal now interacts with T-loop2 and moves outward in the aperture by ∼22Å. This outward motion is likely caused by DNA pressing down on the portal crown and displacing the portal wing from T-loop1.

In support of this hypothesis, the portal moves outward by ∼ 9 Å in mature capsids that are empty compared to mature virions that have packaged DNA, suggesting that ∼40% of portal repositioning is due to direct pressure from packaged DNA. We propose that the rest of portal repositioning is due to the expansion of the capsid, an event that is also triggered by DNA packaging (51). Our structural comparison suggests that movement of the portal wing from T-loop1 to T-loop2 is coupled to capsid expansion, since the T-loops only assume these positions when MCP undergoes conformational changes to transition from the procapsid to expanded capsid. Binding of the tail does not seem to contribute to portal repositioning at all. Thus, our analysis suggests that the pressure from packaged DNA plays a major role in altering the positioning of portal, both through direct pressure of DNA onto portal, as well as through the expansion of capsid. We propose that the portal repositioning due to capsid expansion is irreversible, while the portal repositioning from direct DNA pressure is reversible, as seen in the expanded empty capsids (Fig. S29). Repositioning of the portal complex with DNA packaging has also been observed in Herpes Simplex Virus-2 (52), Varicella-Zoster Virus (53), and Human Cytomegalovirus (54), but the mechanism for this movement is not clear.

### Speculative mechanism of the headful switch

Using the procapsid structure as the starting point and the mature virion structure as the end, we hypothesize a pathway for the headful switch. The procapsid consists of a compacted icosohedron of MCP, without Dec or PVP, and with portal at a single vertex. Scaffold protein is also present inside the procapsid, as well as any other co-assembling proteins. Over the course of maturation, several major changes take place, some of which may occur concomitantly. These include the expansion of the capsid, dissociation of the packaging motor, removal of scaffold protein, addition of Dec and PVP to the capsid, conformational and positional changes in the portal, and attachment of the virion tail.

Crucially, in the procapsid state the clip domain of portal is bound by MCP, which blocks interactions with the adaptor C-terminal tail and prevents further assembly of the neck. To fill the procapsid with the viral genome, the terminase packaging motor binds to the portal vertex and translocates DNA through portal. The terminase is a pentamer with each subunit containing an ATPase domain, a nuclease domain, and a flexible C-terminal tail (Fig. 6A) [(10, 11, 55)]. The ATPase domains oligomerize and translocate DNA through both terminase and portal. The nuclease domains are required to cleave DNA when packaging is complete.

**Figure 5:**
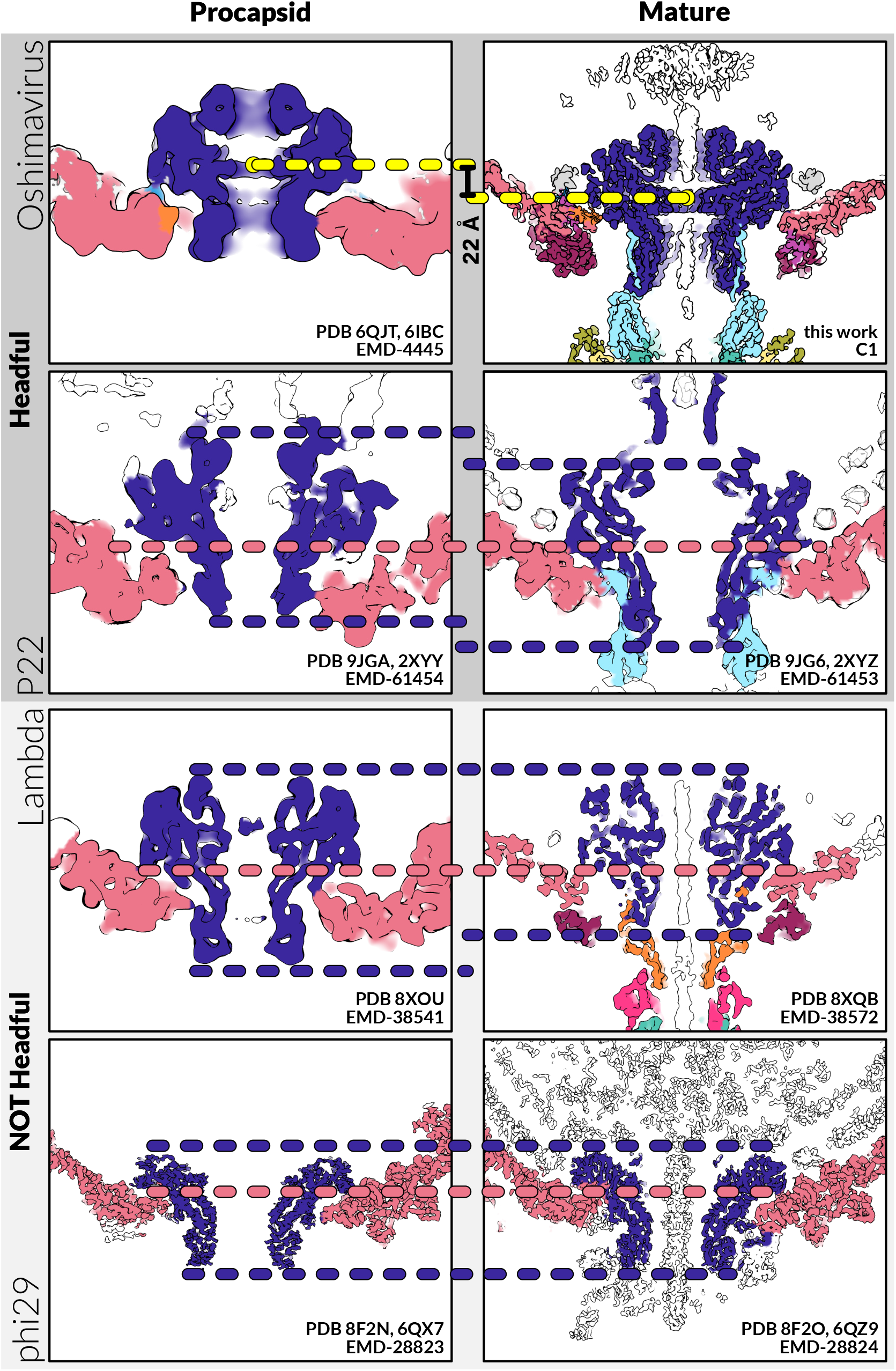
Portal moves downward during maturation and interacts with a different region of MCP. Frontal sections of reconstructions of Oshimavirus and three different bacteriophages. Oshimavirus and P22 package their genomes with a headful mechanism, while Lambda and phi29 do not. Oshimavirus is colored as in Figures 1 and 4, while Lambda and P22 are colored as in Figure S16. The Oshimavirus procapsid and mature virions were aligned on the position of T-loop1. The models of the Oshimavirus procapsid and mature portals were fit into their electron density and the geometric centers were calculated in ChimeraX and depicted with yellow spheres. The distance between their centers is 22Å. Since the other phages have different regions of contact between portal and MCP, the inner surfaces of their capsids and procapsids were visually aligned (salmon dotted lines), and therefore precise distances between portals could not be calculated. The top and bottom of these portals are indicated with indigo dotted lines to visually compare their positions.

**Fig. 6:**
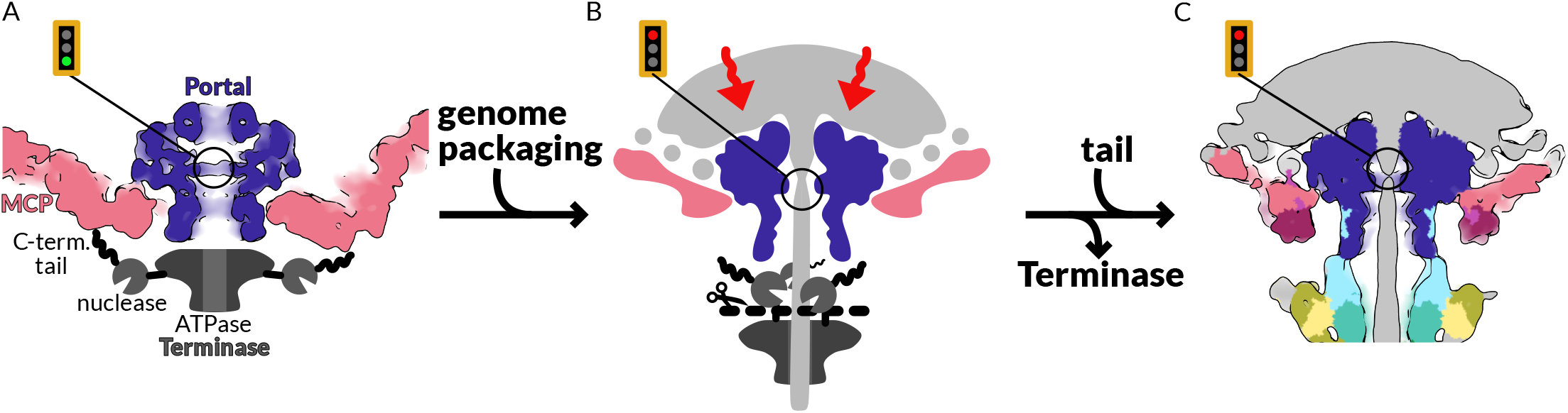
Speculative model of the phage neck maturation pathway. **A. In order for genome packaging to occur, the Terminase motor must attach to the Portal in the vertex of a procapsid**. EM density (EMDB: 4445) of the Portal and procapsid is cross-sectioned and colored according to protein identity as in Figure 1. The Terminase is schematized in grey and black. Scaffold protein and DNA are not shown. In the procapsid, the Portal pore is open to allow DNA packaging. The clip region of Portal, to which Adaptor tails will eventually bind, is occluded by the procapsid. The Terminase is pentameric, which we propose allows it to interact with the 5-fold vertex of the procapsid through its C-terminal tails. This tethers the nuclease domains away from the central channel through which DNA will be translocated. The Terminase can then package DNA into the capsid. During packaging, the capsid also expands. **B. We propose that pressure of the packaged DNA changes the position of Portal, switching it from genome packaging to allow Tail attachment**. When the capsid is full, the enormous pressure of packaged DNA (light grey) presses the Portal downwards like a piston. We propose that this disengages the C-terminal tails of Terminase from the capsid, freeing the nuclease domain so that it can access DNA. Once DNA has been cleaved, the Terminase and unpackaged DNA dissociate from the packaged capsid. After downward motion of the Portal and dissociation of the Terminase, the clip region is now available to bind Adaptor tails. **C. The tail is attached to the fully packaged capsid**. EM density (C1 reconstruction, gaussian filtered) of the mature Portal vertex and Neck is cross-sectioned and colored according to protein identity as in Figure 1.

Though no structures of this complex exist, we speculate that the 5 C-terminal tails each bind to the 5-fold symmetric hexons arrayed around the portal vertex. This would have two benefits: securing the pentameric terminase to the dodecameric portal by tethering it to the C5 capsid, and anchoring the nuclease domains away from DNA to prevent premature cleavage during translocation. We previously hypothesized that the headful signal facilitates dissociation of the terminase from portal, releasing the nuclease domain from a restricted conformation and allowing the nuclease domains to reorient and cut DNA (26).

Here, we propose that the pressure-sensitive outward motion of portal is the headful signal that induces the switch from ATP-dependent DNA translocation to DNA cleavage. Capsid expansion and the pressure of packaged DNA push the portal outward and shifts its interaction from T-loop1 to T-loop2 (Fig. 6B). This large motion pushes the clip domain of portal out of the aperture so that it now protrudes past the capsid. We propose that this motion dislodges the terminase tails from the hexons, freeing the nuclease domains. The nuclease domains then reposition to cleave DNA, dissociating the terminase and unpackaged DNA from the filled capsid, such that they can bind to another procapsid for genome packaging. Likewise, the portal clip is now free for adaptor tails to bind, attaching the filled head to the tail (Fig 6C). The downward motion of portal is conserved in headful phage P22 but not in non-headful phages Lambda and phi29. Thus, we propose that the pressure-sensitive motions reported in this study underscore the headful packaging signal in a broad class of phages.

Additional mechanisms have been proposed for the headful signal, which are not mutually exclusive with that proposed here. Studies of phage T4 have led to the hypothesis that motions between the TerL ATPase and nuclease domains can switch between these two activities (56). Furthermore, numerous studies indicate that internal conformational changes within portal impact the headful signal (42, 57). It is likely that multiple redundant mechanisms are used regulate the headful switch in order to prevent premature cleavage of the genome during packaging. Future studies will test these mechanistic models.

## MATERIALS AND METHODS

### Experimental Design

The objective of this study was to determine the structure of the Oshimavirus P74-26 neck complex. We were interested in this structure because our studies of the P74-26 capsid and tail yielded valuable insights into mechanisms of thermostability and viral assembly, and structure of the neck complex connecting the capsid and tail was still unknown. To obtain this structure, we performed cryoEM on purified Oshimavirus P74-26 virions. We treated the neck region of each virion as a particle and applied single particle cryoEM analysis techniques. We then used these EM maps to determine the identity of the proteins composing the neck complex, and confirmed those identifications with Mass Spectroscopy, as described below.

### Virus purification

P74-26 phage was propagated in the host strain T. thermophilus HB8 (ATCC 27634) using fresh overnight cultures grown at 65 °C in Thermus growth medium (4 g/L yeast extract, 8 g/L tryptone, 3 g/L NaCl, 1 mM MgCl2, 0.5 mM CaCl2). Four milliliters of P74-26 phage stock at 1×106 plaque-forming units per mL (PFU/mL) was combined with 6 mL of fresh T. thermophilus HB8 and incubated at 65 °C for 10 min. The mixture was then inoculated into 1 L of Thermus Medium and incubated at 65 °C while shaking for 4–5 h. The culture containing lysed cells and phage was then spun at 4000 ×g for 20 min, and supernatant was incubated with DNase I (2 U/mL) and RNase A (1 μg/mL) for 1 h at 30 °C. To precipitate virions, solid NaCl was added to 1 M final concentration and Polyethylene Glycol MW 8000 was gradually added to a final concentration of 10% (w/v). The mixture was incubated on ice overnight. The next day, precipitated phage stock was spun in 50 ml aliquots at 11,000 ×g for 20 min at 4 °C. The resulting phage pellet was resuspended in 2 mL of 50 mM Tris pH 7.5, 100 mM NaCl, and 1 mM MgSO4. The resuspension was supplemented with 0.4 g solid CsCl and added to a CsCl step gradient (2 mL steps each of 1.2, 1.3, 1.4, 1.5 g/mL CsCl and 1mL cushion of 1.7 g/mL CsCl, in 50mM Tris pH 7.5, 100mM NaCl, 1mM MgSO4). Gradients were prepared in 12 mL ultracentrifuge tubes and spun in a Beckman SW40-Ti rotor at 38,000 RPM for 18 h at 4°C. The virion band was isolated by fractionation and dialyzed twice overnight into 2L of 50 mM Tris pH 8.0, 10 mM NaCl, 10 mM MgCl2 at 4°C. P74-26 virions were then concentrated to 1×1012 PFU/mL for subsequent use in electron microscopy.

### Electron microscopy

Grids were glow discharged on a PELCO easiGlow (Ted Pella) at 25 mA for 45 s (negative polarity). 3.5 μL of purified virions at 1×1010 PFU/mL was applied to a 400 mesh Lacey Carbon grid (Electron Microscopy Sciences) at 10 °C with 90% humidity in a Vitrobot Mark IV (FEI). Sample was blotted from both sides for 10 s after a wait time of 15 s, then immediately vitrified by plunging into liquid ethane. Micrographs were collected on a 300 kV Titan Krios electron microscope (FEI) equipped with a K3 Summit direct electron detector (Gatan) and a Gatan Bioquantum Imaging Filter set to a slit width of 20 eV. Images were collected at a magnification of 81000x in super-resolution mode with an unbinned pixel size of 0.53 Å per pixel and a total dose of 49.0571 e−/Å2 per micrograph, with a target defocus range set from −0.4 to −1.5 μm. In total, 16,184 micrographs, each 30 frames, were collected using SerialEM.

Broken phage samples were applied to either a 200 mesh Lacey Carbon grid (Heavy Band 1) or a 200 mesh UltrAuFoil grid (Heavy band 2) and imaged on a 200kV Glacios electron microscope (FEI) equipped with a Falcon4 (Thermo Fisher) direct electron detector and a Selectris energy filter operated at 10 eV. The Falcon4 camera was operated in electron event representation (EER) mode and micrographs were collected at 79,000x magnification, a pixel size of 1.53 Å, a target defocus range set from −1.0 to −2.0 μm, and a total dose of 49.2668 e−/Å2. In total, 1,814 micrographs, each 30 frames, were collected using SerialEM.

Negative Stain EM: Carbon-coated 200 mesh copper grids (Electron Microscopy Sciences) were glow discharged on a PELCO easiGlow (Ted Pella) at 20 mA for 20 s (negative polarity). 4 μL of sample was applied to the grid and incubated for 1 min. Excess sample was blotted, then the grid was washed with water followed by staining with 1% uranyl acetate (pH 4.5). Grids were viewed with a Philips CM120 electron microscope at 120 kV on a Gatan Orius SC1000 camera.

### Data processing and reconstruction

All processing was performed in CryoSPARC4 (58). Micrographs were preprocessed using Patch Motion Correction and Patch CTF Estimation. We first reconstructed the entire capsid with icosahedral symmetry. An initial stack of 208,224 particles was extracted using Blob Picker with a diameter of 77-87 nm (the P74-26 capsid has diameter 82 nm) and cropped in Fourier Space (F-cropped) from a box size of 1260 pixels (px) to 256 px to reduce computational load. After 2 rounds of 2D classification, we selected 31,014 particles and used their 2D classes as a template. Template Picker extracted a stack of 677,203 particles, which yielded 38,732 capsid particles after 2 rounds of 2D classification. This stack was reconstructed and refined with icosahedral symmetry. We then used symmetry expansion around the icosahedral space group and 3 rounds of 3D classification with a spherical focus mask over one vertex of the icosohedron to select particles with density protruding from the capsid at this vertex. After removing duplicates, 34,452 particles were reconstructed to show the capsid and Neck. Particles were then re-centered and re-extracted with a box size of 360 px to isolate the Neck and exclude most of the capsid density. These particles were then used to train a convolutional neural network via the Topaz wrapper available in CryoSPARC to pick Necks (59). We then used the model to pick 116,158 particles from the entire dataset, which were F-cropped from a box size of 360 px to 180 px to reduce computational load. 2D classification removed false positives and low-quality particles, yielding a stack of 41,024 Neck particles.

To visualize the Neck, we reconstructed and refined this stack with C12 symmetry using CryoSPARC Non-Uniform Refinement and re-extracted the particles with a box size of 360 px (uncropped) (60). 3D classification into 3 classes to remove junk left 39,678 particles. Subsequent 3D classification into 2 classes with a focus mask over the collar region separated Neck particles with and without the collar region. C12 symmetry expansion followed by 3D classification with a focus mask applied to the neck lumen revealed double-stranded B-form DNA in the central channel. The clearest class was selected for further processing. After duplicate removal, 19,306 particles were reconstructed and refined using Local Refinement. To resolve the transition between C12 and C3 symmetry, we performed symmetry expansion of the particle set around the C12 symmetry group followed by 3D classification into 10 classes with a focus mask applied to the collar region. Four resulting classes contained the P74-26 Neck, while the remaining 6 contained junk. After duplicate removal, the largest class had 30,748 particles and was refined with C3 symmetry using CryoSPARC Non-Uniform Refinement.

To visualize the Portal vertex, we reconstructed and refined the particle set with C5 symmetry using CryoSPARC Non-Uniform Refinement and re-extracted the particles with a box size of 360 px (uncropped). 3D classification with 3 classes to remove junk left 40,773 particles. To resolve the transition between C12 and C5 symmetry, we performed symmetry expansion of the particle set around the C12 symmetry group followed by 3D classification into 10 classes with a focus mask applied to the Neck. Four resulting classes contained the P74-26 Neck and Portal vertex, while the remaining 6 contained junk. After duplicate removal, the largest class had 39,382 particles and was asymmetrically refined using CryoSPARC Non-Uniform Refinement.

We also noticed particles in our micrographs that resembled the C12 Neck without any capsid. We manually picked 303 of these particles from micrographs collected at a range of defocus values. These particles were used to train a Topaz model which was then used to extract 163,113 particles, of which 12,754 contained the capsid-less Neck after 2 rounds of 2D classification. This particle set was used to train a second, independent Topaz model which then was used to extract 280,700 particles, of which 26,019 contained the capsid-less Neck after 3 rounds of 2D classification. The above refinement and symmetry-breaking pipeline was applied to this stack of particles to yield C12 and C3 reconstructions of the capsid-less Neck.

The reconstruction of broken-tailed phages was performed as above. Micrographs were preprocessed using Patch Motion Correction and Patch CTF Estimation. We first reconstructed the entire capsid with icosahedral symmetry. An initial stack of 39,870 particles was extracted using Blob Picker with a diameter of 77-87 nm (the P74-26 capsid has diameter 82 nm) and cropped in Fourier Space (F-cropped) from a box size of 882 pixels (px) to 256 px to reduce computational load. After 2 rounds of 2D classification, we selected 14,351 particles and used their 2D classes as a template. Template Picker extracted a stack of 113,830 particles, which yielded 15,679 capsid particles after 2 rounds of 2D classification. This stack was reconstructed and refined with icosahedral symmetry. We then used symmetry expansion around the icosahedral space group and 2 rounds of 3D classification with 10 classes and a spherical focus mask over one vertex of the icosohedron to select particles with density protruding from the capsid at this vertex. After regrouping similar classes and removing duplicates, 13,186 particles were reconstructed to show the capsid and neck and 3,964 particles were reconstructed to show the capsid and portal without necks. Particles were cropped in Fourier Space (F-cropped) from a box size of 882 pixels (px) to 512 px and refined using Local Refinement.

### Identification of Neck proteins

To identify the unknown proteins comprising the Neck, we used the automated cryo-EM atomic model-building software ModelAngelo (34). We ran ModelAngelo without sequence inputs on C3, and C5 maps of the Neck, and extracted the sequences of all built peptides >10 residues in length in FASTA format. These peptide sequences were used to search the non-redundant RefSeq protein database of the NCBI BLAST server. Hits are reported in Supplementary Table 1.

### Mass spectrometry

Mature P74-26 virion samples were precipitated out of solution using Trichloroacetic acid (TCA) precipitation. 1 volume of TCA was added to 4 volumes of protein sample and incubated for 10 min at 4 ºC. Samples were spun down at 13,000 RPM in a microcentrifuge for 5 min and the resulting pellet was washed 3 times with 200 μL of ice-cold acetone. The pellet was air dried until it was white and cloudy. Pellets were lyophilized and shipped on dry ice for mass spectrometry analysis at the Taplin Biological Mass Spectrometry Facility.

50 μL of 50 mM ammonium bicarbonate with 10% acetonitrile was added to the dry tubes containing the TCA precipitated protein and gently vortexed. 10 μL of 20 ng/μL modified sequencing-grade trypsin (Promega, Madison, WI) was spiked into the solutions and the samples were incubated at 37 ºC overnight. Samples were acidified by adding 5 μL 20% formic acid solution and then desalted by STAGE tip(61). Before analysis, the samples were reconstituted in 10 µL of HPLC solvent A. A nano-scale reverse-phase HPLC capillary column was created by packing 2.6 µm C18 spherical silica beads into a fused silica capillary (100 µm inner diameter x ∼30 cm length) with a flame-drawn tip(62). After equilibrating the column each sample was loaded via a Famos auto sampler (LC Packings, San Francisco CA) onto the column. A gradient was formed, and peptides were eluted with increasing concentrations of solvent B (97.5% acetonitrile, 0.1% formic acid). As peptides eluted, they were subjected to electrospray ionization and then entered into an LTQ Orbitrap Velos Elite ion-trap mass spectrometer (Thermo Fisher Scientific, Waltham, MA). Peptides were detected, isolated, and fragmented to produce a tandem mass spectrum of specific fragment ions for each peptide. Peptide sequences were determined by matching protein databases with the acquired fragmentation pattern by the software program, Sequest (Thermo Fisher Scientific, Waltham, MA) (63). All databases include a reversed version of all the sequences and the data was filtered to between a one and two percent peptide false discovery rate. The number of sequenced peptides mapping to each protein is reported in Supplementary Table 2.

### Model building and refinement

We ran ModelAngelo using the consensus sequence of each identified protein on C3 and C5 maps of the Neck. These models were then Real-Space Refined in Phenix and used as the starting point for manual refinement in Coot and ISOLDE (64–66). ISOLDE was accessed as a plug-in to ChimeraX (67, 68). We selected a single subunit of each protein for manual refinement based on the number of residues built and the geometry statistics. Each subunit was manually built in Coot in the highest-resolution map available. For example, Portal was built in the C12 map rather than the C3 map. Though most of Adaptor has C12 symmetry, regions that contact Terminator have C6 symmetry. After Adaptor was built in the C12 map, it was duplicated, rotated around the symmetry axis, and rigid body fit into the density for the neighboring subunit so that the differing loops could be rebuilt into their density. The same workflow was used to rebuild the loops of Terminator where they contact TTP with C3 symmetry. ISOLDE simulations were also used to fix phi/psi angles and to resolve clashes of selected loops (66). The asymmetric units were then duplicated, rotated around the symmetry axis, and fit into the density for neighboring units to generate the complete complex. The neck complex was Real-Space Refined in the C3 map and the portal vertex complex was Real-Space Refined in the C5 map. The asymmetric map was used to align the C3 and C5 maps and models relative to each other. Loops of MCP and Portal that contact each other were rebuilt into the asymmetric map in ISOLDE. The capsid-portal complex was Real-Space Refined in Phenix for a final time in the asymmetric map, with reference model restraints applied to the entire model except for the loops that were rebuilt in the asymmetric map. A 66-bp segment of ideal B-form DNA was generated in Coot and fit into the central DNA in ChimeraX. This model was merged with the composite model of the neck and portal vertex and simulated in ISOLDE with all other components of the neck and portal vertex restrained to their initial positions. The central DNA was Real-Space Refined with reference model restrains applied to the rest of the complex. Final refinement statistics are reported in Supplementary Table 3.

#### Modeling the procapsid Portal and Portal Vertex

5 hexamers, without the pentamer they encircle, were extracted from the published model of the icosahedral P23-45 procapsid (PDB 6IBC). This model was fit into the asymmetric procapsid reconstruction (EMD-4445) around the density for Portal, along with the C12 model of the procapsid Portal (PDB 6QJT), to create a composite model of the procapsid Portal and Portal Vertex. Asymmetric loops of MCP and Portal were adjusted in ISOLDE to resolve clashes and fit EMD-4445.

### Statistical Analysis

No statistical analyses were performed during the course of this study, with the exception of those incidental to the computational methods listed above.

## SUPPLEMENTARY MATERIAL

**This pdf file includes**:

Figures S1 to S30

Tables S1 to S5

## ACKNOWLEDGMENTS

We thank Dr. Joshua Pajak and members of the Kelch, Grigorieff, and Schiffer labs for helpful discussions. We thank Dr. Terje Dokland and Dr. James Kizziah for helpful discussions. We thank Dr. Christna Ouch, Dr. Gregory Hendricks, and Dr. KyoungHwan Lee for assistance with data collection.

## Funding

This work was supported by National Science Foundation awards (MCB-1817338 and MCB-2342927) and the National Institutes of Health (R35GM156361) to B.A.K. The Titan Krios and Glacios at the UMass Chan Medical School Cryo-EM core facility, which were used to collect the data cited in this study, were funded by a grant from the Massachusetts Life Sciences Center. Negative Stain TEM was performed at the Electron Microscopy Core Facility at UMass Chan Medical School, which receives financial support from NIH (Specific Grant #) and the UMMS Chan Office of Research. Molecular graphics and analyses performed with UCSF ChimeraX, developed by the Resource for Biocomputing, Visualization, and Informatics at the University of California, San Francisco, with support from National Institutes of Health R01-GM129325 and the Office of Cyber Infrastructure and Computational Biology, National Institute of Allergy and Infectious Diseases.

## Author contributions

Conceptualization: E.L.S. and B.A.K. Formal Analysis: E.L.S. Funding Acquisition: B.A.K. Investigation: E.L.S., E.A., J.E.H., R.A., X.C., and K.S. Methodology : X.C. Visualization: E.L.S. Supervision: B.A.K. Writing—original draft: E.L.S., B.A.K. Writing—review & editing: E.L.S., E.A., R.A., K.S., B.A.K.

## Competing interests

Authors declare that they have no competing interests.

## Data and materials availability

EM maps and atomic models have been deposited to the Electron Microscopy Data Bank (EMDB) and Protein Data Bank (PDB) and will be released upon publication.

## Notes

### Competing Interest Statement

The authors have declared no competing interest.

### Summary of Updates

We have added additional analysis comparing our structures to previously published structures of other phages. These analyses broaden our conclusions to apply to a broad class of phages, those that package their genomes using a headful mechanism. For more details, we have included a changelog in the Supplemental Materials.

