## Supplementary Materials for "The structure of a thermostable phage’s portal vertex and neck complex illuminates the headful maturation mechanism"

bioRxiv

Emma Sedivy, *et al.*

**This PDF file includes:**

Supplemental text: version changelog

Figures S1 to S30

Tables S1 to S5

### ADDED DATA/ANALYSES:

•We resolved C1 density for the DNA running through the central channel of the neck.
•We reconstructed EM volumes of the filled phage heads with broken tails.
•We also show that the portal position in filled heads with no necks is the same as in mature phage, indicating that repositioning of the portal is driven by "pushing" by DNA rather than "pulling" by the tail.
•We measured the buried surface area, solvation free energy, and numbers of hydrogen bonds and salt bridges in the interface between capsid and portal in P74-26 and 2 mesophilic phages, T4 and P22. This interface is substantially weaker in P74-26 than in T4 and P22, despite being hyper-thermophilic. Thus, topological interweaving is the primary method of generating stability at the portal vertex.
•We estimated the motion of portal before and after packaging in two non-headful phages, Lambda and phi29, and one headful phage, P22. Portal repositions in P22 and P74-26, both headful phages, but does not move in non-headful phages Lambda and phi29.

### ADDED FIGURES:

•C1 DNA in the central channel was added to the composite density in figure 1C.
    •A supplementary figure detailing the reconstruction process was added as figure S4.
    •Local resolution maps of this reconstruction were added as panel S5C.
•A map of filled, tail-less phage heads was added as an inset to figure 2D.
    •The reconstruction of filled tail-less and filled broken-tailed phage heads was added as figure S14.
    •Figure S30 compares the position of portal in tail-less and broken-tailed heads to mature phage.
•Version 1 Figure 3 was moved to figure S25.
•Version 2 Figure 3 was made by combining Version 1 Figure S22 with 3 new panels showing the movement of N-arms in MCP and Dec.
    •Figure S26 was added to compare the change in MCP N-arm conformation between normal and portal vertices in a selection of mesophilic phages.
•Version 1 Figure 4 was moved to figure S27.
•Version 2 Figure 4 was made by combining Version 1 Figure S23 with Version 1 Figure 5B.
•Panels comparing motion of portal before and after packaging in different phages was added to Figure 5.
•A supplemental figure comparing the neck channels of different siphophages was added as figure S23.

### ADDED TEXT:

•Description of reconstructions with DNA in the central channel was added to Results section "The Oshimavirus neck includes regions of C12, C6, C5, and C3 symmetry" and Methods.
•Description of reconstructions of phage with broken tails was added to Results section "Portal is in the closed conformation" and Methods.
•Quantification of interactions at the portal vertex was added to Results section "Alterations to a capsid vertex allow it to accommodate the neck."
•Comparison between Oshimavirus MCP N-arm and the MCP N-arm of mesophilic phages was added to Results section "Alterations to a capsid vertex allow it to accommodate the neck."
•Comparison of portal position in filled phage capsids without necks and tails was added to Results section "Portal is repositioned in response to DNA packaging."
•Comparison of portal position in Oshimavirus to P22, Lambda, and phi29 was added to Results section "Portal is repositioned in response to DNA packaging."

### MINOR CHANGES:

•Added protein names and ORF numbers to the genetic map in figure 1B
•Panels 2A and B were updated to better orient the viewer to what part of the portal is being viewed in B.
•Removed capitalization (treating them as proper nouns) of component parts of the phage throughout
•Replaced "particle stack" with "particle set" throughout
•In the results section "Portal is in the closed conformation", added the names of phages with comparable structures
•In results section "Portal is in the closed conformation", specified and named the residues hypothesized by Bayfield et al. 2020 to play a role in genome retention, which our data also supports
•In results section "Consecutive rings transition...", changed the description of loop-and-socket interactions in consecutive rings of the neck from "promiscuous" to "bivalent" to convey that they can take either of 2 states, rather than a multitude of different states
•In results section "Portal is repositioned in response to DNA packaging", specified that the C1 reconstruction was used for this analysis
•In discussion section "Stabilization of the portal vertex", removed suggestion that multiple energetically equivalent registers of portal and capsid "may facilitate assembly and stability" and specified that this analysis was done by Bayfield et al.
•In the section beginning "Consecutive rings transition ...", changed description of loop-and-socket interactions from "promiscuous" to "bivalent" to reflect that the terminator loop can make 2 interactions, rather than only 1
•Minor changes for clarity and readability throughout

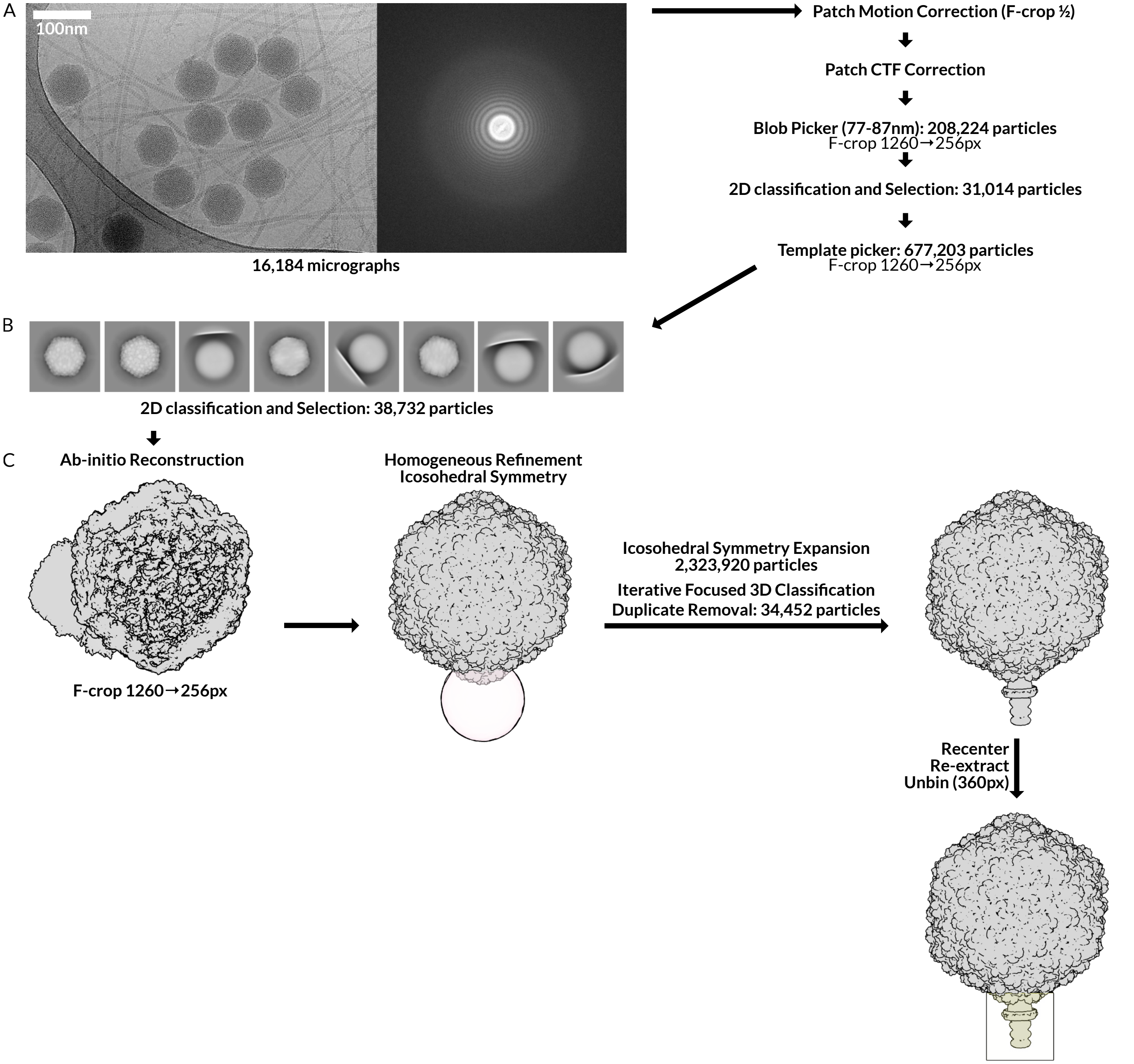


**Figure S1: Focused 3D classification of the phage capsid allows reconstruction of the P74-26 neck. A. Representative micrograph and pre-processing workflow.** We applied purified virions to lacy carbon grids and imaged them on the Titan Krios (Methods). One representative micrograph, with its Fourier transform, of a total 16,174 collected micrographs, is shown. We performed Patch Motion Correction and Patch CTF correction and used Blob Picker and Template picker to pick capsid particles in CryoSPARC . **B. Selected 2D classes.** We used 2D classification to select 38,732 particles representing filled capsids. Four of the eight selected classes also show lacy carbon edge from the grid film. **C. Reconstruction, refinement, and 3D classification of the whole capsid and neck.** First, we performed an asymmetric *ab-initio* reconstruction on the particle stack in B after cropping each particle in Fourier space (F-cropping) from a box size of1260 pixels to 256 pixels to reduce computational time and memory usage. We then refined this *ab-initio* map with icosahedral symmetry enforced, and performed symmetry expansion around the same space group. We placed a spherical mask (pink) over a single vertex of the capsid, and used focused 3D classification to select particles with density for the neck at this vertex. We performed three rounds of iterative 3D classification using the same focus mask. We removed duplicate particles before reconstructing the particle stack. Finally, we recentered this volume upon the neck and re-extracted a new particle stack without F-cropping at a box size of 360 pixels. The size and location of the new particle box is shown in yellow.

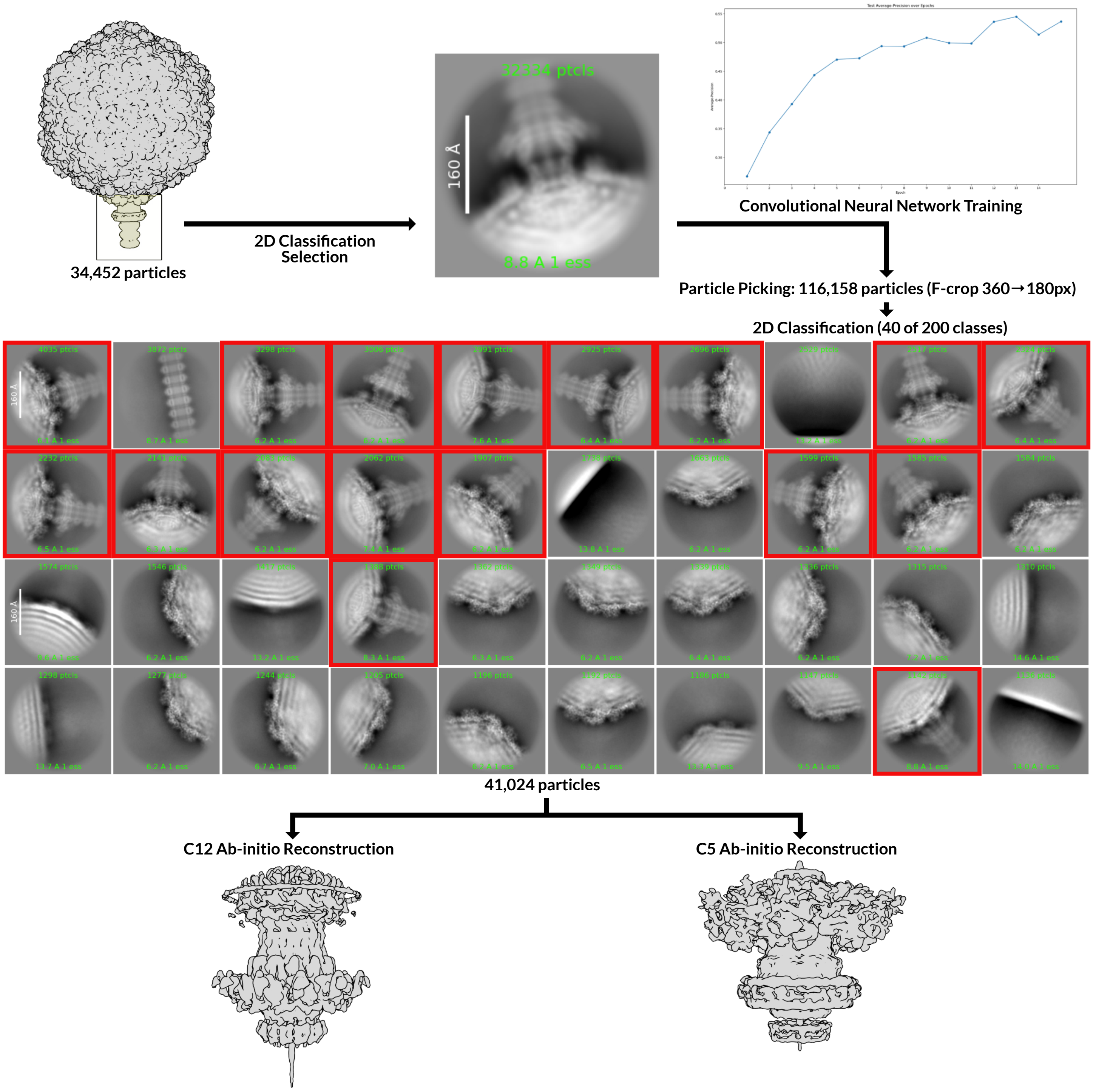


**Figure S2: Deep neural network particle picking increases the number of neck particles.** We screened the particle stack generated in Figure S1 using 2D classification to generate a stack of 32,334 particles with the 2D average shown at center. We used this particle stack to train a convolutional neural network using the Topaz wrapper in CryoSPARC and used the trained model to pick particles from the entire dataset. We F-cropped the picked particles by a factor of ½ to reduce computational time and memory usage and 2D classified into 200 classes, of which 40 are shown. 19 of these contained the neck (red boxes), generating a particle stack of 41,024. To avoid flattening caused by the highly symmetrical particle, we applied symmetry to the ab-initio reconstructions. Based on the known symmetries of the neck, we performed a C12 *ab-initio* reconstruction to visualize the portal and associated proteins, and a C5 *ab-initio* reconstruction to view the capsid and associated components.

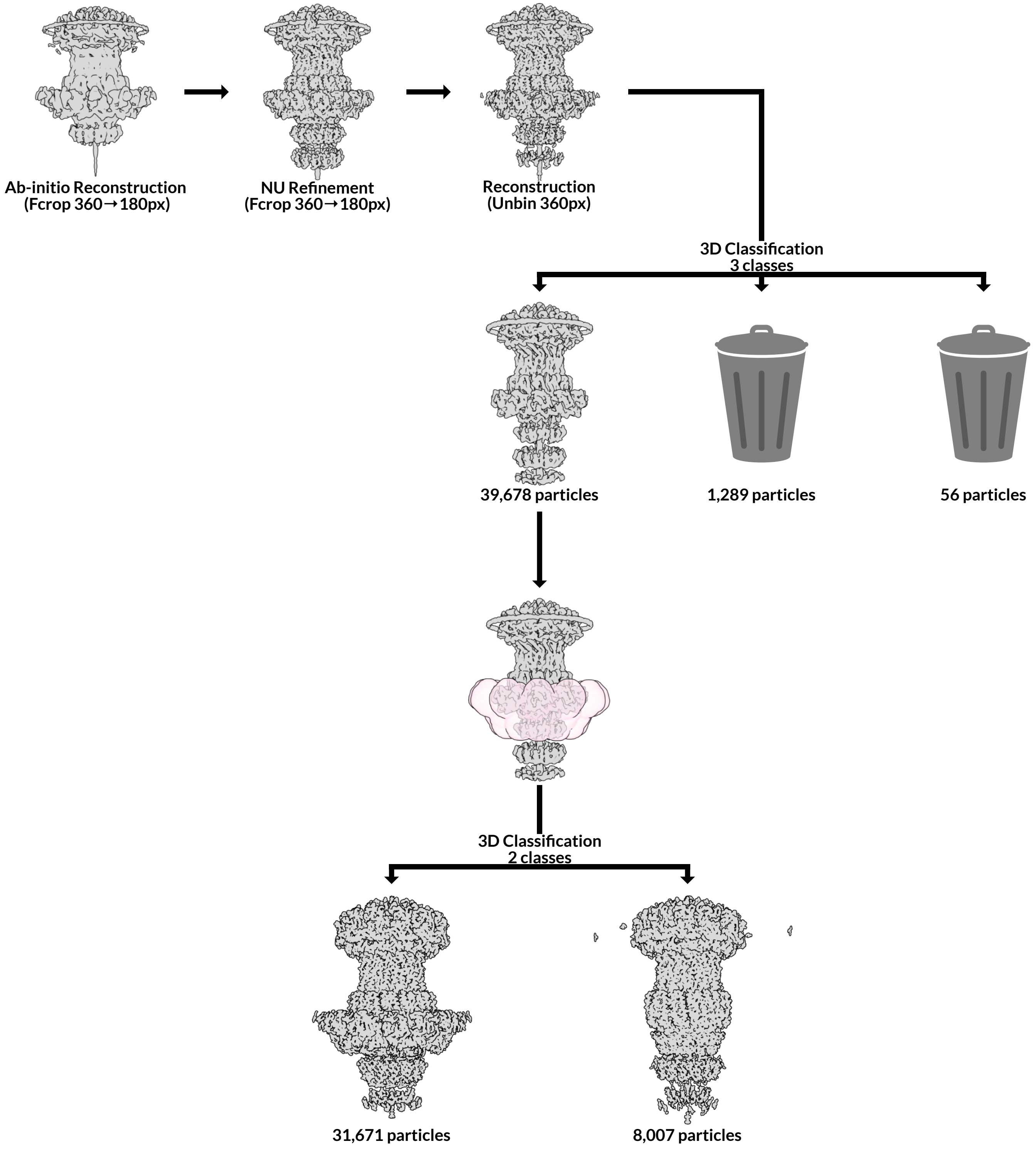


**Figure S3: Reconstruction of C12 neck maps reveals a sub-set of particles with no collar.** We refined the C12 *ab-initio* map generated in Figure S2 using CryoSPARC Non-Uniform (NU) Refinement, then un-binned and reconstructed the particle stack. We used one round of unmasked 3D classification with 3 classes to remove 1,345 junk particles from 2 classes, leaving behind one large class with 39,678 particles. We 3D classified the remaining particle stack into 2 classes using a focus mask around the collar region (pink). One class had density in the collar region (left), while the other did not (right). Both classes are shown after NU-refinement with C12 symmetry, followed by global and local CTF refinement, and a final round of NU-refinement with C12 symmetry. See Figure S5 and Table S5 for resolution and comparison of these maps.

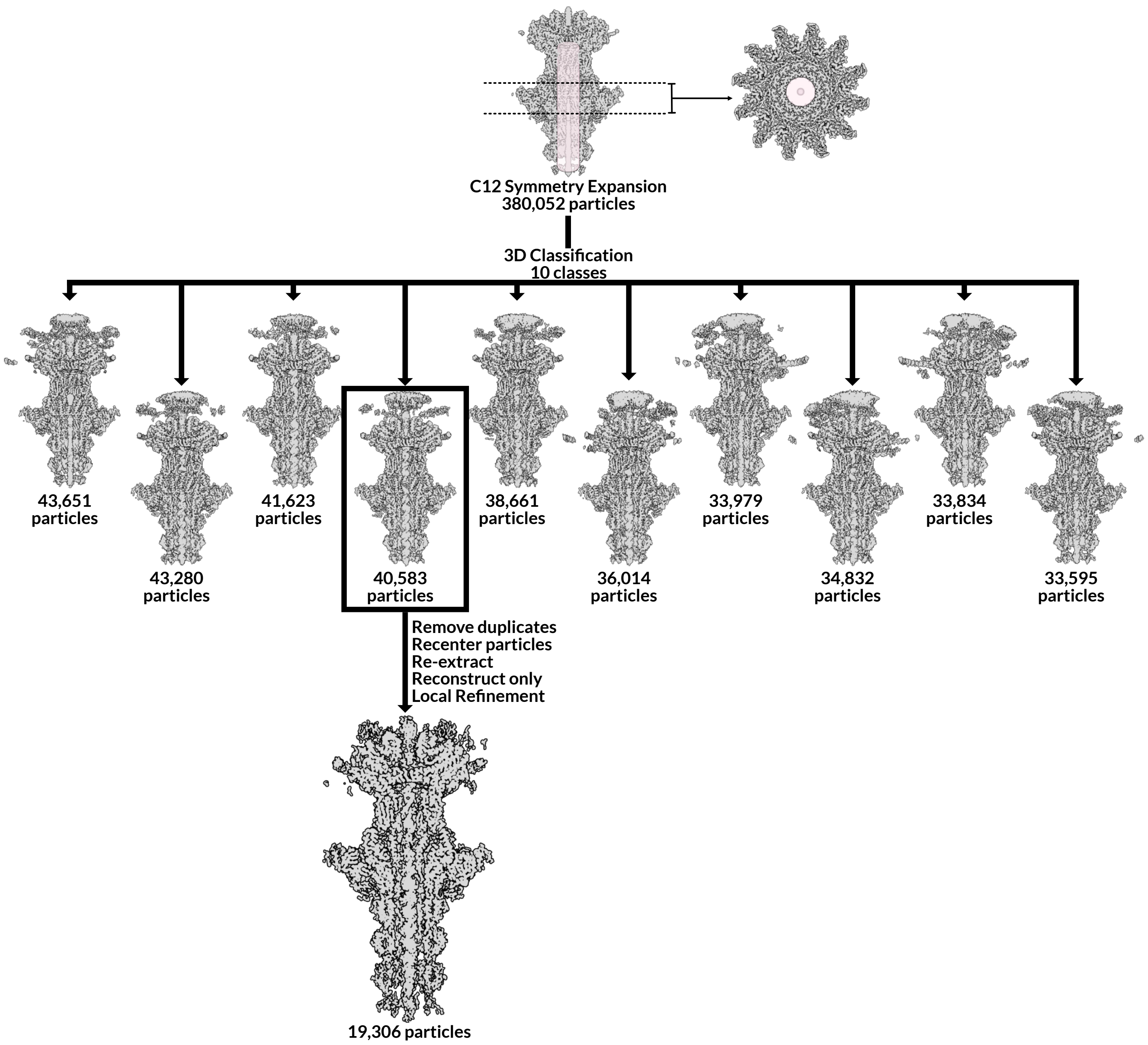


**Figure S4: Symmetry expansion and 3D classification reveals DNA in the central channel of the neck.** We symmetry expanded the C12 reconstruction generated in Figure S3 around the C12 space group. We generated a cylindrical mask, pink, covering the region of the central channel below the portal pore loops. We performed masked 3D classification with 10 classes . Double-stranded DNA is visible in many of these classes, and we selected the one with the clearest double helix for further refinement. We removed duplicate particles and re-extracted the remaining particles with the center of the box moved below the portal pore loops, since including this region negatively impacted alignment and resolution. We then reconstructed these particles and used Local Refinement to generate the final map. See Figure S5 and Table S4 for resolution of this map and statistics of the model built into it.

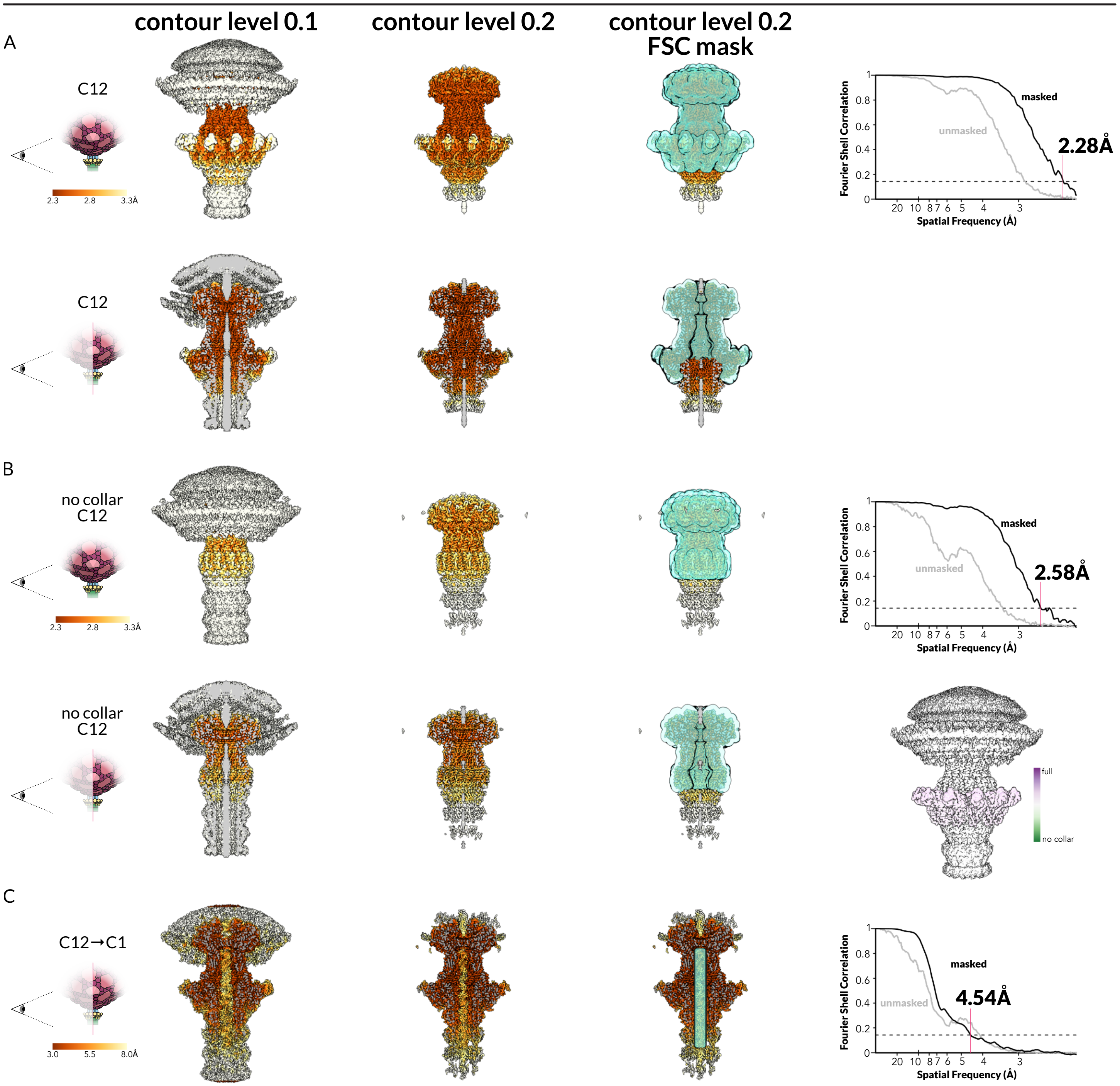


**Figure S5: Neck maps with C12 symmetry reach 2.3-Å and 2.6-Å global resolution. A. Local resolution of the C12 neck map at two different contour levels.** The map generated in figure S3 (left) is colored with high resolution regions in brown and low resolution regions in yellow and white. In turquoise, right, a mask generated to exclude C3 and C5 regions. Bottom row: the neck complex is cut to show the inside of the complex. Far right: GSFSC curve of the C12 neck map. Grey line: unmasked GSFSC. Black line: GSFSC inside the turquoise mask. Dotted line: 0.143 threshold. **B. Local resolution of the collar-less C12 neck map at two different contour levels.** The map generated in figure S3 (right) is colored and masked as in A. Top far right: GSFSC curve of the C12 neck map. Grey line: unmasked GSFSC. Black line: GSFSC inside the turquoise mask. Dotted line: 0.143 threshold. Bottom far right: the complete C12 neck map (A) colored according to similarity to the collar-less neck map in B. White regions of the map have equal density in both maps, while regions in purple have more density in the complete map and regions in green have more density in the collar-less map. **C. Local resolution of the central channel DNA density in a C1 neck map at two different contour levels.** The map generated in figure S4 is colored as in A. In turquoise, right, a mask generated to include only the region of the map that we built DNA into. Far right: GSFSC curve of the central DNA map. Grey line: unmasked GSFSC. Black line: GSFSC inside the turquoise mask. Dotted line: 0.143 threshold.

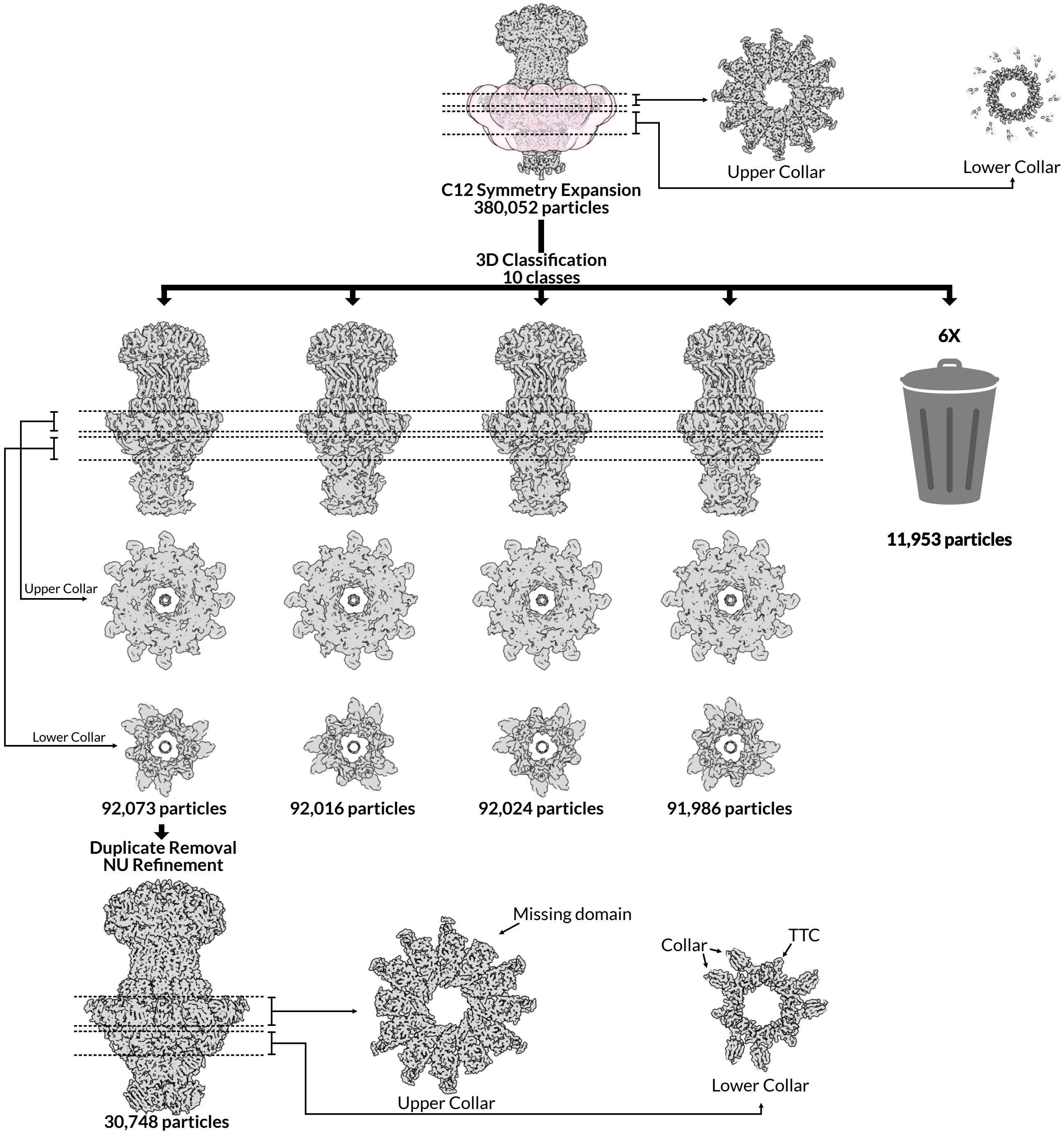


**Figure S6: Breaking the symmetry of the C12 neck shows C3 and C6 symmetry.** We examined the C12 reconstruction with full collar region generated in figure S3. Slices through the upper collar region (right) and lower collar region (far left) show that most of the upper collar region has C12 symmetry, although density in the distal domains is poor and the innermost ring is not resolved. The lower collar region has poor density with C12 symmetry enforced. We applied C12 symmetry expansion and then used 3D classification focused on the pink mask around the collar region to separate four classes of the neck from 6 junk classes. Each of the neck classes is identical except for the collar region, which showed the same C3 symmetry rotated around the central axis. Every fourth domain of the upper collar region is unresolved, likely unstructured. The inner portion of the upper collar ring appears to have 6-fold symmetry. In the lower collar, three pairs of collar domains are located at thirds around the barrel of the neck. The TTP ring also has C3 symmetry, which is visible in the gaps between the lower collar. The largest neck class was chosen for further processing. After duplicate removal, it was refined with NU-refinement and C3 symmetry enforced. See Figure S8A for resolution of this map.

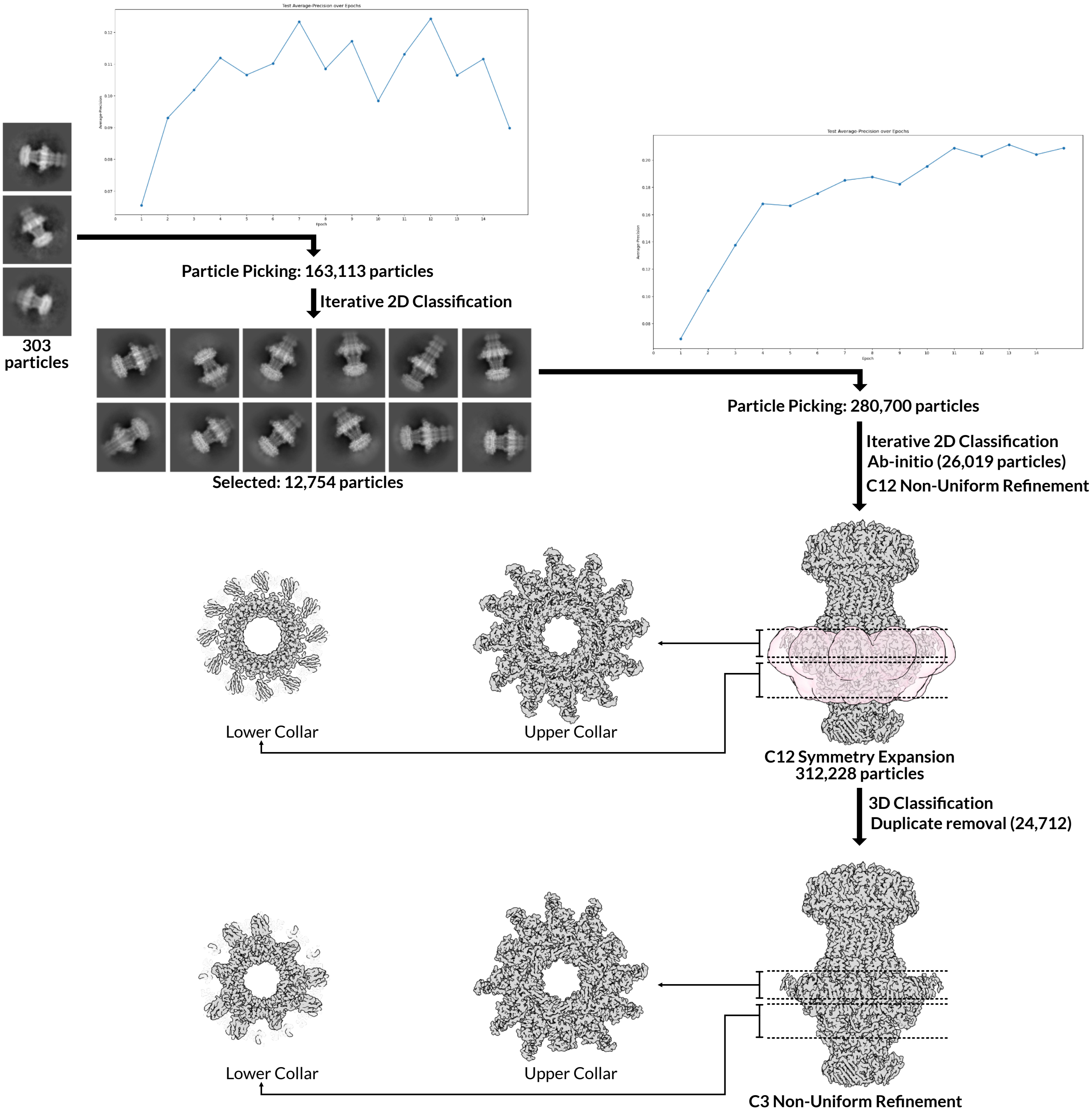


**Figure S7: A subset of neck particles have no capsid attached.** We observed particles that resembled the neck attached to tail filaments, but free of capsids. We manually picked 303 such particles and used them to train a convolutional neural network through the Topaz wrapper in CryoSPARC. We 2D classified the particles picked by this model and used selected particles were used to train a second neural network. After 2D classificatiion, we used selected particles to reconstruct and refine a volume with C12 symmetry. We symmetry expanded this particle stack around the C12 space group and 3D classified it with a focus mask over the collar region, as in Figure S4. We selected the largest class for further processing. After removal of duplicate particles, we refined 24,712 unique particles with C3 symmetry. As with the complete neck in Figure S4, the upper and lower collar both have C3 symmetry, while the innermost portion of the lower collar has C6 symmetry. See Figure S7B for resolution of this map.

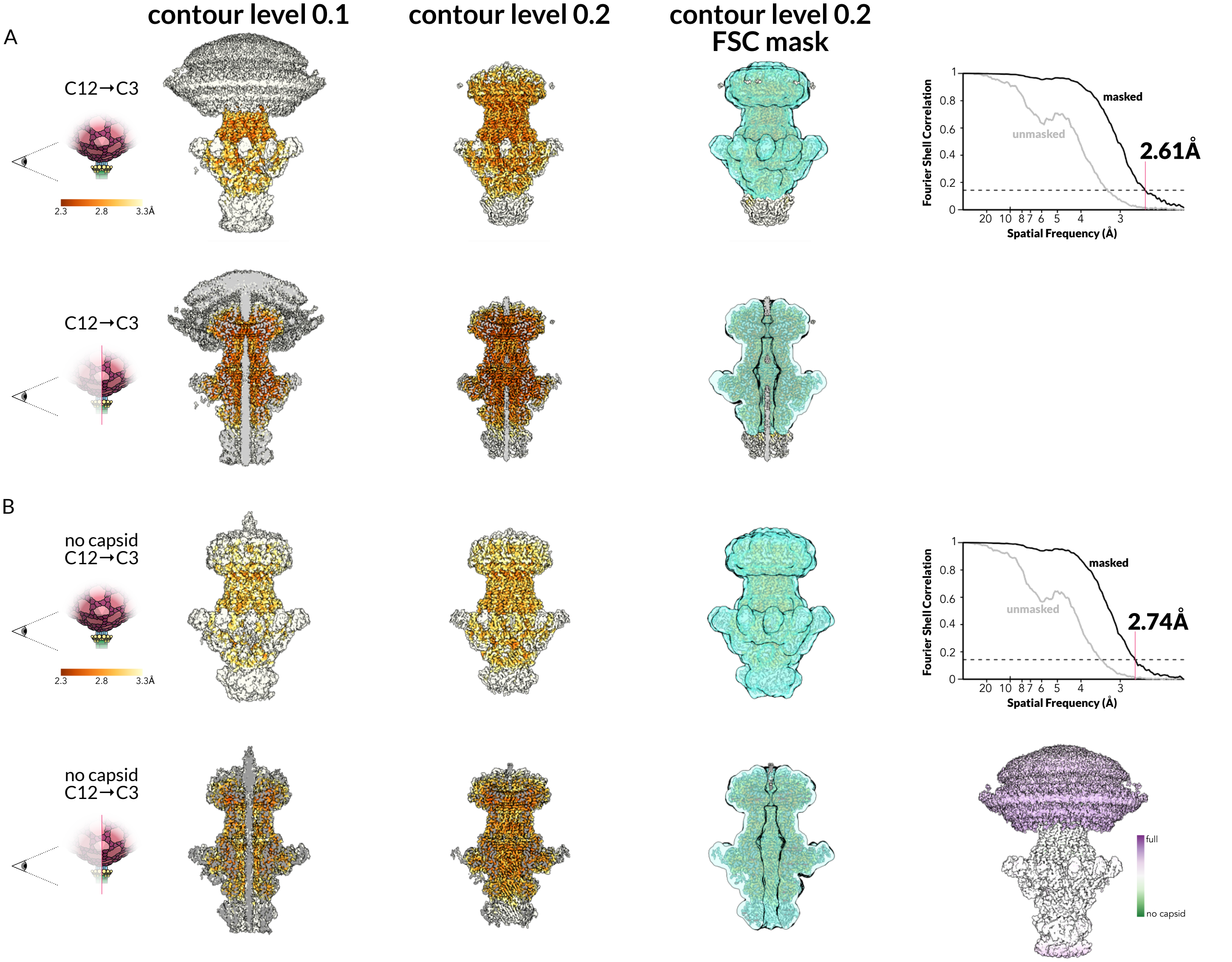


**Figure S8: Maps with C3 symmetry reach 2.6-Å and 2.7-Å global resolution, respectively. A. Local resolution of the symmetry broken C3 neck map at two different contour levels.** The map generated in figure S6 is colored with high resolution regions in brown and low resolution regions in yellow and white. In turquoise, right, a mask was generated to exclude C5 regions. Bottom row: the neck complex is cut to show the inside of the complex. Far right: GSFSC curve of the C3 neck map. Grey line: unmasked GSFSC. Black line: GSFSC inside the turquoise mask. Dotted line: 0.143 threshold. **B. Local resolution of the symmetry broken capsid-less C3 neck map at two different contour levels.** The map generated in figure S7 is colored and masked as in A. Top far right: GSFSC curve of the capsid-less C3 neck map. Grey line: unmasked GSFSC. Black line: GSFSC inside the turquoise mask. Dotted line: 0.143 threshold. Bottom far right: the complete C3 neck map (A) colored according to similarity to the capsid-less neck map in B. White regions of the map have equal density in both maps, while regions in purple have more density in the complete map and regions in green have more density in the capsid-less map.

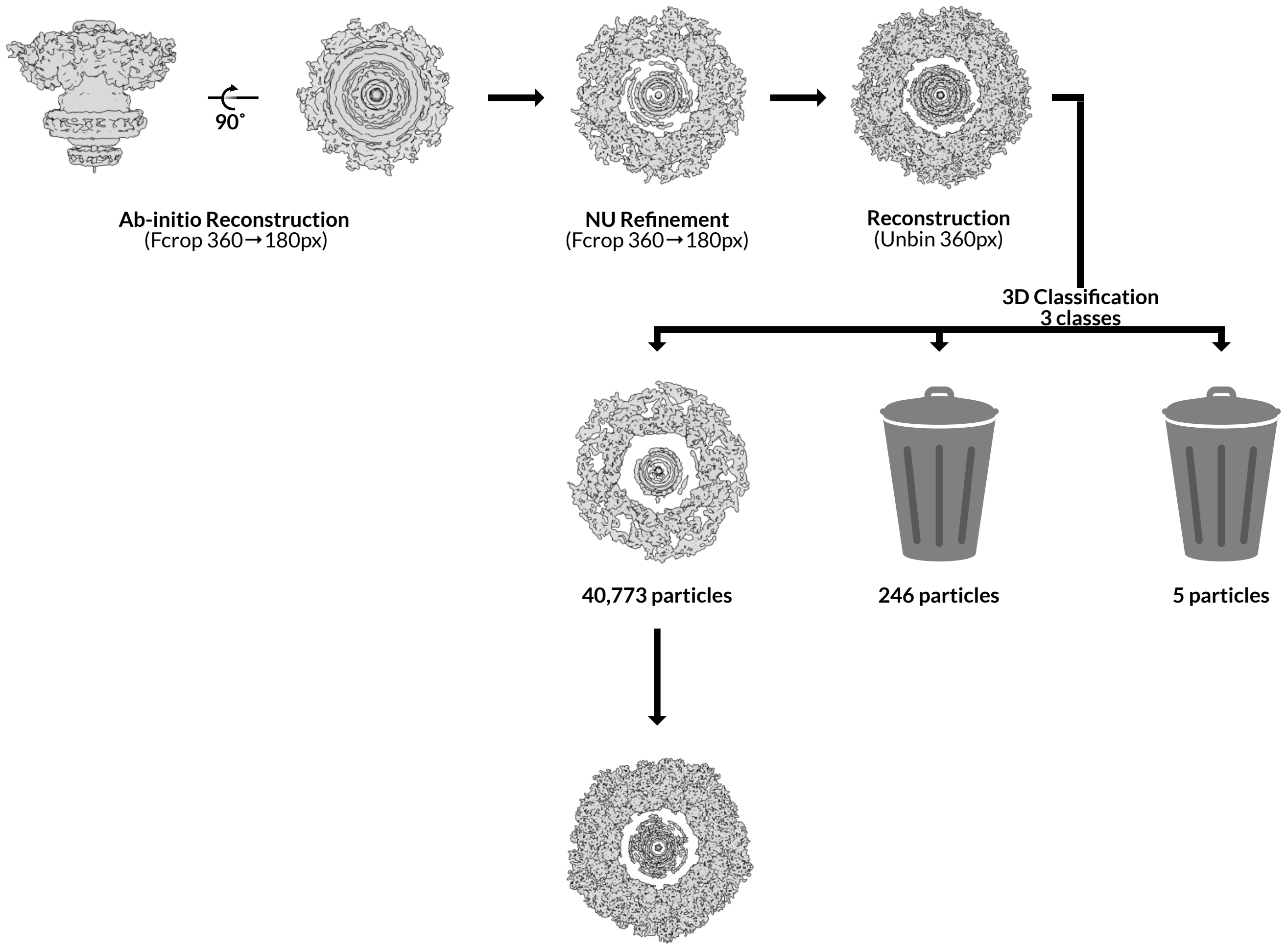


**Figure S9: Reconstruction of the C5 HTCC map reveals the unique portal vertex.** We refinef the C5 *ab-initio* map generated in Figure S2 using CryoSPARC Non-Uniform (NU) Refinement, and un-binned and reconstructed the particle stack. We performed one round of unmasked 3D classification with 3 classes to remove 251 junk particles from 2 classes, leaving behind one large class with 40,773 particles. We NU-refined this class with C5 symmetry, followed by global and local CTF refinement, and a final round of NU-refinement with C5 symmetry. See Figure S10 for resolution of this map.

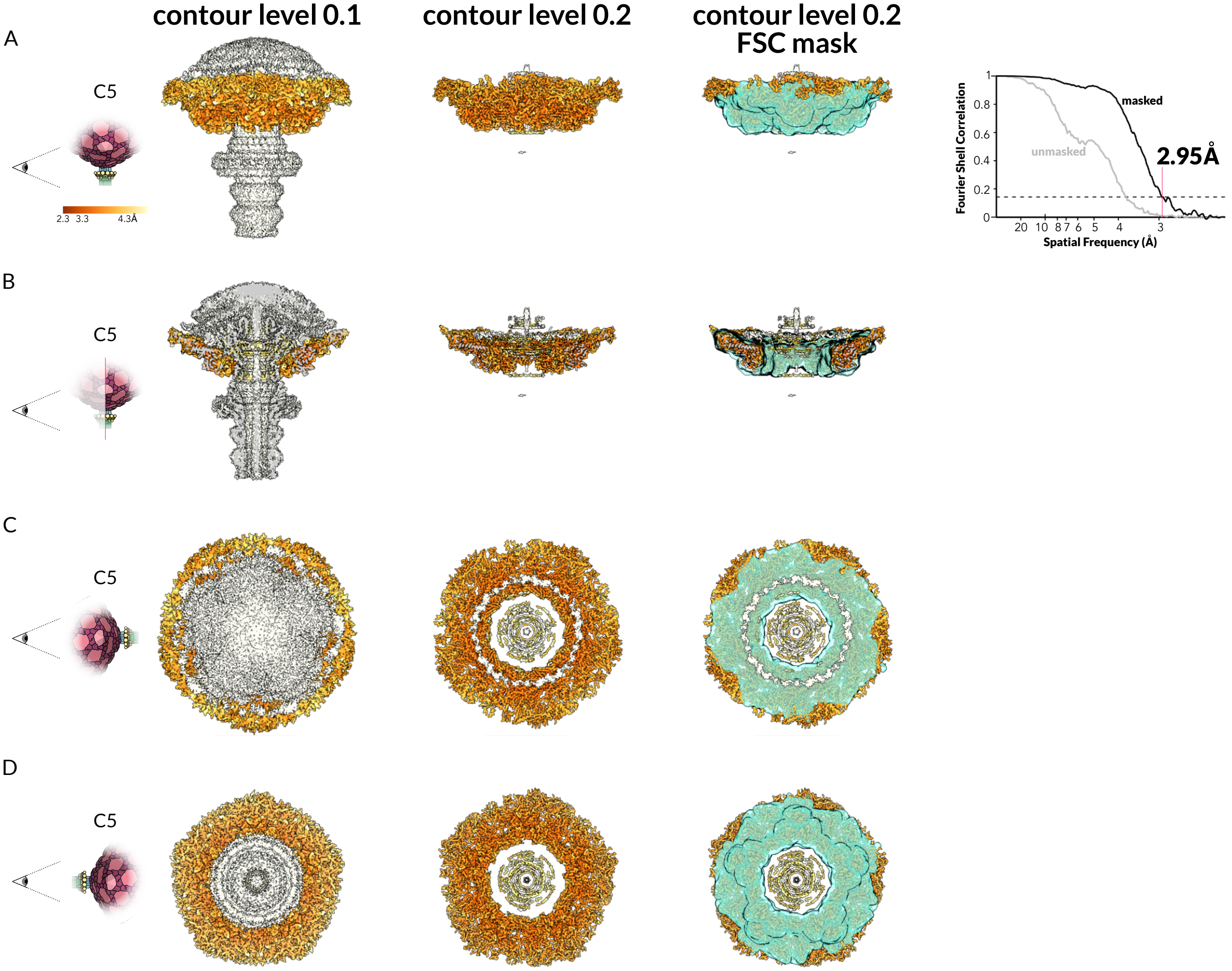


**Figure S10: Neck map with C5 symmetry reaches 3.0-Å global resolution. A. Local resolution of the C12 neck map at two different contour levels.** The map generated in figure S9 is colored with high resolution regions in brown and low resolution regions in yellow and white. In turquoise, right, we generated a mask to cover only C5 regions of the map, excluding C12 and C3 regions. Far right: global GSFSC curve of the C5 neck map. Grey line: unmasked GSFSC. Black line: GSFSC inside the turquoise mask. Dotted line: 0.143 threshold. **B. Local resolution of the C5 neck map and mask cut to show in inside of the complex.** Colors are as in A. **C. Local resolution of the C5 neck map and mask from inside the capsid.** Colors are as in A. **D. Local resolution of the C5 neck map and mask from outside the capsid.** Colors are as in A.

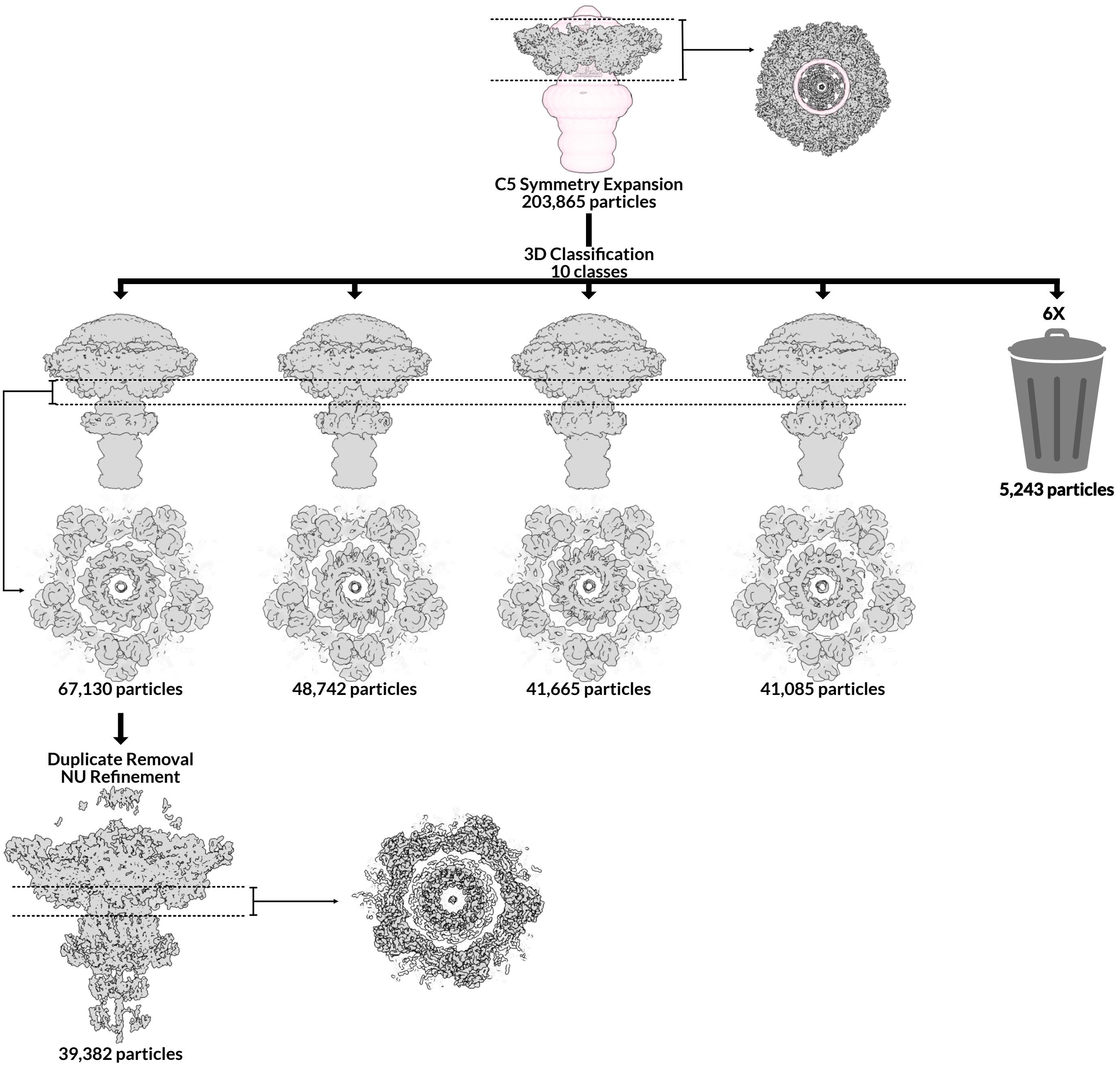


**Figure S11: Breaking the symmetry of the C5 neck shows interactions with the C12 portal.** We symmetry expanded the C5 reconstruction region generated in figure S9 around the C5 space group. 3D classification focused on the pink mask around the neck region separated four classes of the neck from 6 junk classes. After symmetry expansion and 3D classification, both the C5 capsid and C12 portal are clearly visible. Each of the neck classes is identical except that the portal is rotated with respect to the capsid. The largest neck class was chosen for further processing. After duplicate removal, it was asymmetrically NU-refined. See Figure S11 for resolution of this map.

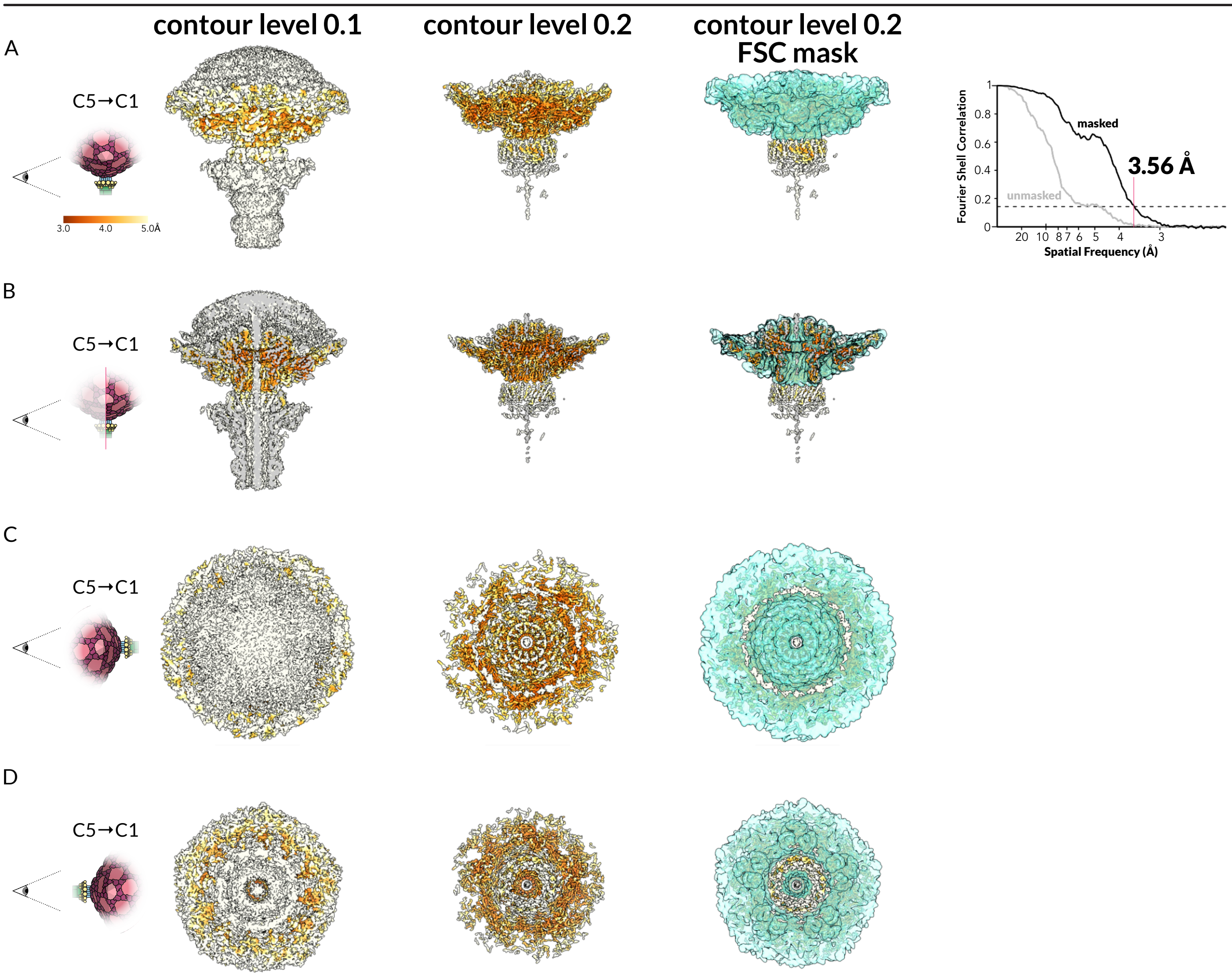


**Figure S12: Neck map with symmetry broken to show both C5 and C12 components reaches 3.6Å global resolution.** A. Local resolution of the asymmetric neck map **at two different contour levels.** The map generated in figure S11 is colored with high resolution regions in brown and low resolution regions in yellow and white. In turquoise, right, a mask was generated to cover only the portal and capsid regions of the map. Far right: global GSFSC curve of the asymmetric neck map. Grey line: unmasked GSFSC. Black line: GSFSC inside the turquoise mask. Dotted line: 0.143 threshold. **B. Local resolution of the asymmetric neck map and mask cut to show in inside of the complex.** Colors are as in A. **C. Local resolution of the asymmetric neck map and mask from inside the capsid.** Colors are as in A. **D. Local resolution of the asymmetric neck map and mask from outside the capsid.** Colors are as in A.

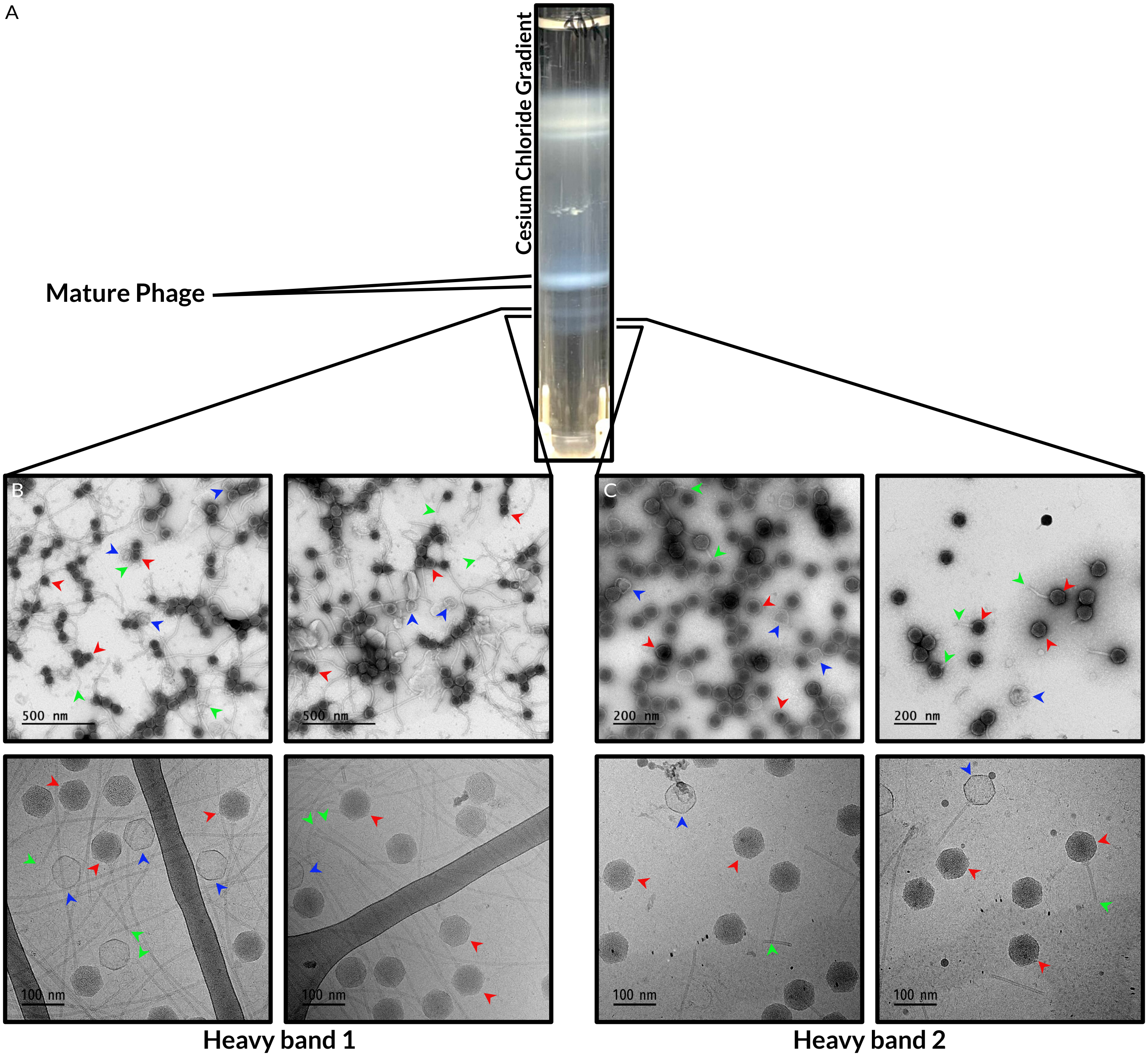


**Figure S13: Virions with broken tails retain DNA in their capsids. A. Cesium Chloride gradient with a large band of mature virions and two minor dense bands.** We applied PEG-precipitated virions to the top of a Cesium Chloride step gradient and centrifuged for 3h in a Beckman SW40-Ti rotor at 30,000 RPM. We collected the resulting bands with a long needle. **B. Negative stain (top row) and cryogenic (bottom row) electron micrographs of the higher dense band.** After collection, we concentrated the higher dense band and exchanged it into Cesium-free buffer, then applied it to grids. Red arrows highlight examples of DNA-filled capsids, while blue arrows highlight examples of empty capsids. Green arrows highlight examples of the ends of broken tails. **C. Negative stain (top row) and cryogenic (bottom row) electron micrographs of the lower dense band.** After collection, we concentrated the higher dense band and exchanged it into Cesium-free buffer, then applied it to grids. Red arrows highlight examples of DNA-filled capsids, while blue arrows highlight examples of empty capsids. Green arrows highlight examples of the ends of broken tails.

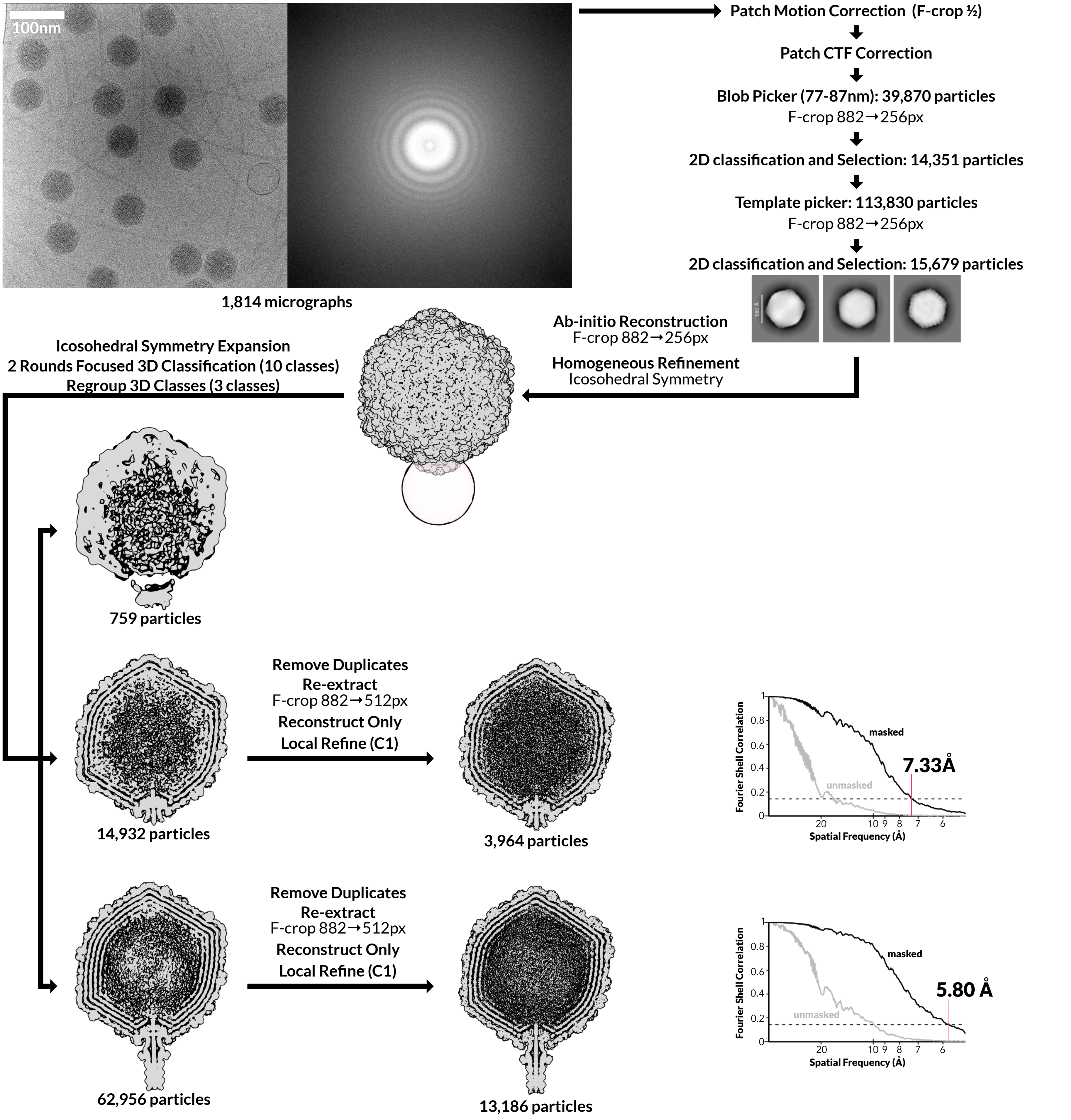


**Figure S14: Phage particles with broken tails still contain DNA.** We applied purified virions to lacy carbon grids and imaged them on the Titan Krios (Methods). One representative micrograph, with its Fourier transform, of a total 1,814 collected micrographs of phage with broken tails (heavy band 1, figure S13), is shown. We performed Patch Motion Correction and Patch CTF correction and used Blob Picker and Template picker to pick capsid particles in CryoSPARC . We used 2D classification to select 38,732 particles representing filled capsids. First, we performed an asymmetric *ab-initio* reconstruction on the particle stack in B after cropping each particle in Fourier space (F-cropping) from a box size of 882 pixels to 256 pixels to reduce computational time and memory usage. We then refined this *ab-initio* map with icosahedral symmetry enforced, and performed symmetry expansion around the same space group. We placed a spherical mask (pink) over a single vertex of the capsid, and used focused 3D classification to select particles with density for the neck at this vertex. We performed 2 rounds of iterative 3D classification using the same focus mask and 10 classes. We regrouped resulting classes into 3 super classes, removed duplicate particles, and reconstructed the particle sets.

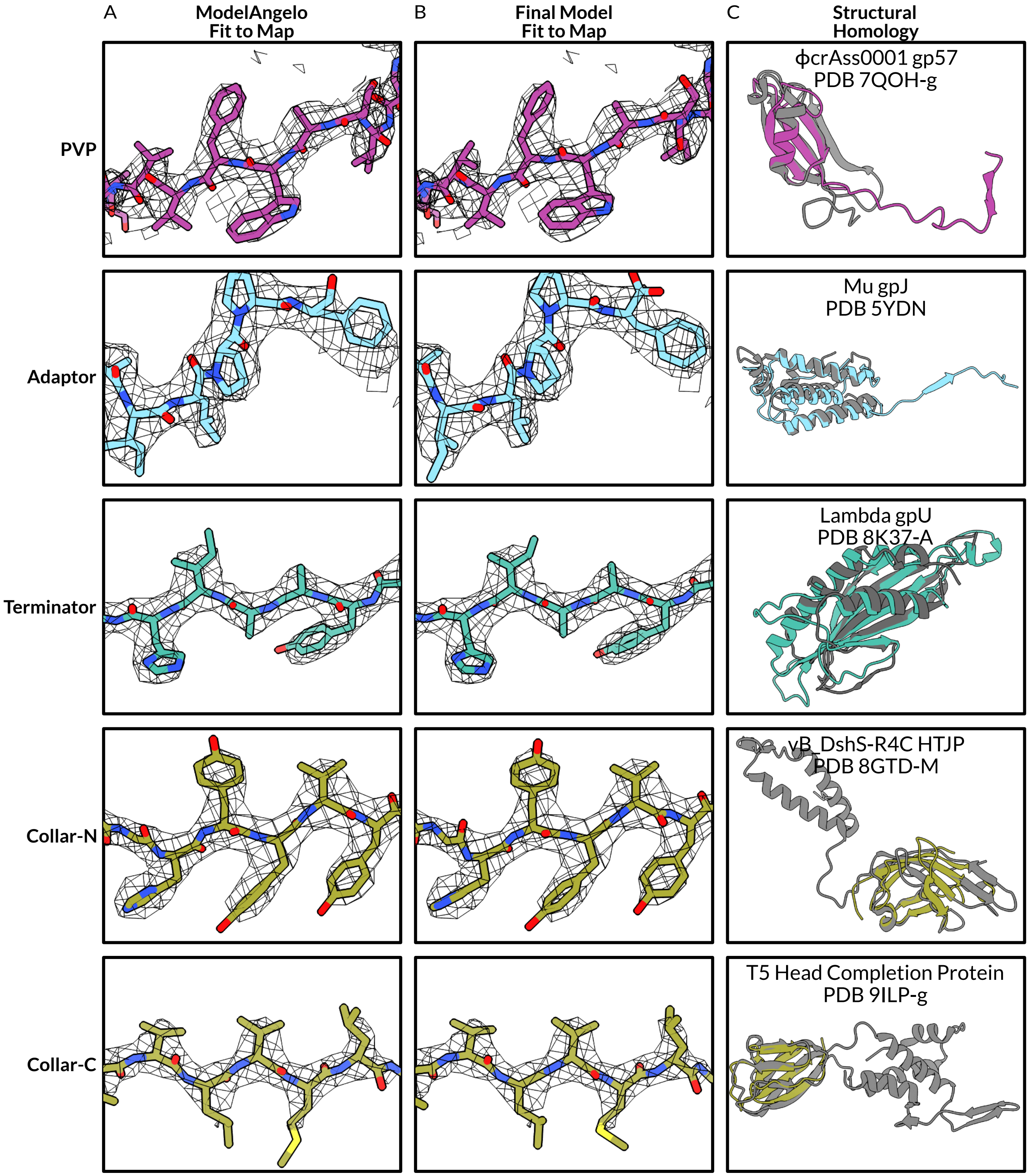


**Figure S15: Computational protein identifications were corroborated by model-building and structural homology. A. Representative sections of the ModelAngelo output in EM density.** We ran ModelAngelo without sequence input on the C5 (PVP) and C3 (adaptor, terminator, collar) maps. **B. Representative sections of the final model in EM density.** After automated building with sequence input, we built models in Coot and Isolde and refined in Phenix. **C. Overlay of the final model with its structural homolog.** We identified structural homologs using the DALI server, then selected a representative homolog and aligned it to the P74-26 protein in ChimeraX.

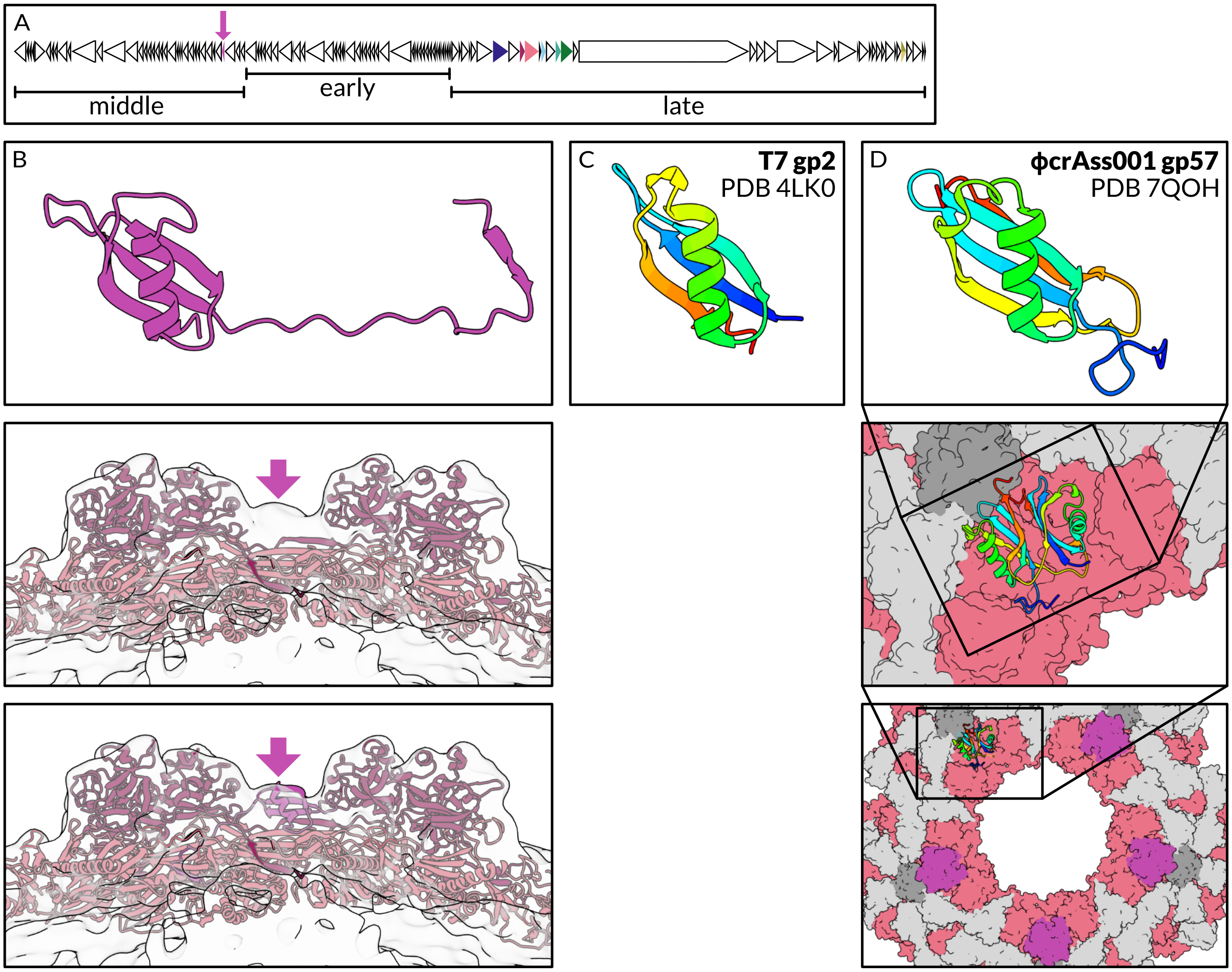


**Figure S16: The Portal Vertex Protein shares homology with a phage-encoded RNA Polymerase inhibitor. A. Linear map of the P74-26 genome with each ORF displayed as a triangle proportional to the length of the ORF.** Genes are colored as in Figure 1 and the location of PVP is highlighted with a large purple arrow. Underneath, bars indicate which genes are expressed early, middle, and late in P74-26 (Berdygulova et al. *Journal of Molecular Biology.* 2011.). **B. Final atomic model of PVP, and fit of capsid proteins into the C5 map of the expanded empty capsid.** *Top:* Final atomic model of PVP in purple. Middle: Map of the C5 expanded empty capsid (EMDB 4446) with the atomic models of the mature MCP and Dec fit into the density. Purple arrow indicates the un-modeled density. *Bottom:* The C5 expanded empty capsid with the atomic models of the mature MCP, Dec, and PCP fit into the density. The location of PVP is highlighted with a purple arrow. **C. PVP homolog gp2 of T7.** gp2 (PDB 4LKO) is colored in rainbow from N-terminus (blue) to C-terminus (red). **D. PVP homolog gp57 of ϕcrAss001.** *Top:* Chain g of PDB 7QOH depicted in rainbow. *Middle:* The 7QOH assembly depicted as a surface with MCP in salmon and auxiliary capsid proteins (not homologous to Dec or other P74-26 proteins) in greys. gp57 dimer depicted as rainbow cartoons. *Bottom:* Location of gp57 in the portal vertex. One pair of gp57 subunits depicted as rainbow cartoons, while the other 4 are depicted as purple surfaces.

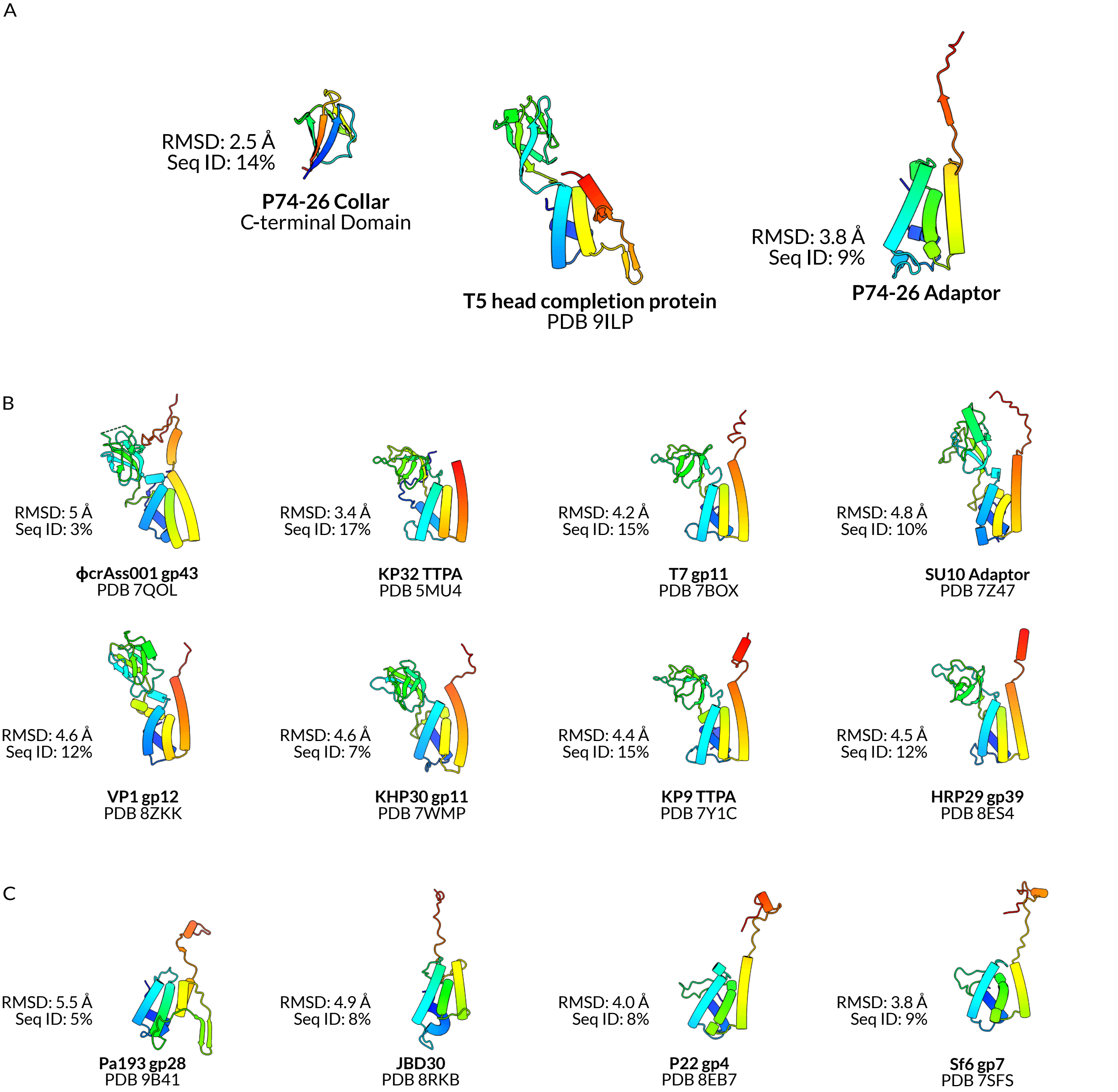


**Figure S17: The adaptor C-terminal tail is conserved in some other phages, while some phages also have an Ig domain incorporated into adaptor. A. The T5 head completion protein with the the collar protein C-terminal domain and the adaptor protein.** The DALI server identified homology between the T5 head completion protein (PDB 9ILP) and both P74-26 proteins. All models are depicted in rainbow from N-terminus (blue) to C-terminus (red). The Ig domain may mediate interaction with the portal Clip region in T5. **B. Structural homologs of adaptor that also contain an Ig domain homologous to the collar C-terminal domain.** The DALI server identified homologous adaptors. The molecular models of eight such adaptors, all belonging to podoviruses, are shown. With the exception of KP32, all of these adaptors also include a C-terminal tail. The Ig domains are commonly used to bind tail fibers. **C. Structural homologs of adaptor that include a C-terminal tail, but no Ig domain.** Homologs were identified by the DALI server. Unlike the Ig domains, C-terminal tails are spread throughout myoviruses (Pa193), siphoviruses (JBD30), and podoviruses (P22, Sf6). Some adaptors, like Mu gpJ (Fig S12B, PDB 5YDN), have neither an Ig domain or a C-terminal tail, while others have the addition of different functional domains.

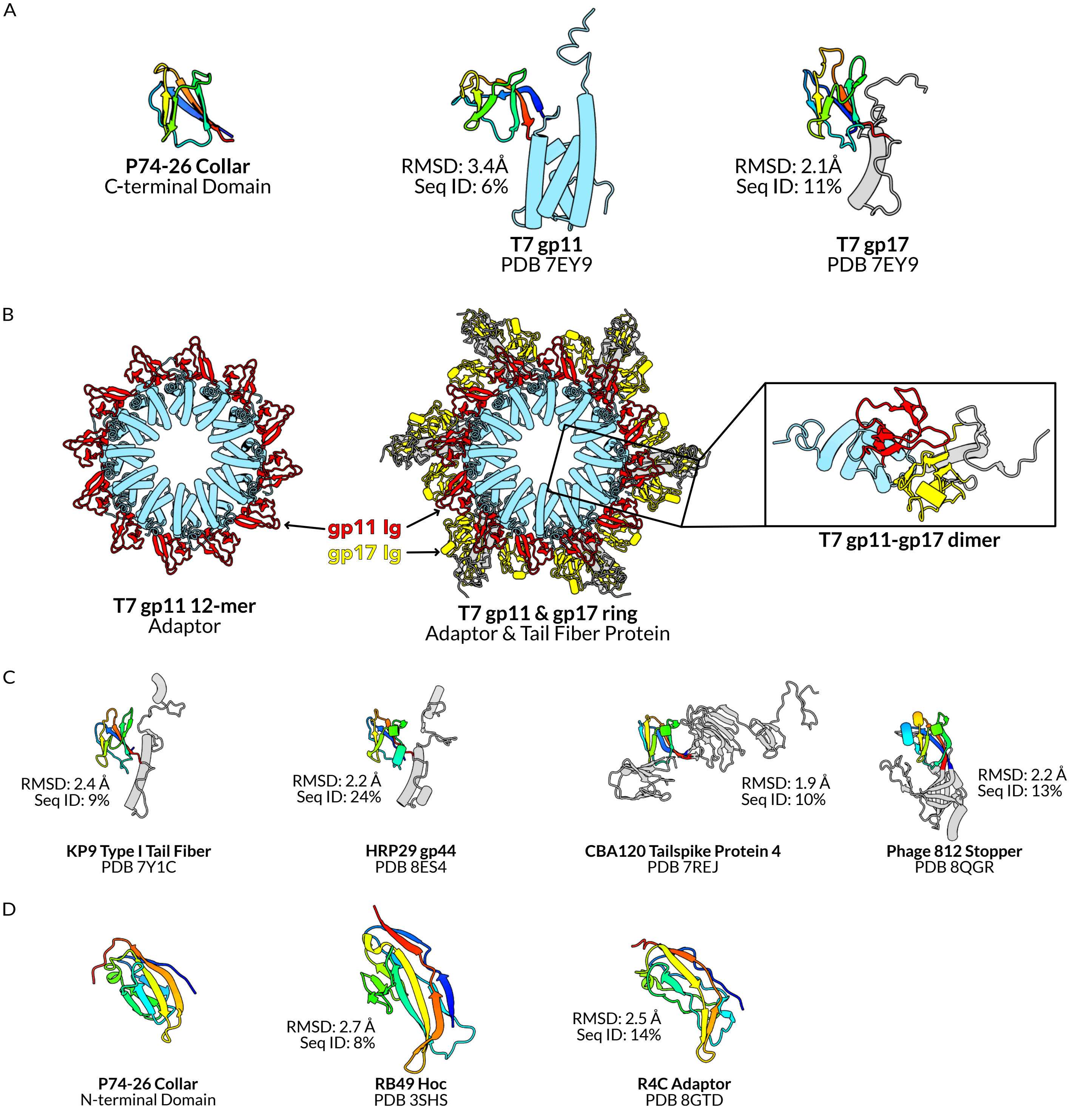


**Figure S18: The Immunoglobulin domains of collar protein are homologous to both tail fiber proteins and other neck proteins. A. The C-terminal domain of collar protein with the adaptor (gp11) and Tail Fiber Protein (gp17) of T7.** *Left:* Residues 88-146 of collar protein are colored sequentially from N-terminal (blue) to C-terminal (red). *Center:* Regions of the T7 adaptor protein (gp11) that bear homology to the adaptor protein of P74-26 are colored cyan. Residues 62-125 are colored sequentially. *Left:* Residues 6-80 of T7 TFP are colored sequentially. Note that residues 1-5 and 142-553 of gp17 were not modeled. **B. The T7 adaptor ring in isolation, and with gp17.** *Left*: 12 adaptor subunits form a ring, with their Ig domains (red) decorating the outer surface of the tube. *Center:* Six trimers of gp17 bind to the outer surface of the adaptor ring with their Ig domains (yellow) contacting the Ig domains of adaptor. *Left:* A gp11-gp17 dimer interacting through their Ig domains. **C. Other phage proteins containing this Ig fold.** T7-like phages KP9 and HRP29 also contain a similar Ig fold in their TFPs, as does CBA Tailspike Protein 4 and 812 Stopper protein. **D. The N-terminal domain of collar protein with the highly immunogenic outer capsid protein (Hoc) of a T4-like phage (RB49) and the adaptor of marine siphophage R4C.** Residues 1-85 of collar protein are colored sequentially, as are residues 183-304 of Hoc and 1-104 of R4C adaptor. Residues 1-184 of Hoc and 105-178 of R4C adaptor are hidden for clarity.

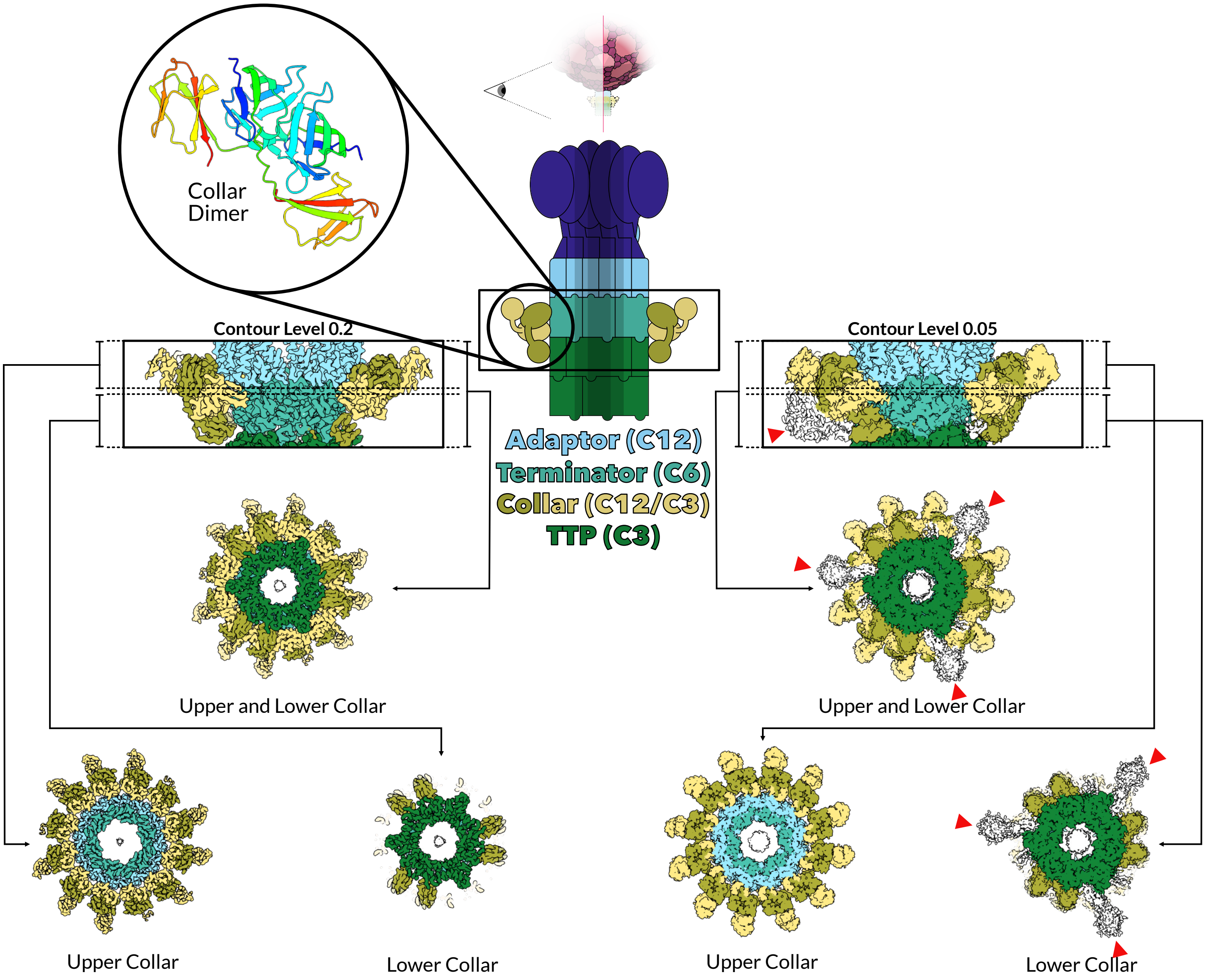


**Figure S19: The collar region has C3 symmetry and some domains may be flexibly tethered or unstructured.** The collar consists of 12 dimers of collar protein, and each collar protein consists of two flexibly tethered globular domains. Two subunits of collar protein interact through their N-terminal domains (circular inset) to form a dimer. In the lower collar, six domains of collar protein are not visible. Additionally, at a conservative contour level (0.2, left), the outermost domain of every fourth lobe of the upper collar is unresolved. At a generous contour level (0.5, right), all 12 lobes of the upper collar are visible, and there is indistinct density below every fourth lobe (red arrow heads).

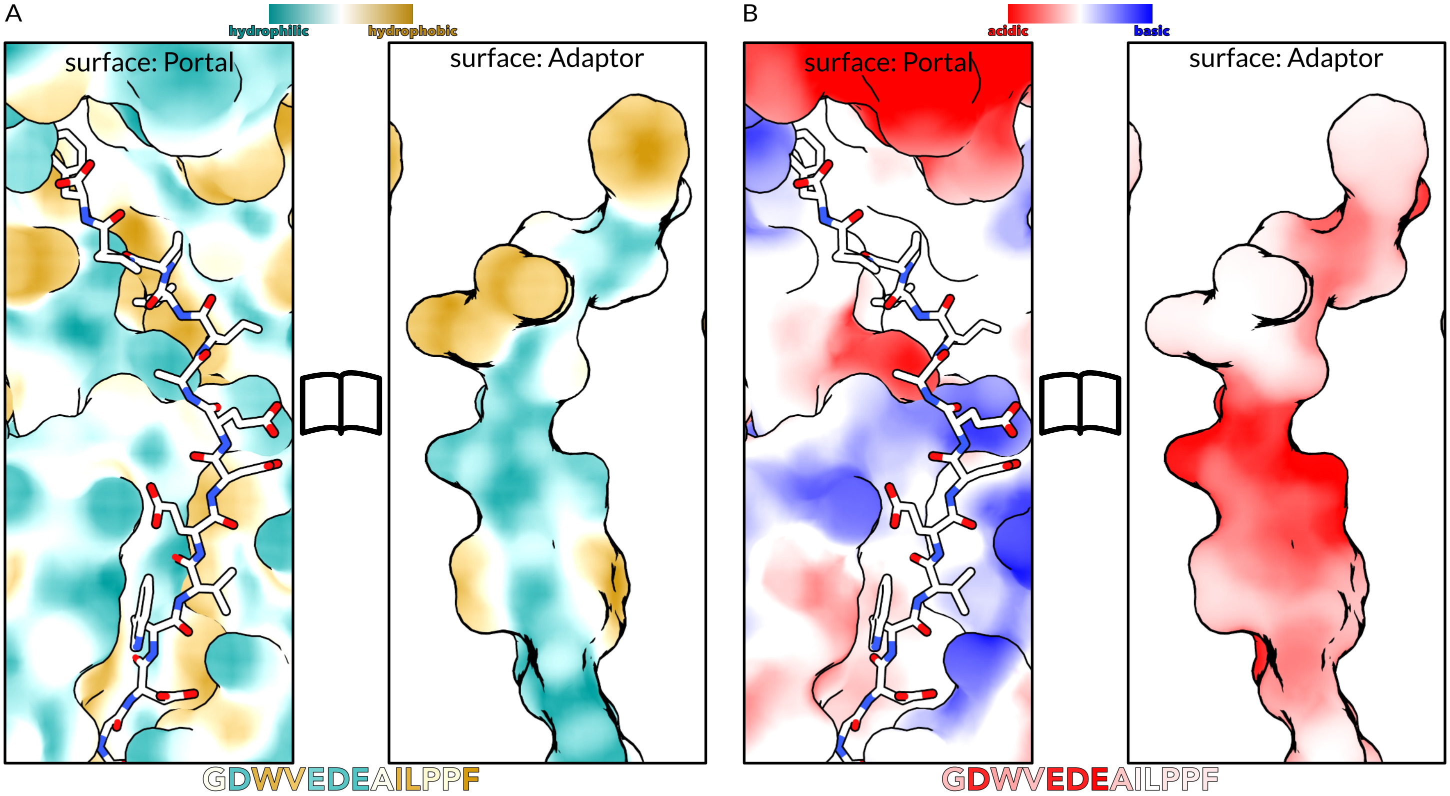


**Figure S20: The adaptor C-terminal tail and the portal clip are secured through both hydrophobic and electrostatic interactions. A. EM density of the neck complex with portal in rainbow, reproduced from Figure 2E.** Adaptor and collar proteins are colored as in Figure 1. Inset shows the atomic model of the adaptor tail (sticks) interacting with the clip region of three different portal subunits (surfaces). **B. Atomic models of the portal-adaptor interaction, colored by hydrophobicity.** *Far left:* Surface model of the portal, colored by hydrophobicity, with the adaptor tail in white sticks. *Left:* Adaptor C-terminal tail shown as a surface colored by hydrophobicity. **C. Atomic models of the portal-adaptor interaction, colored by coulombic potential.** *Right:* Surface model of the portal, colored by charge, with the adaptor tail in white sticks. *Far right:* Adaptor C-terminal tail shown as a surface colored by charge.

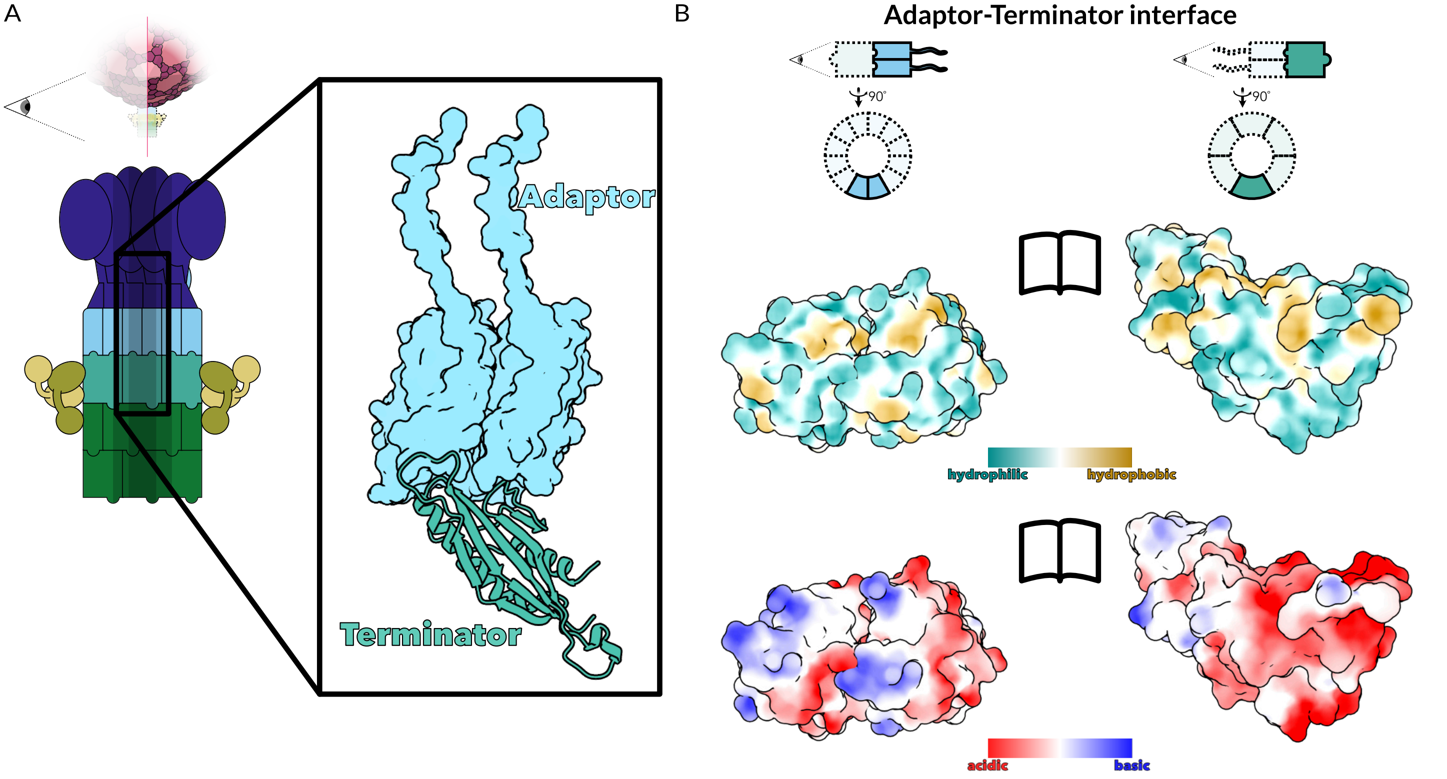


**Figure S21: The adaptor is secured to the terminator through loop-and-socket interactions. A.** Diagram of the neck complex, with enlargement showing the **adaptor-terminator interaction.** Enlargement shows the atomic model of two adaptor subunits (light blue, surface view) and one terminator subunit (teal, cartoon view), as viewed from the inside of the neck channel. **B. The interface between the adaptor and terminator colored by hydrophobicity (top) or coulombic potential (bottom).** Atomic models of two adaptor subunits (left) and one terminator subunit (right) are shown as surfaces.

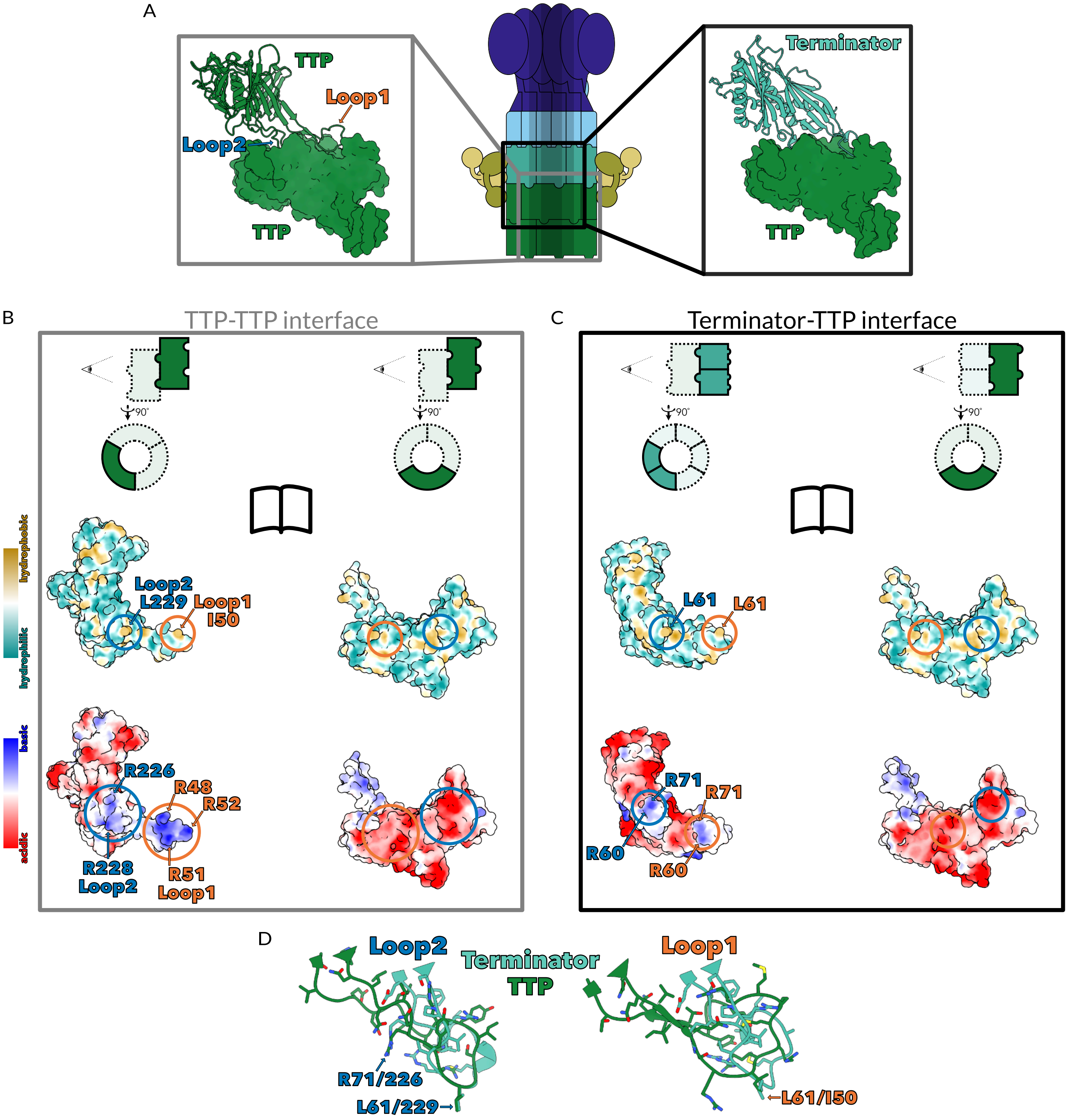


**Figure S22: Terminator connects to Tail Tube Protein using loops that mimic the loop-and-socket connections between rings of TTP. A. Diagram of the neck complex (center), with enlargements showing the TTP-TTP interaction (left) and terminator-TTP interaction (right).** *Left*: One subunit of TTP is shown in cartoon, while the TTP subunit below it is shown as a surface. Loops 1 (blue) and 2 (orange) are highlighted with arrows. *Right*: Two subunits of terminator are shown in cartoon, while the TTP subunit below them is shown as a surface. **B. The interface between two TTP subunits, colored by hydrophobicity (top) or coulombic potential (bottom).** Surfaces of the TTP loops (left) and Sockets (right) shown as surfaces colored by hydrophobicity (top) and coulombic potential (bottom). Residues of the TTP loops making key interactions are indicated with arrows, while the area of the socket surface that makes contact with those residues is circled. **C. The interface between two terminator subunits and one TTP subunit, colored by hydrophobicity (top) or coulombic potential (bottom).** Depictions of terminator loops and TTP sockets are as in B. **D. Structural alignment of TTP and terminator loops.** Cartoons of TTP and terminator loops were superimposed by aligning the TTP sockets they interact with.

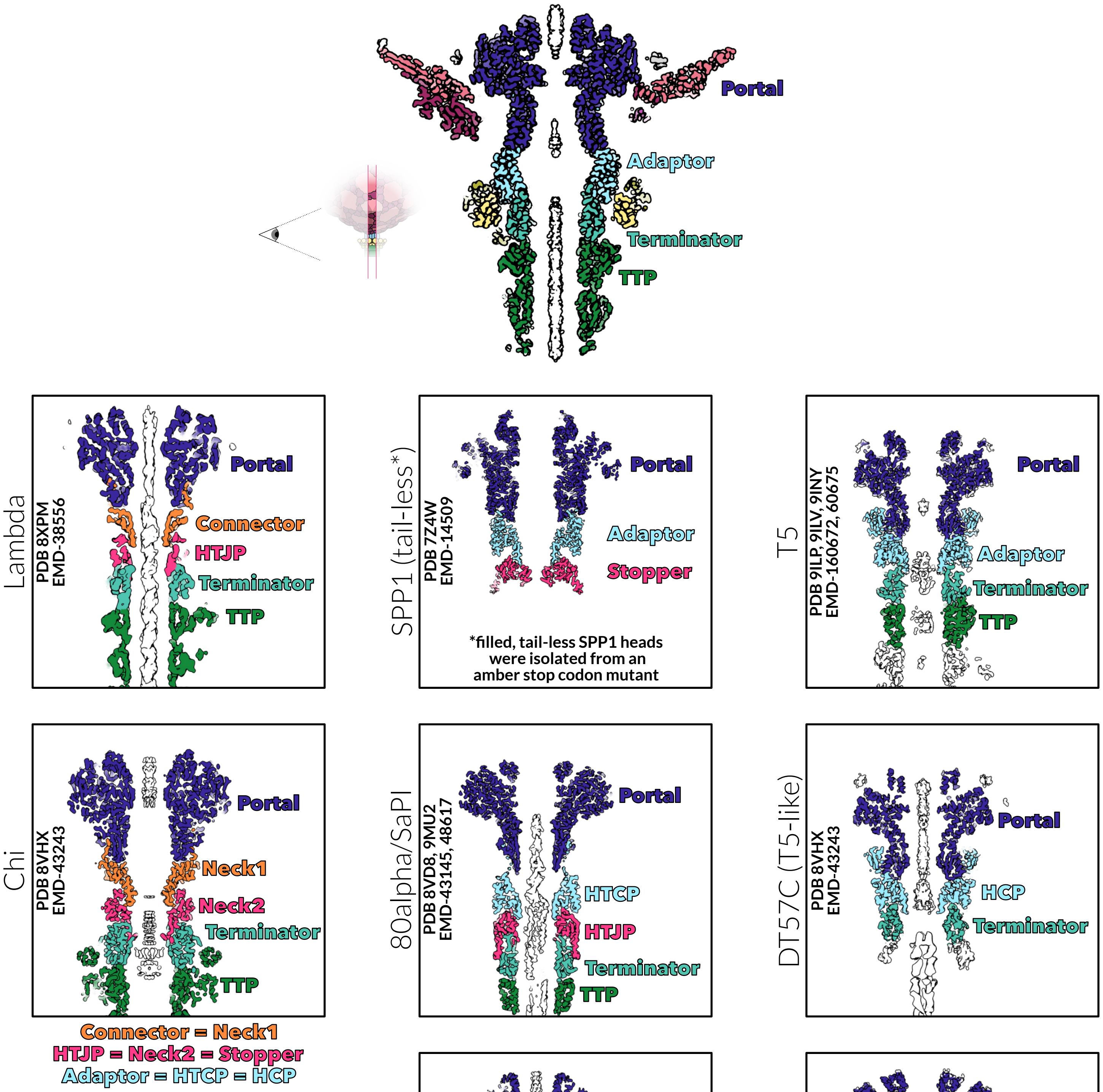


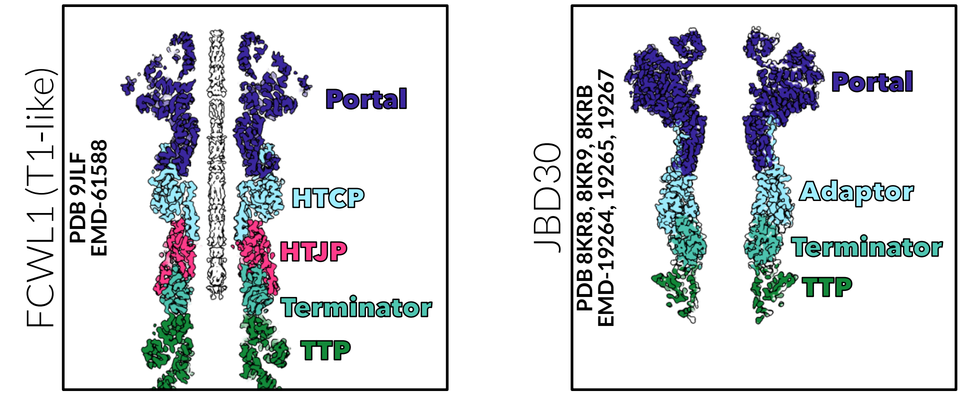


**Figure S23: The P74-26 neck is most similar to those of phage JBD30 and the T5 family.** EM density from reconstructions of P74-26 and 8 other siphophages are shown as central slices along the neck. P74-26 is colored as in Figure 1, while other phages are colored according to structural homology of protein components. While the Lambda connector and P74-26 adaptor fill analogous spaces in the neck, they do not share structural homology.

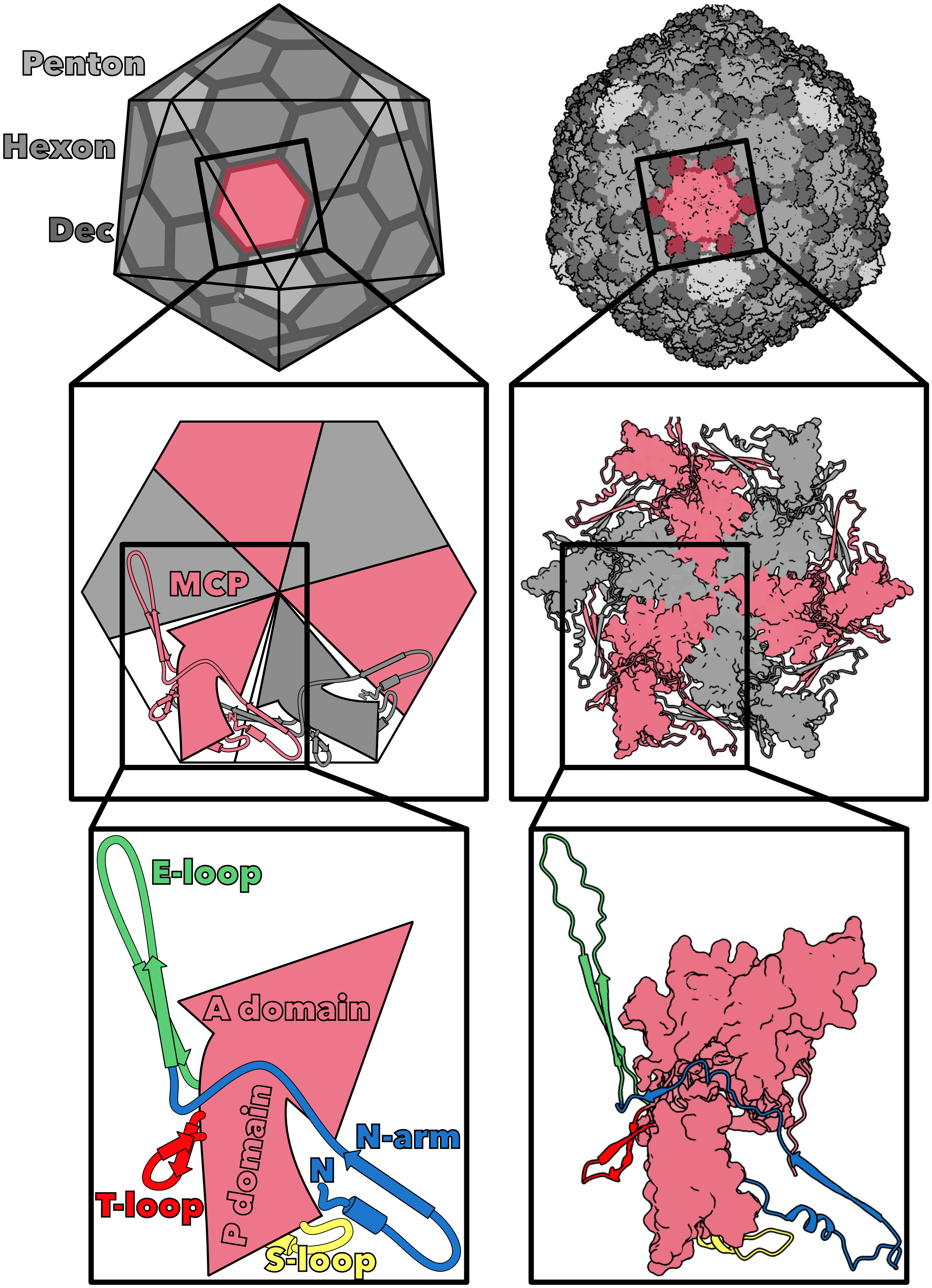


**Figure S24: The icosahedral capsid is composed of MCP subunits.** PDB 6O3H is colored to illustrate how the capsid is composed of hexamers (grey) and pentamers (light grey), how hexamers are composed of MCP subunits, and how MCP subunits contain two major domains and four loops. In the complete capsid, Dec is shown in dark grey and crimson. Hexamers and subunits are viewed as if from inside the capsid.

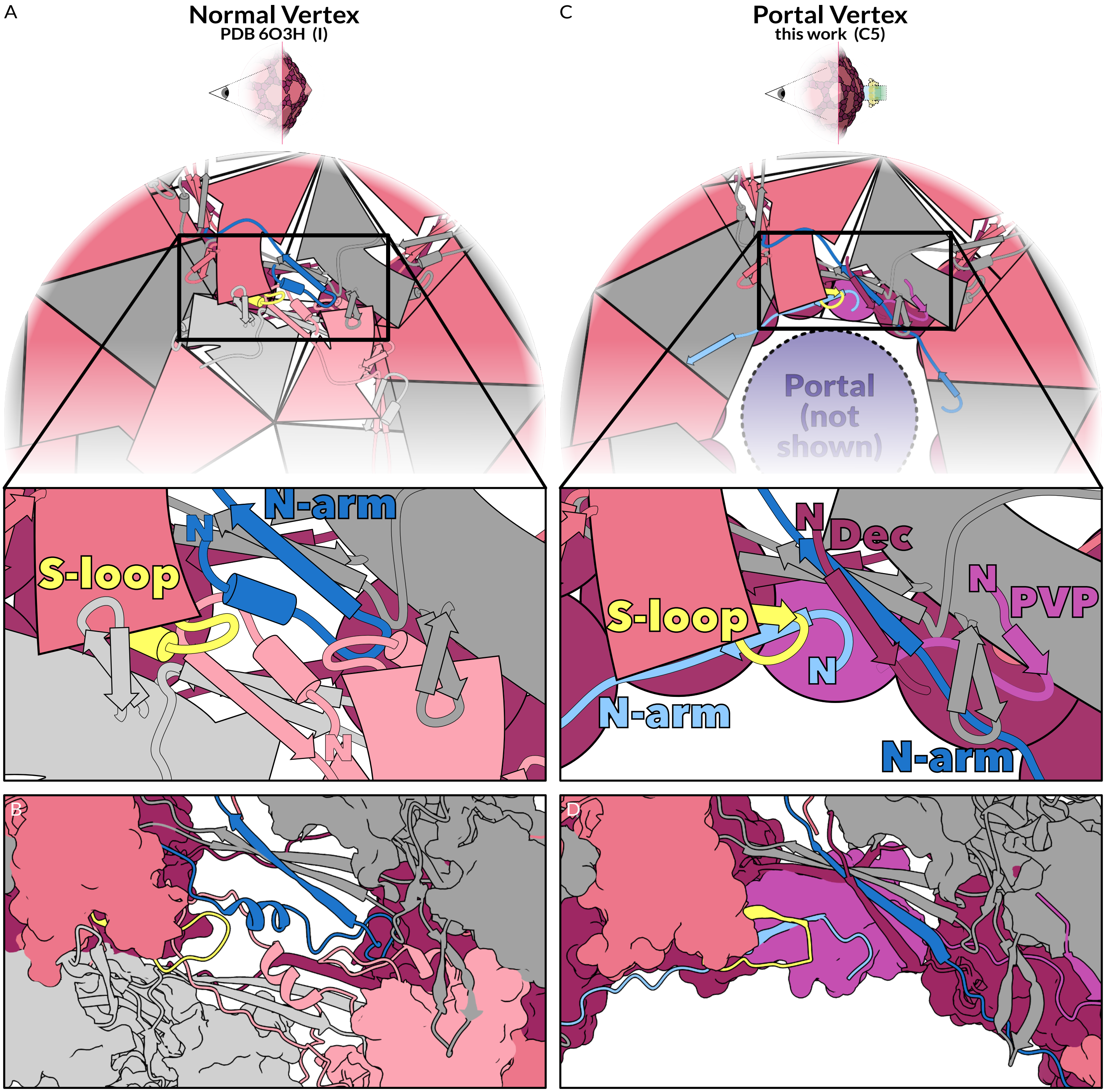


**Figure S25: The portal vertex components are highly interconnected. A,B. Interactions between a hexon MCP and the neigboring penton at a normal vertex (PDB 6O3H).** Hexon MCPs (pink and grey) and penton MCPs (light pink and light grey) interact through their S-loops and N-arms. The hexon MCP S-loop (yellow) and N-arm (blue) interact with the penton N-arm (light pink), while the N-terminus forms an intra-subunit interaction with its P-domain. Both the S-loop and N-arm have helical secondary structures. The N-arms of Dec (maroon) interact with the outer surface of the capsid. **C,D. Interactions between MCP, Dec, and PVP at the unique portal vertex (portal not shown).** Colors are as in A, except as noted. Rather than folding back to interact with the penton, the MCP N-arm (blue) extends into the hexamer to the right. The MCP N-arm from the hexamer to the left (light blue) interacts with the S-loop (yellow). While both the N-arm and S-loop form helices in the icosohedral MCP, they form β-strands that interact with each other in the unique vertex. The N-arms of one Dec monomer (maroon) and PVP (magenta) fold into the pore to interact with the inner surface of the capsid.

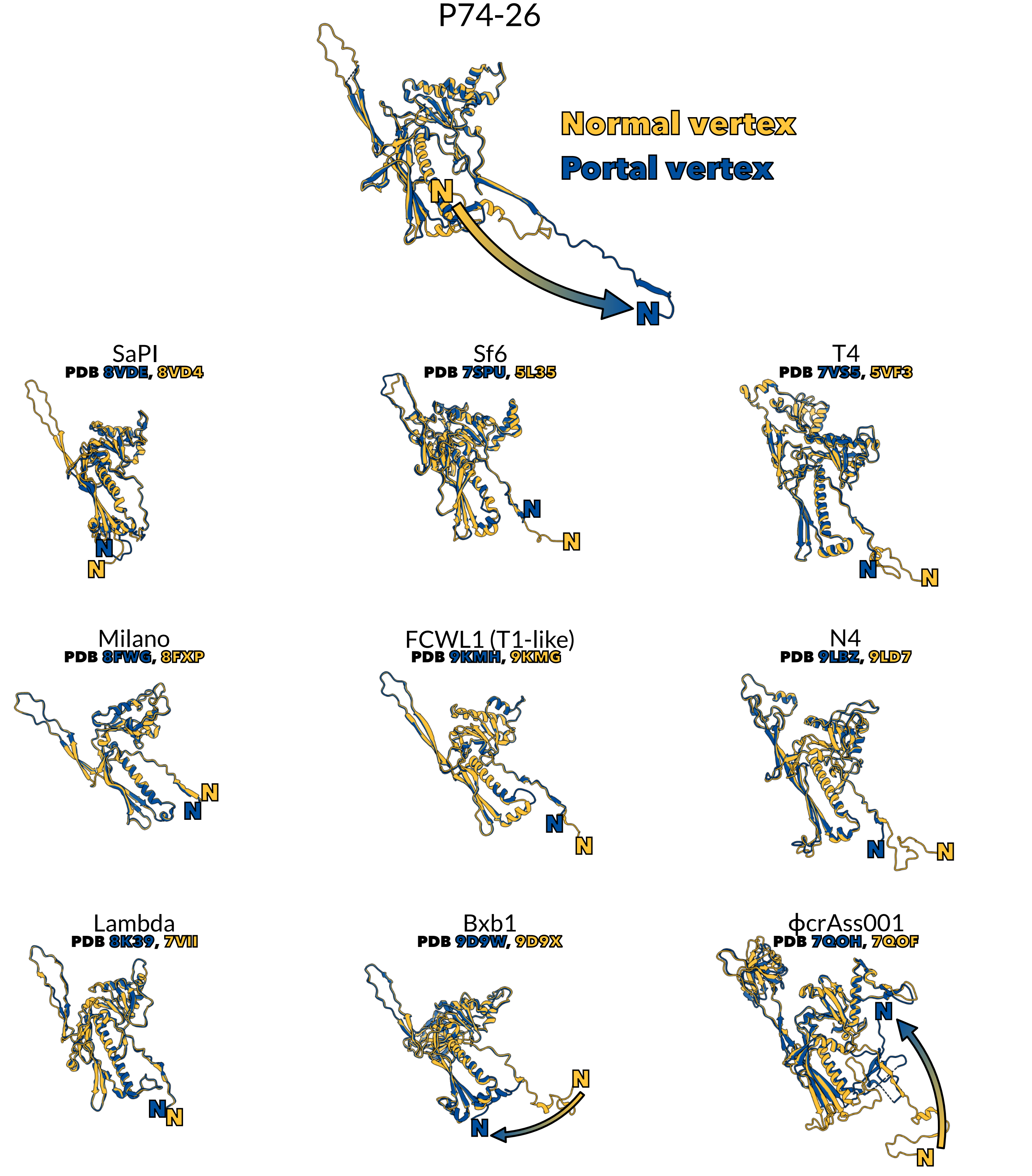


**Figure S26: MCPs from mesophilic phages do not intertwine at the portal vertex.** Top, a single MCP subunit from the junction of a hexamer with a pentamer at a normal vertex (yellow) overlaid with the corresponding MCP subunit from the portal vertex (blue). The N-termini are labeled and the arrow highlights the structural rearrangement made between the two conformations. The corresponding MCP subunits from 9 mesophilic phages were obtained from PDB and overlaid and labeled as above. Note that in some cases, the authors could not build the entire N-arm, likely because these regions are unstructured. In subunits where the normal N-arm contacts the adjacent pentamer, it either becomes unstructured (T4, FCWL1, N4) or rearranges to interact with its own hexamer (Bxb1, ϕcrAss001).

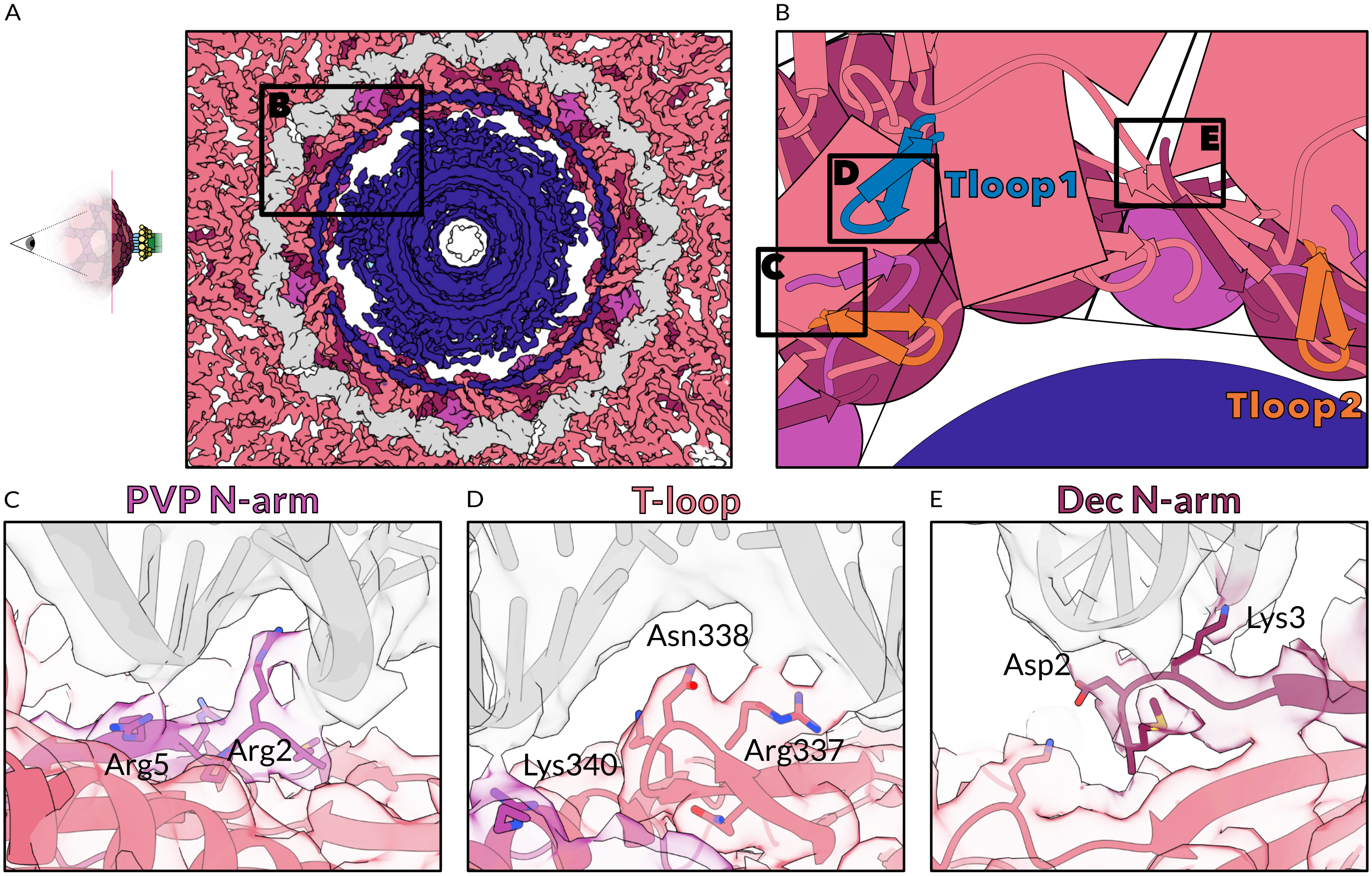


**Figure S27: One loop of packaged DNA surrounds portal with 5-fold symmetry. A. The C5 map, viewed from inside the capsid, colored by protein identity as in Figure 1 with DNA in grey.** The area enlarged and diagrammed in B is marked. **B. Diagram of DNA-contacting capsid regions (DNA not shown).** The areas enlarged in panels C-E are marked. Proteins are colored as in Figure 1. MCP, except for its T-loops, is shown in salmon. T-loop1, which contacts portal in the procapsid (Figure 4A), is shown in blue. T-loop2, which contacts portal in mature virions (Figure 4B), is shown in orange. **C. Interactions between the N-arm of PVP and DNA.** Arg2 inserts into the minor groove, while Arg5 interacts with the DNA backbone. **D. Interactions between the T-loop of MCP and DNA.** Arg337 interacts with the DNA backbone, while Asn338 inserts into the major groove. **E. Interactions between the N-arm of Dec and DNA.** Asp2 and Lys3 interact with the DNA backbone.

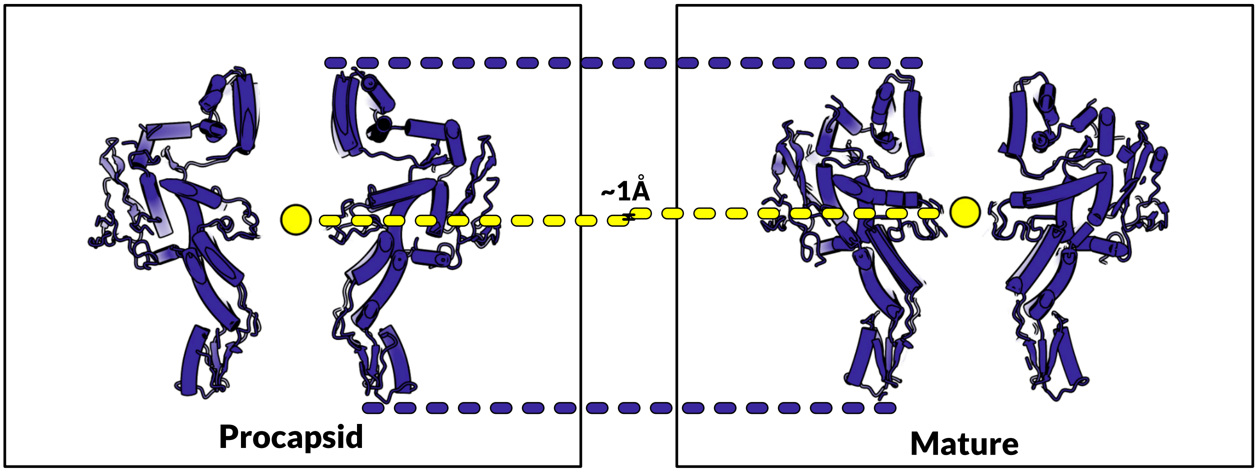


**Figure S28: The centroids of the procapsid portal and the mature portal are only 1Å apart.** *Left:* The procapsid portal (PDB 6QJT) with centroid (yellow sphere). *Right:* The mature virion portal with centroid (yellow sphere). The two models were aligned, the centroids determined, and the Δy between centroids calculated in ChimeraX.

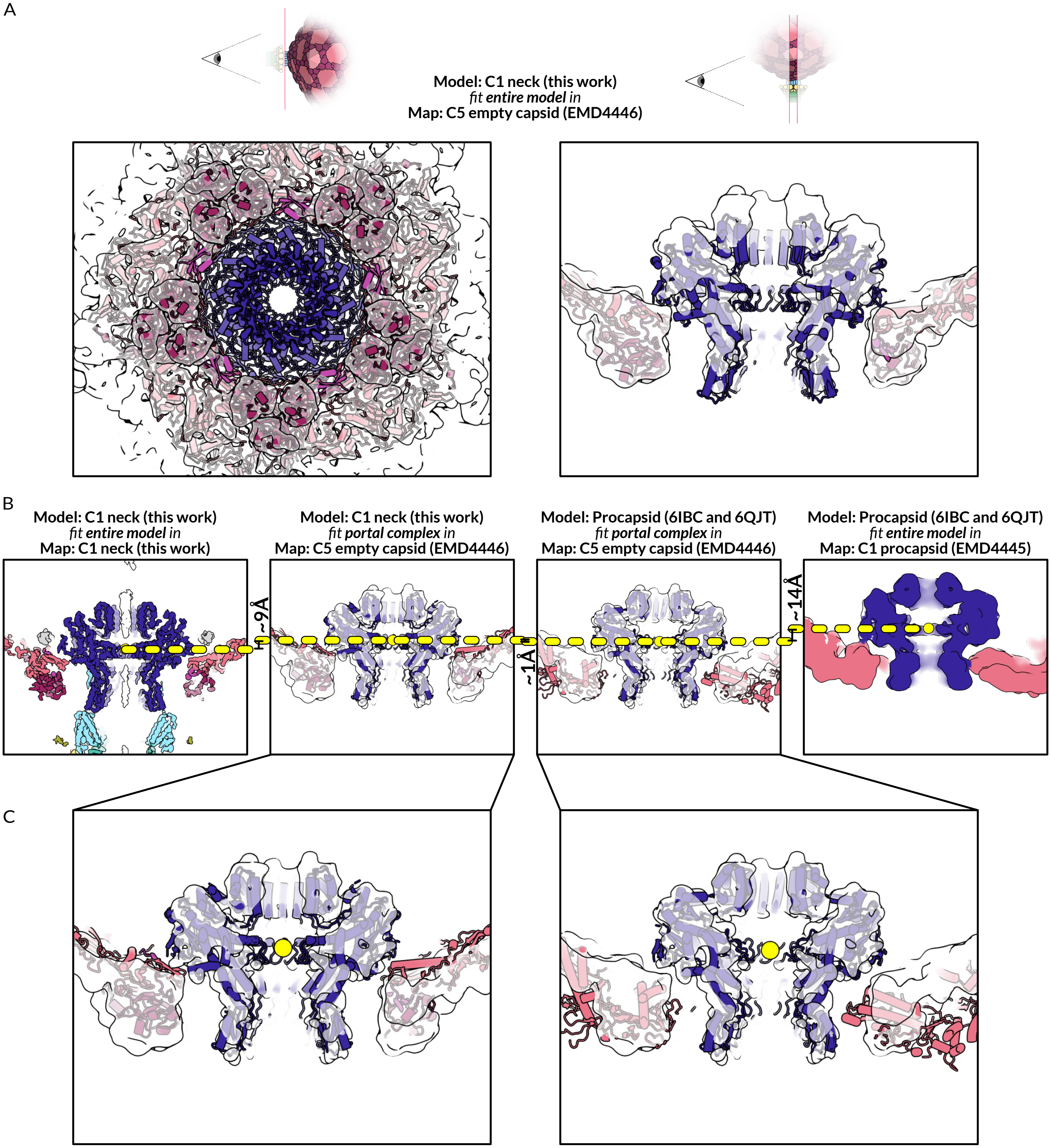


**Figure S29: The portal complex appears to relax in expanded empty capsids when the pressure of packaged DNA is no longer present. A. Two views of the portal and capsid of the mature neck model fit into the C5 expanded empty capsid map.** Note that the position of the mature portal model is significantly displaced from the portal density in empty capsids. **B. Comparison of the position of portal in mature virions (left), empty capsids (center right, center left) and procapsids (right).** When the mature portal complex is fit into the empty capsid density, its centroid is shifted 9Å higher than in mature virions. Note that the capsid proteins are also pulled out of their density. Likewise, when the procapsid portal is fit into the empty capsid density, its centroid is shifted 14Å lower than in procapsids. The centroids of the mature and procapsid portals are not significantly different when both are fit into the empty capsid map. **C. Enlargement of the mature and procapsid portals fit into the empty capsid map.** The portal complex in expanded empty capsids may be in a different conformation, or mixture of conformations, compared to mature virions and procapsids.

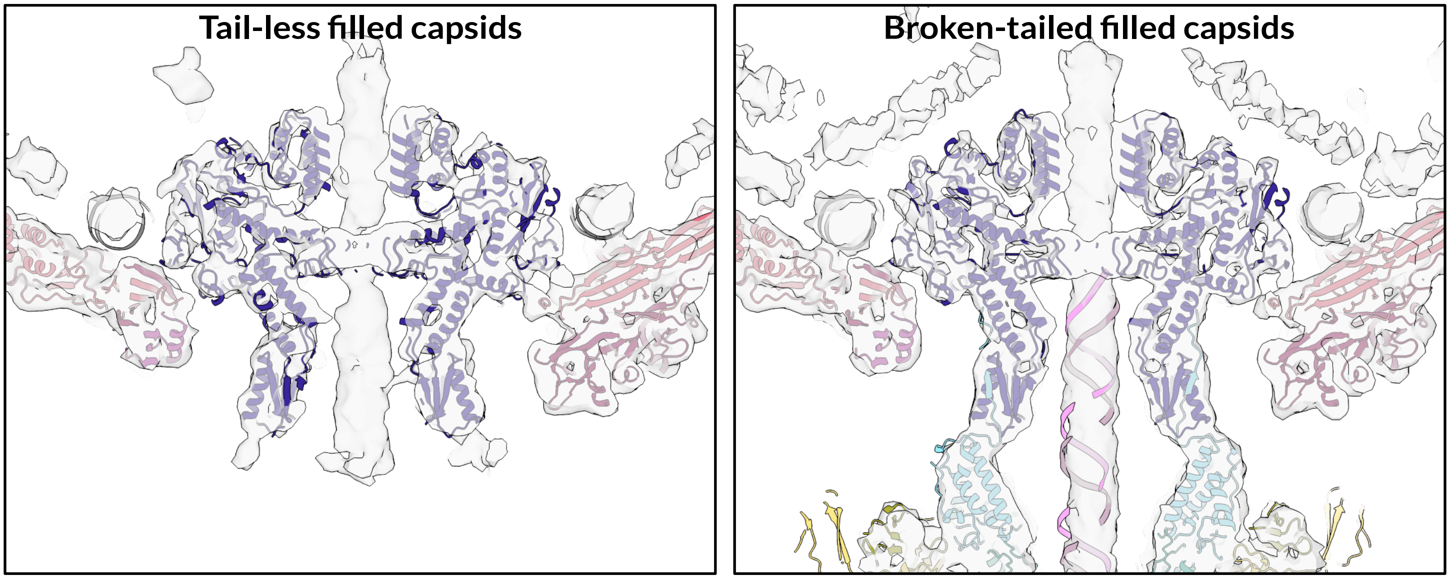


**Figure S30: The portal complex does not relax when the tail is removed.** Models for the mature portal and capsid complex (ribbon diagrams) fit into reconstructions of tail-less and broken-tailed phage from figures S13-14 (surfaces).

| Accession | Annotation | Species | Protein | Identity | Coverage | Map |
|---|---|---|---|---|---|---|
| YP_001468060.1 | hypothetical protein P74p90 | Thermus phage P74-26 | Adaptor | 89% | 96% | C3 |
| API81894.1 | hypothetical protein G20c_86 | G20C | Adaptor | 89% | 96% | C3 |
| YP_001467944.1 | hypothetical protein P23p91 | Thermus phage P23-45 | Adaptor | 88% | 96% | C3 |
| QAY18169.1 | hypothetical protein | Thermus phage TSP4 | Adaptor | 56% | 95% | C3 |
| YP_001468082.1 | hypothetical protein P74p112 | Thermus phage P74-26 | Collar | 81% | 75% | C3 |
| YP_001467966.1 | hypothetical protein P23p113 | Thermus phage P23-45 | Collar | 80% | 75% | C3 |
| 4ZJN_A | Portal protein | G20C | Portal | 82% | 98% | C3 |
| 5NGD_A | Portal protein | P74-26 | Portal | 82% | 97% | C3 |
| 6IBG_A | Portal protein | P23-45 | Portal | 81% | 99% | C3 |
| YP_001468055.1 | Portal protein | Thermus phage P74-26 | Portal | 81% | 99% | C3 |
| YP_001467939.1 | Portal protein | Thermus phage P23-45 | Portal | 81% | 99% | C3 |
| QAY18164.1 | Portal protein | Thermus phage TSP4 | Portal | 65% | 99% | C3 |
| YP_001468062.1 | hypothetical protein P74p92 | Thermus phage P74-26 | Terminator | 83% | 99% | C3 |
| YP_001467946.1 | hypothetical protein P23p93 | Thermus phage P23-45 | Terminator | 82% | 99% | C3 |
| API81896.1 | hypothetical protein G20c_88 | G20C | Terminator | 82% | 59% | C3 |
| QAY18171.1 | hypothetical protein | Thermus phage TSP4 | Terminator | 59% | 95% | C3 |
| YP_001468063.1 | hypothetical protein P74p93 | Thermus phage P74-26 | TTP | 71% | 73% | C3 |
| YP_001467947.1 | hypothetical protein P23p94 | Thermus phage P23-45 | TTP | 72% | 69% | C3 |
| API81897.1 | hypothetical protein G20c_89 | G20C | TTP | 69% | 69% | C3 |
| QAY18172.1 | hypothetical protein | Thermus phage TSP4 | TTP | 62% | 66% | C3 |
| 6BL5_A | Head decoration protein | Oshimavirus P7426 | Decoration Protein | 64% | 82% | C5 |
| YP_001468057.1 | hypothetical protein P74p87 | Thermus phage P74-26 | Decoration Protein | 64% | 74% | C5 |
| API81891.1 | hypothetical protein G20c_83 | Thermus phage G20c | Decoration Protein | 64% | 74% | C5 |
| YP_001467941.1 | hypothetical protein P23p88 | Thermus phage P23-45 | Decoration Protein | 54% | 68% | C5 |
| QAY18166.1 | hypothetical protein | Thermus phage TSP4 | Decoration Protein | 52% | 65% | C5 |
| YP_001468058.1 | major head protein | Thermus phage P74-26 | Major Capsid Protein | 54% | 68% | C5 |
| API81892.1 | major head protein | Thermus phage G20c | Major Capsid Protein | 52% | 60% | C5 |
| UYB98502.1 | hypothetical protein | Thermus phage P23-45 | Major Capsid Protein | 52% | 60% | C5 |
| YP_001467942.1 | major head protein | Thermus phage P23-45 | Major Capsid Protein | 52% | 60% | C5 |
| YP_001468004.1 | hypothetical protein P74p34 | Thermus phage P74-26 | Portal Vertex Protein | 61% | 97% | C5 |
| YP_001467889.1 | hypothetical protein P23p36 | Thermus phage P23-45 | Portal Vertex Protein | 61% | 97% | C5 |
| API81842.1 | hypothetical protein G20c_34 | Thermus phage G20c | Portal Vertex Protein | 60% | 97% | C5 |

**Table S1: ModelAngelo built peptides into the C3 and C5 maps that NCBI Blast aligned to both known and unknown P74-26 virion proteins, as well as homologs in closely related Thermus phages.** We used ModelAngelo to build peptides into the C3 and C5 maps without sequence input. We extracted the sequences of these peptides, filtered them to remove any sequences shorter than 10 amino acids, then used Blastp to search the Non-redundant protein sequence database. Blastp hits were filtered to exclude any hits with lower than 60% sequence identity. The Accession numbers for significant hits, along with their annotation in the NCBI Protein database and the source species are reported along with the name of each protein as reported and discussed in this work. Because of the high symmetry of the portal vertex and neck, multiple peptides aligned to each reference sequence. Thus, we report the average percent identity between all peptides and the reference sequence. We also report the percentage of the final manually-built model covered by the ModelAngelo model.

| | Accession | Annotation | MW (kDa) | Unique peptides | Total peptides | Stoichio metry |
|---|---|---|---|---|---|---|
| gp13 | A7XXH5 | Ribonucleoside-triphosphate reductase | 74.24 | 2 | 2 | ? |
| gp14 | A7XXH6 | Cytosine-specific methyltransferase | 40.48 | 3 | 4 | ? |
| gp26 | A7XXJ1 | dNMP kinase | 20.28 | 3 | 3 | ? |
| gp28 | A7XXJ4 | Uncharacterized protein | 9.32 | 2 | 4 | ? |
| gp29 | A7XXJ5 | Uncharacterized protein | 10.37 | 1 | 3 | ? |
| gp30 | A7XXJ6 | Deoxycytidine triphosphate deaminase | 19.91 | 4 | 10 | ? |
| **gp34** | **A7XXK1** | **Portal vertex protein (this work)** | **8.42** | **3** | **7** | **5** |
| gp35 | A7XXK3 | Uncharacterized protein | 35.36 | 1 | 2 | ? |
| gp41 | A7XXL0 | Uncharacterized protein | 11.46 | 3 | 5 | ? |
| gp43 | A7XXL2 | Uncharacterized protein | 21.16 | 1 | 1 | ? |
| gp49 | A7XXL9 | Chaperone/ATP-dependent lon protease | 29.09 | 2 | 2 | ? |
| gp63 | A7XXN7 | Uncharacterized protein | 11.57 | 2 | 2 | ? |
| gp84 | A7XXR1 | Terminase, large subunit | 55.1 | 14 | 47 | ? |
| **gp85** | **A7XXR3** | **Portal protein** | **49.64** | **26** | **273** | **12** |
| gp86 | A7XXR4 | Capsid scaffolding protein | 37.77 | 5 | 49 | ? |
| **gp87** | **A7XXR5** | **Decoration protein (Stone et al)** | **16.35** | **6** | **136** | **420** |
| **gp88** | **A7XXR6** | **Major capsid protein** | **46.6** | **41** | **2048** | **415** |
| **gp90** | **A7XXR8** | **Adaptor (this work)** | **14.53** | **8** | **78** | **12** |
| gp91 | A7XXS0 | Uncharacterized protein | 33.48 | 12 | 23 | ? |
| **gp92** | **A7XXS1** | **Tail terminator** | **17.55** | **3** | **23** | **6** |
| **gp93** | **A7XXS2** | **Tail tube protein (Agnello et al)** | **37.9** | **31** | **2202** | **?** |
| gp95 | A7XXS4 | Tape tail measure protein | 549.9 | 335 | 2414 | 3 (?) |
| gp96 | A7XXS6 | Uncharacterized protein | 21.69 | 3 | 15 | ? |
| gp97 | A7XXS7 | Lectin-like sugar binding domain | 25.21 | 17 | 42 | ? |
| gp98 | A7XXS8 | Uncharacterized protein | 40.64 | 18 | 90 | ? |
| gp99 | A7XXS9 | Uncharacterized protein | 128.98 | 41 | 249 | ? |
| gp100 | A7XXT0 | Uncharacterized protein | 57.11 | 21 | 225 | ? |
| gp101 | A7XXT2 | Uncharacterized protein | 13.51 | 4 | 14 | ? |
| gp102 | A7XXT3 | Uncharacterized protein | 61.55 | 20 | 99 | ? |
| gp104 | A7XXT5 | Uncharacterized protein | 33.24 | 11 | 178 | ? |
| gp106 | A7XXT8 | Uncharacterized protein | 14.38 | 2 | 4 | ? |
| gp107 | A7XXT9 | Peptidoglycan hydrolase | 17.13 | 2 | 3 | ? |
| gp108 | A7XXU0 | Holin | 12.62 | 7 | 41 | ? |
| gp109 | A7XXU1 | Metal-dependent hydrolase | 30.85 | 2 | 3 | ? |
| gp110 | A7XXU2 | 50S ribosomal protein L20 | 7.09 | 2 | 21 | ? |
| **gp112** | **A7XXU5** | **Collar protein (this work)** | **16.12** | **3** | **55** | **24** |
| gp113 | A7XXU6 | Uncharacterized protein | 19.58 | 6 | 18 | ? |
| gp114 | A7XXU7 | Uncharacterized protein | 29.93 | 1 | 2 | ? |

**Table S2: Mass Spectrometry of mature virions corroborates identification of neck and portal vertex proteins.** We used mass spectrometry to sequence the component peptides of mature virions. All sequenced peptides mapping to P74-26 proteins are reported here. The molecular weight of each protein, as well is its stoichiometry in mature virions (if known), is also provided for context. Proteins present in models deposited with this work are highlighted in yellow.

| | P74-26 | P22 | T4 |
|---|---|---|---|
| **Buried area (Å)**[1] | 9,275.8 | 28,050.2 | 16,246.5 |
| **$\Delta G^{int}$ (kcal/mol)**[2] | -0.7 | -120.7 | -66.8 |
| **$N_{HB}$**[3] | 57 | 116 | 68 |
| **$N_{SB}$**[4] | 38 | 24 | 37 |
| **PDB** | TBD | 8U10 | 6UZC |

[1] total solvent-accessible surface area that is buried in the interface between portal and capsid

[2] solvation free energy gain upon formation of the assembly, in kcal/M. The value is calculated as difference in total solvation energies of portal and capsid when isolated versus assembled . This value does not include the effect of satisfied hydrogen bonds and salt bridges.

[3] number of potential hydrogen bonds across the interface. Each hydrogen bond contributes about 0.5 kcal/mol into the free energy of protein binding

[4] number of potential salt bridges across the interface. Each salt bridge contributes about 0.3 kcal/mol into the free energy of protein binding

**Table S3: The interface between portal and capsid is less stable in P74-26 than in similar mesophilic phages.** We used the PDBePISA server to calcualte buried surface area, solvation free energy, and electrostatic interactions of the interface between portal and capsid. We calculated the energetics of both portal and capsid in isolation, then in complex, and report the difference between the complex and the sum of both component parts.

| | Neck | Portal Vertex | Portal and Portal Vertex | Central DNA |
|---|---|---|---|---|
| **EMDB accession** | TBD | TBD | TBD | TBD |
| **Number of Particles** | 30748 | 40773 | 39382 | 19306 |
| **Imposed Symmetry** | C3 | C5 | C1 | C1 |
| **Final Resolution (masked)** | 2.61 Å | 2.95 Å | 3.56 Å | 4.54 Å |
| **Final Resolution (unmasked)[1]** | 3.18 Å | 3.71 Å | 4.65 Å | 4.00 Å |
| **PDB accession** | TBD | TBD | TBD | TBD |
| **Map Correlation[2]** | 0.8927 | 0.8324 | 0.7982 | 0.7542 |
| **R.M.S.Z. (lengths)[1]** | 0.26 | 0.33 | 0.30 | 0.32 |
| **R.M.S.Z. (angles)[1]** | 0.49 | 0.58 | 0.54 | 0.77 |
| **All-atom Clash Score[2]** | 2.58 | 4.22 | 3.35 | 9.55 |
| **Ramachandran Favored[1]** | 98.76% | 97.73% | 98.06% | NA |
| **Ramachandran Allowed[1]** | 1.24% | 2.27% | 1.94% | NA |
| **Ramachandran Outliers[1]** | 0.00% | 0.00% | 0.00% | NA |
| **Rotameric Sidechains[1]** | 99.97% | 100.00% | 100.00% | NA |
| **Outlier Sidechains[1]** | 0.03% | 0.00% | 0.00% | NA |
| **C-beta deviations[2]** | 0.00% | 0.00% | 0.00% | NA |

[1]Reported by PDB validation server

[2]Reported by Phenix Real-Space Refine

**Table S4:** **EMDB accession numbers and resolutions of each map, with the PDB accession numbers of assoicated models and relevant measures of model quality.**

| | C12 neck | Collar-less | Capsid-less | Tail-less | Broken Tail |
|---|---|---|---|---|---|
| **EMDB accession** | TBD | TBD | TBD | TBD | TBD |
| **Number of Particles** | 31,671 | 8,007 | 24,712 | 3,964 | 13,186 |
| **Imposed Symmetry** | C12 | C12 | C3 | C1 | C1 |
| **Final Resolution (masked)** | 2.28 Å | 2.58 Å | 2.74 Å | 7.33 Å | 5.80 Å |
| **Final Resolution (unmasked)[1]** | 2.87 Å | 3.38 Å | 3.42 Å | 16.29 Å | 9.93 Å |

[1]Reported by PDB validation server

**Table S5:** **EMDB accession numbers and resolutions of each map that does not have an associated model.**
